# Development of a DNA-encoded library screening method “DEL Zipper” to empower the study of RNA-targeted chemical matter

**DOI:** 10.1101/2024.09.13.612806

**Authors:** Zhongyao Ma, Bin Zou, Jiannan Zhao, Rui Zhang, Qiaoqiao Zhu, Xiaofeng Wang, Linan Xu, Xiang Gao, Xinyue Hu, Wei Feng, Wen Luo, Wang Min, Yunyun He, Zhifeng Yu, Weiren Cui, Qi Zhang, Letian Kuai, Wenji Su

## Abstract

To date, RNA-targeted chemical matter is under explored due to a lack of robust screening assays. In this study, we present a novel RNA-targeted small molecule screening approach using a specialized DNA-encoded library (DEL). Our findings reveal that the specialized DEL library, called “DEL Zipper”, can significantly reduce single-stranded DNA-RNA region interaction signals during various kinds of RNA selection. By performing the selection against both G-quadruplex, we have identified novel hits that interact with RNA targets and the results are validated through binding. This study demonstrates that the “DEL Zipper” method is a robust screening assay that has potential for discovering small molecule ligands for diverse RNA targets.

## INTRODUCTION

Compared to conventional protein-targeted medicines, RNA-targeted therapeutics are a source of keen interest due to their unique physiological properties and novel mode-of-actions. RNA is now known to play crucial roles in physiological condition and disease progression through transcription and translation regulation, rather than only a genetic information carrier for protein translation^1^. First, every part of RNA, from untranslated regions (UTRs) to coding regions, can form diverse functional secondary and tertiary structures^2–4^ through base pairing and hydrogen bond formation^5^, which provide the foundations of how RNA may interact with wide range of small molecules. Furthermore, one recent report estimated that while ∼85% of the human genome is transcribed into RNA, only ∼3% of those transcripts are encoded for proteins^6^, indicating that the majority of RNAs are noncoding RNAs^7^. One class of noncoding RNAs, mircoRNAs, has already been validated as potential drug targets^8^, which suggests that RNA-targeted medicines have the potential to greatly expand the druggable human genome. Also, RNA-targeted therapeutics allows upstream modulation of protein expression, such as directly regulating mRNA translation ^9^ and splicing^10^, which can provide an alternative mode-of-action for modulating disease-relevant and otherwise undruggable protein targets. Comparing to gene therapy through genome editing at the DNA-level, which raise the concern of off-target gene editing, the effect of RNA-targeted therapeutics are transient and thus may provide superior long-term safety in the clinic^11^. For example, Risdiplam, approved recently by the FDA recently, directly interferes with the pre-mRNA splicing of survival of motor neuron 2 (SMN2), resulting in a rescue of SMN2 exon 7 exclusion which could substitute functionally for inactivation of survival of motor neuron 1 (SMN1) and hence as a treatment for spinal muscular atrophy (SMA)^12,13^.

In the past decade, finding novel chemical matter has been one of the bottlenecks of RNA-targeted therapeutics development, mostly due to the limitation in availability of screening assays. Many existing examples of RNA-targeted therapeutics were discovered via phenotypic screening rather than direct screening against purified RNA, which makes it challenging to identify a successful workflow for novel RNA targeted ligand discovery^6^. To date, affinity-screening-mass-spectrometry (ASMS)^14,15^ and fragment-based drug discovery (FBDD)^16,17^ are the most popular target-based screening approaches against purified RNA. However, the limited size of fragment libraries combined with historical high-throughput screening compound collections being geared towards protein as opposed to RNA interactions, limits the efficiency of finding novel ligand against RNA targets. Therefore, it is necessary to develop simple and high-throughput affinity-based screening methodologies which empowers the screening of novel RNA-targeting chemical matter to further accelerate RNA-targeted hit finding.

DNA-encoded library (DEL) technology has been recognized as one of the major screening methods with its unique advantages in vast chemical diversity and multiplexed affinity-based screening^18,19^. Though much interests have been shown in employing DEL as a major hit finding method for RNA targets, limited success has been reported. We believe a key challenge of applying DEL to RNA screening is the potential of DNA-RNA interaction. (Note that in most implementations of DEL screening, the DEL barcode includes a region of ssDNA with a randomized sequence. More details are provided in the Results section). An initial study that applies DEL to RNA targets was conducted by Benhamou et al., where the library molecules and DNA barcodes were built on a solid polymer bead. In the case of solid-phase DELs, the small molecule warhead is physically separated from the DNA barcode during the screening, and perhaps more importantly regarding potential interactions between the barcode and the RNA target, each bead is encoded with only a small amount of DNA barcode relative to screened chemical matter. The authors performed a DEL library screening against a RNA-encoded library and successfully identified selective ligands that inhibit pre-miR27a RNA processing, which is the oncogenic precursor of microRNA-27a^20^. Another recent study reported by Gibaut et al. utilized a similar solid-phase DEL to screen RNA ^21^, and successfully identified novel ligands against a r(CUG) repeat expansion using this methodology. More relevant to our work, employing a solution-phase DEL, Chen et al. reported a successful screening campaign against flavin mononucleotide (FMN) riboswitch by applying a RNA patch solution, in which the RNA patch is added to the DEL library before the selection, and blocks any potential interactions between the DNA barcode and the RNA target ^22^. This pre-blocking treatment utilizes structural information on the RNA target and a case-by-case treatment of DEL library for each selection. Thus, it’s necessary to further develop DEL-based selection methods against RNA target, which should be widely applied to the most Implemented DEL library format as well as require minimum case-by-case treatment and pre-exist structure knowledge, so that DEL selection could be easily applied to those RNA targets which hasn’t been previously explored.

In this paper, we demonstrate an optimized DEL screening strategy against RNA over the current practice. We show that the single stranded random region of the DNA barcode in a DEL molecule can be the major source of non-specific binding if not treated properly. Instead of blocking the DNA barcode with exogenous RNA prior to selection, a new approach (i.e. convert the ssDNA region of the barcode into dsDNA) was adopted to seal the region and successfully reduced the non-specific binding. We have named this method the “DEL Zipper”. With this method, we ensure the robustness of the screen and therefore alleviate the concern of screening RNA using DEL. Another advantage of this approach is it allows pre-treatment of the library in bulk which to reduce labour and should increase reproducibility in the screening outcome.

With the new “DEL Zipper” strategy, successful screening campaigns against G-quadruplex (G4) RNA within the hepatitis C virus (HCV) core were conducted. In the G-quadruplex (G4) RNA screening, DEL Zipper Library shows more post-selection signals and patterns compared to conventional DEL library. The binding of the novel ligand against G4 RNA was first demonstrated by affinity-based mass-spectrometry and the binding affinity was further determined by surface plasmon resonance (SPR). These findings demonstrate the feasibility of DEL selection on various RNA targets and this study could pave the way of greater success in the field for RNA-targeting therapeutics.

## RESULTS

Specialized “DEL Zipper” library decreases ssDNA-RNA interaction signals in THF-riboswitch RNA selection The most widely applied method for DNA-encoded libraries (DEL) synthesis is a combinatorial split and pool approach^23^, in which individual steps of chemical synthesis are recoded by corresponding DNA tag ligations. DEL libraries utilized in our study consist of two different regions of DNA tags: a double-stranded DNA (dsDNA) tag region that contains the coding sequence for the corresponding chemical building block (BB) and a single-stranded DNA (ssDNA) tag region that mainly includes the pool ID and unique molecule identifier (UMI), which is important for reducing PCR and sequencing biases^24^ (Fig. 1A). (Note that the UMI region of each library member is a random sequence, and thus for practical reasons cannot be readily incorporated into the barcode as dsDNA during library synthesis). Previous studies have demonstrated that both dsDNA and ssDNA region can bind to RNA^22^, in which for the ssDNA tag region, an oligonucleotide could binds to RNA through complementary Watson-Crick base pairing^25^, while for the dsDNA tag region, oligonucleotide triplex formation may occur through Hoogsteen base-pairing, which involves specific hydrogen-bonding patterns^26^.

**Fig. 1.**
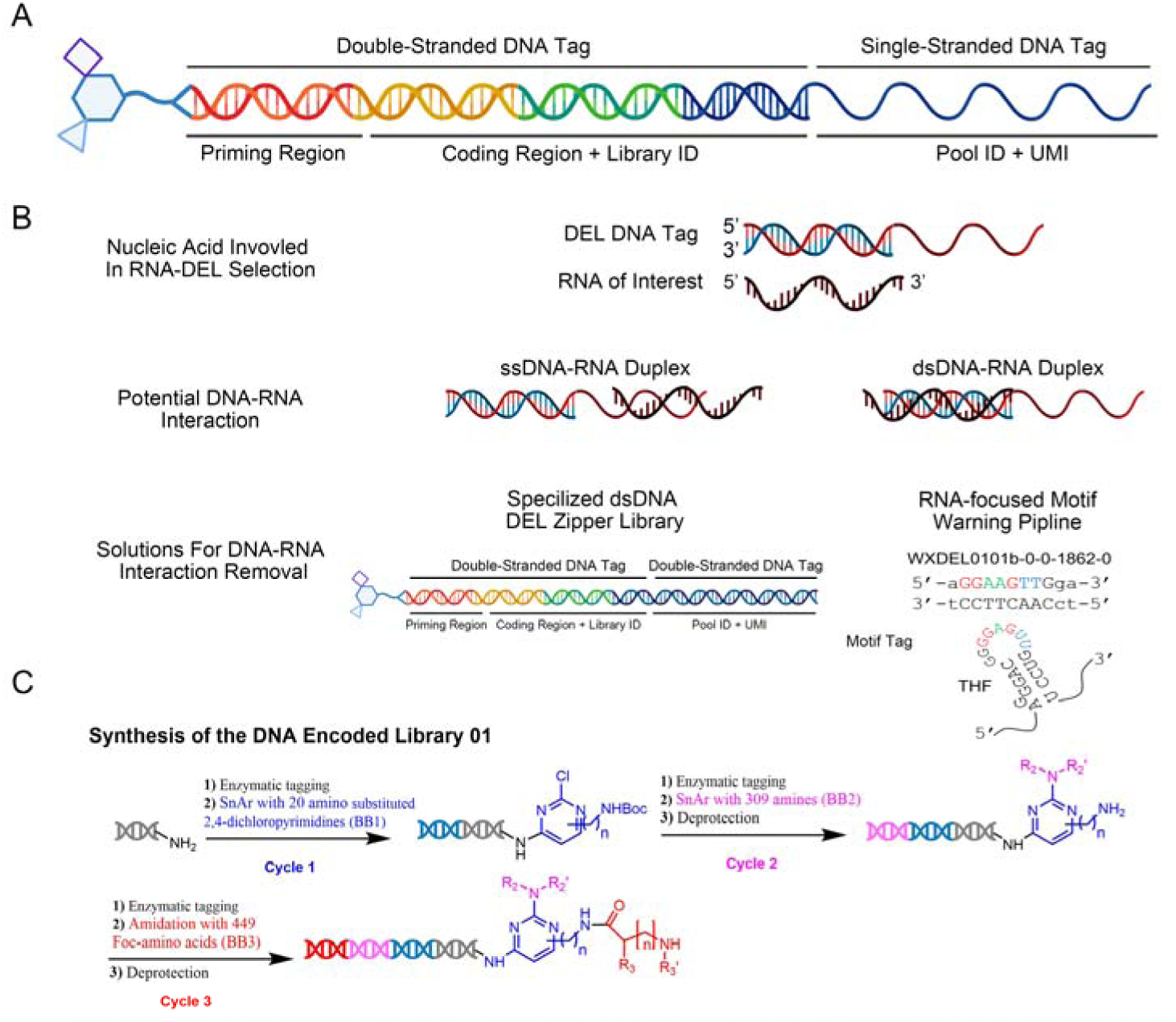
Schematic diagram of DNA-RNA interaction solutions in DEL selection. (A) General representation of conventional DNA encoded library (DEL). In general, our DNA barcode of DNA encoded library consist of two major parts, the double stranded DNA tag region and the single stranded DNA tag region. Priming region, coding region and library ID region are located in the double stranded DNA region while Pool ID and unique molecular identifier (UMI) are located in the single stranded DNA tag region. (B) Schematic diagram shows the potential DNA-RNA interaction during DEL selection and our solutions for DNA-RNA interactions. Basically, for ssDNA-RNA interaction, specialized dsDNA “DEL Zipper” library is used to remove the potential interaction signals. For dsDNA-RNA interaction, an in-house RNA-focused motif warning pipeline are developed to remove any potential motif in the downstream data analysis (see also supplementary Figure 1 for more details). One motif detected in THF-RNA selection is shown as an example. (C) Detailed synthesis route of DNA encoded library01.

To develop a novel DNA-encoded library screening method to empower the study of RNA-targeted chemical matter, corresponding noise-reduction method was developed to solve both dsDNA & ssDNA inference in RNA selection (Fig. 1B). In brief, our DEL library was converted into a specialized blunt-end dsDEL library called “DEL Zipper” to avoid the ssDNA-ssRNA interaction-derived noise (Fig. 1B). Additionally, an RNA focused bioinformatics pipeline was involved for detecting codon region motifs so that we could remove all the potential dsDNA-ssRNA interaction derived motifs from downstream analysis. (Fig. 1B & Supplementary Fig. 1). An example synthesis route of DNA encoded library and all library chemical formulas used in this study for RNA ligand validation are provided as an example (Fig. 1C). The detailed “DEL Zipper” library construction process and RNA focused motif analysis pipeline is described in the experimental section.

To evaluate the performance and feasibility of this “DEL Zipper” based selection method, a selection against a highly structured RNA was performed. Riboswitches, which are structured mRNA segments residing in 5’ untranslated regions (UTRs), play a crucial role in bacteria’s biological process^27^. Tetrahydrofolate (THF) riboswitch controls the expression of folate synthesis by conformation change of the sensing domain. Since the THF riboswitch has a function-related structure including both stem, apical loop & hairpin region^28^, it serves as a good candidate for testing the DNA-RNA interaction (or lack thereof) in our specialized “DEL Zipper” library pipeline. First, we synthesized the barcoded tool compound 5-formyl-tetrahydrofolate (folinic acid) employing historical procedures and then zippered the barcode using the methods described were produced (Fig. 2A). The zipped versus un-zipped tool molecules showed a clear size difference in gel electrophoresis, and the zipped molecule appears to possess high purity (Fig. 2B). Since RNA needs to be immobilized to a solid matrix during the DEL selection, biotinylated THF-riboswitch was acquired for testing. The structure of biotinylated THF-riboswitch was validated using folinic acid as tool compound, and a binding affinity of 1.9 µM was observed by SPR, which is comparable to previous report^29^ (Fig. 2C). Immobilization of RNA under DEL selection conditions was confirmed by comparing capture of biotinylated to non-biotinylated THF riboswitch. Only biotinylated THF-riboswitch remained on beads after immobilization, indicating a high efficiency of immobilization and minimal nonspecific interactions between RNA itself and matrix (Fig. 2D). We further tested THF riboswitch activity after immobilization by pre-selection. Using the zipped tool compound, immobilized THF riboswitch showed good enrichment signals as well as recovery rate compared to the blank beads control group as well as the zipped no-compound spacer-headpiece (SHP) control (Fig. 2E and Fig. 2F). Strikingly, when we performed the pre-selection experiment using unzipped spacer-headpiece (SHP) control molecule, the final molecule copies and recovery rate were significantly different between RNA group and NTC group. These results demonstrated that unzipped DNA barcode could interact with RNA under normal selection conditions (Supplementary Fig. 2).

**Fig. 2.**
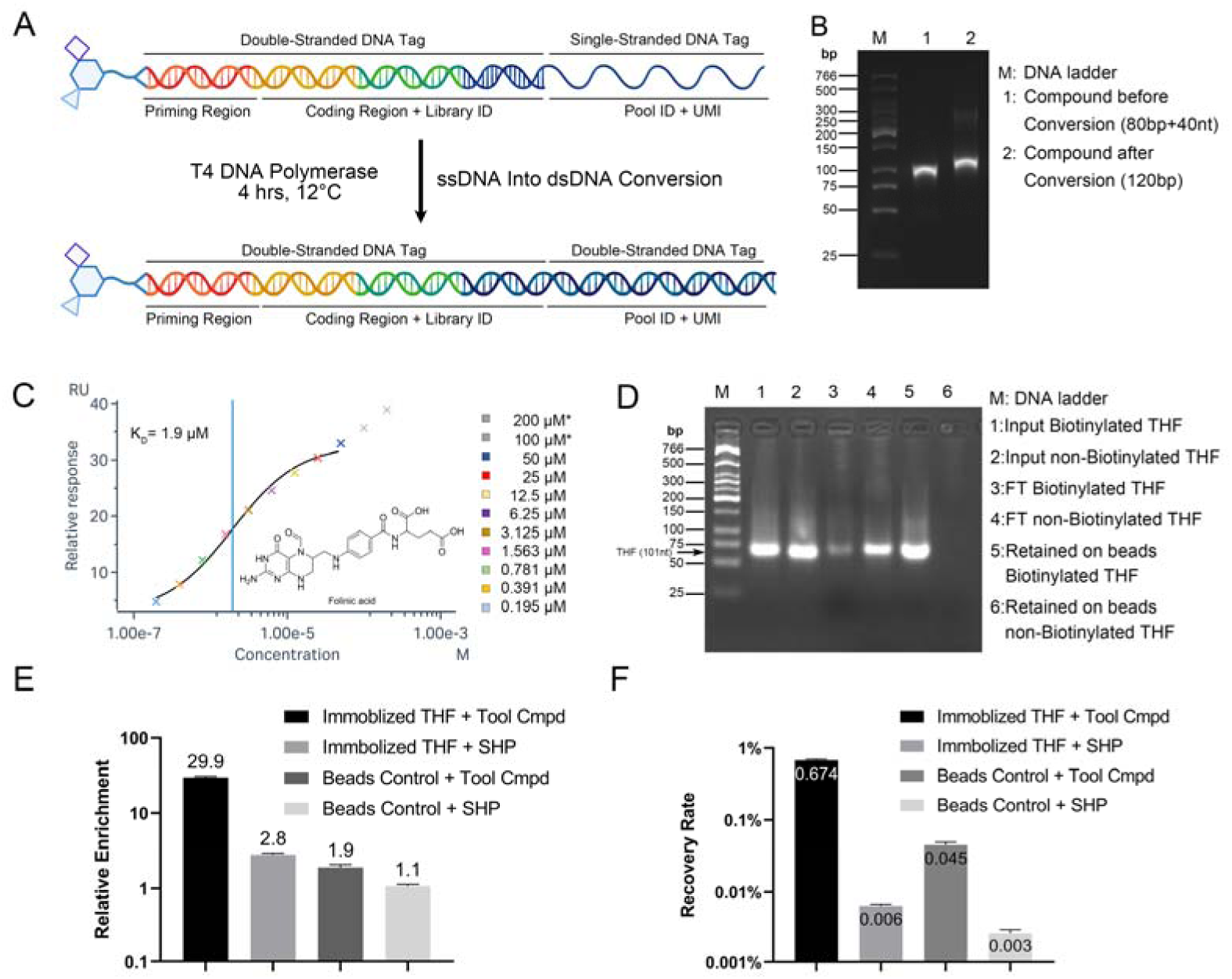
“DEL Zipper” library construction & THF riboswitch RNA pre-selection validation. (A) Schematic diagram shows the conversion from ssDNA conventional DEL library into “DEL Zipper” library. In general, single-stranded region of DEL DNA was blunted using T4 DNA polymerase kit (NEB M0203L) in 150 μl 1X NEBuffer r2.1 supplemented with 100 μM of each dNTP. 1unit T4 DNA polymerase per microgram DEL DNA was added. The reaction system was incubated 4hrs at 12°C then stopped by heating to 75°C for 20 minutes (B) Gel electrophoresis shows that a clear size difference between conventional compound molecules and after conversion compound molecules (C) SPR demonstrated that biotinylated THF riboswitch is still active and maintains binding ability (K_D_=1.9µM) to tool compound (Folinic acid) after immobilization. 100µM & 200µM are removed from curve fitting due to the second binding event. (D) Gel electrophoresis of beads capture assay shows that biotinylated THF riboswitch RNA could be successfully immobilized to beads under DEL selection condition. Majority of the biotinylated THF RNA was retained on beads while a small amount was observed in the flow through (FT). No immobilization is observed in non-biotinylated THF riboswitch RNA. (E to F) THF riboswitch RNA retains its binding activity to tool compound (5-forml-THF) after immobilized to beads. Enrichment (E) and recovery rate (F) are significantly higher (15.9-fold change in enrichment and 15.2-fold change in recovery rate) compare to blank beads. Data are the mean ± SEM of 3 replicates (n = 3).

Since biotinylated THF-riboswitch showed good immobilization efficiency and preserved tool compound binding ability, we proceeded with a DEL selection to compare conventional unzippered DEL versus “DEL Zipper” performance. “DEL Zipper” library showed a clear size increase as well as acceptable product purity after conversion, and thus was deemed ready for selection (Supplementary Fig. 3). After affinity selection and high-throughput sequencing, data was processed using the in-house pipeline (Supplementary Fig. 1). Briefly, raw FASTQ files were trimmed. Then sample identifiers, library tags and BB tags were decoded in order based on tag-to-BuildingBlock (BB) code table. Enrichment of Nsynthons were calculated^30^. Similar pipeline has been used in DEL decoder software package. DEL decoder is available upon request. Theoretically, the unique molecule identifier (UMI) sequence is randomly distributed in all library molecules and should have no correlation to compound structure (recall that the UMI is comprised of ssDNA in the conventional unzipped barcode). Thus, an even distribution of UMI sequence should be observed post-selection regardless of the selection conditions. In the No Target Control (NTC) group, both conventional DEL & “DEL Zipper” showed an even distribution of UMI sequence as expected (Fig. 3A). However, over 10x average enrichment of specific UMI sequences in THF-riboswitch selection was observed for the conventional DEL group (Fig. 3A). These results indicate a strong ssDNA-RNA interaction may exist when conventional DEL was used in RNA selections. To further investigate this potential DNA-RNA interaction-derived UMI motif, a copy of top 10000 UMI motifs versus the total useful sequence was analyzed. Surprisingly, these potential UMI motifs comprise 40% of total useful data in RNA selection when using conventional DEL. In comparison, “DEL Zipper” and NTC groups have less than 0.4% data that is potentially sequence motif driven, 100-fold lower than for the conventional DEL group (Fig. 3B). Potential motif sequences were aligned against the THF riboswitch reference sequence, and a large number of reverse complementary pairing in the position of 35-55nt in the RNA target were found only in conventional DEL group, while in “DEL Zipper” group, this RNA sequence-specific signal was almost undetectable (Fig. 3C). For example, UMI motif alignment average counts within the nucleotide 36 to 48 region in THF riboswitch RNA using the “DEL Zipper” selection was 130.3-fold lower compared to the conventional DEL selection (Conventional DEL: 8.768E+05 copies vs “DEL Zipper”: 6.729E+03 copies) (Fig. 3C). Together, these results indicated that a potential ssDNA-RNA interaction was present when using conventional DEL. This DNA-RNA interaction takes up the majority of post-selection data and could cause significant data loss and potential of RNA structure interference. By using “DEL Zipper”, such ssDNA-RNA interaction events were almost completely removed.

**Fig. 3.**
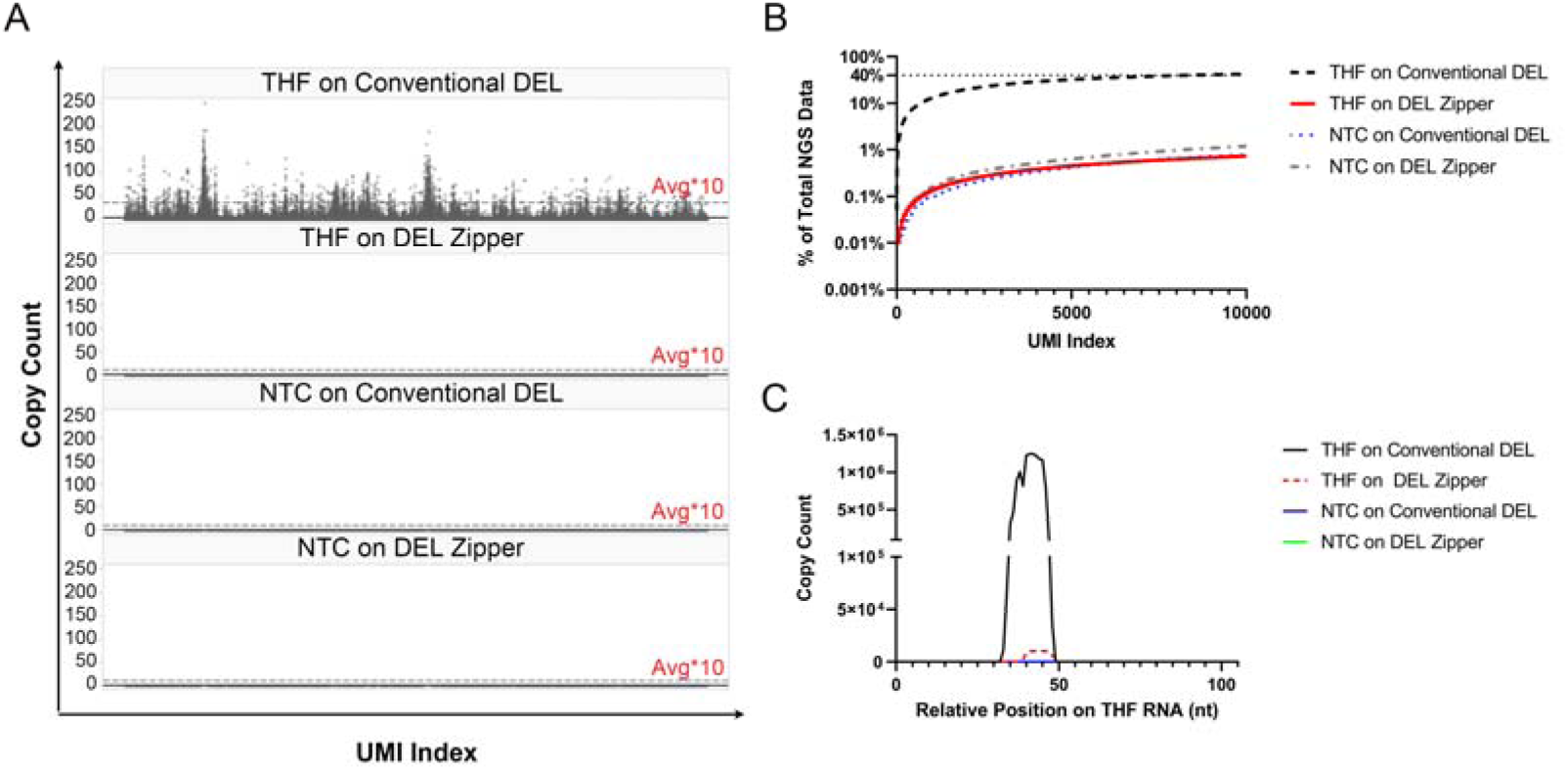
DNA binding events comparison between conventional DEL library and “DEL Zipper” library within THF riboswitch RNA selection. (A) Overall unique molecule identifier (UMI) copy count distribution of warhead-less library 10341 in THF riboswitch RNA DEL selection using conventional DEL library and “DEL Zipper” library. Significant UMI copy count bias is detected only in THF riboswitch RNA selection using conventional DEL group, in which vast UMI are 10x above the average copy count cutoff. (B) Copy accumulation curve of potential UMI motif and overall proportion of total NGS data. In THF on conventional DEL groups around 40% of total NGS data is DNA binding events related. (C) Copy count of potential UMI motif following alignment of reference RNA of THF riboswitch. Majority of the UMI motifs are restricted to a specific region of THF riboswitch RNA. UMI motif alignment average counts between nucleotide 36 to 48 in THF riboswitch RNA in “DEL Zipper” selection is 130.3-fold lower than conventional DEL selection (Conventional DEL: 8.768E+05 vs “DEL Zipper”: 6.729E+03).

As described above, an RNA-specific Motif Warning pipeline for dsDNA-RNA interaction was also performed during data analysis. For THF-riboswitch, only a small proportion (0 motif after 2 rounds selection (R2) and 6 motifs after 3 rounds of selection (R3)) of barcodes were affected by dsDNA-RNA interaction. The major motif sequence ‘AGGAAGTTGGA’ was detected and removed from the downstream analysis (Supplementary Table 1).

Small molecule binder against HCV G-quadruplex (G4) identified by “DEL Zipper” selection Since the “DEL Zipper” plus bioinformatics motif warning proved to be sufficient in removing unwanted DNA-RNA interactions in the DEL selection against THF riboswitch, this optimized “DEL Zipper” selection system was next tested on a short but structured virus RNA, hepatitis C virus (HCV) G-quadruplex (G4)-RNA. G4-RNA’s structure consists of stacked planar G-quartets, mainly located on 5’ or 3’ untranslated regions (UTRs) of mRNA^31^. In the HCV, the G4 sequence represents a potential drug target since ligand binding, such as pyridostatin derivative (PDP), can significantly inhibit HCV RNA replication. Thus, HCV-G4-RNA and its mutant version as previously reported ^32^ were chosen as our DEL selection targets, and pyridostatin (PDS) and PDP as tool compounds (Fig. 4A). The biotinylatedG4-RNA’s activity was confirmed by microscale thermophoresis (MST) in which tool compound PDS binding affinity was determined as K_D_ ∼2.9±1.0 µM (Fig. 4B). The activity of immobilized wild-type G4-RNA was confirmed by an on-DNA tool compound PDP binding test. In this test, only immobilized G4-RNA plus PDP group showed significant enrichment, which was greater than 33-fold over the control groups (Fig. 4C). We then proceed to the DEL selection employing the “DEL Zipper”, a further analysis of the enriched compounds was performed and visualized in a cubic scatter plot (Fig. 4D). In the cube, each axis represents the unique ID of building blocks used in each corresponding chemistry cycle, and the size of spheres is proportional to the copy count of each chemical structure. The structure patterns observed in the cube can be identified by cluster analysis of building blocks in each cycle and yields the structural features of potential G4-RNA hit molecules. After filtering out signals below 5-folds enrichment, an obvious plane with molecules containing the same cycle-1 synthon was observed. 2-amine substituted 6,7-dihydro-5H-pyrrolo [3,4-d] pyrimidine (a combination of cycle-1 and cycle-2 synthons) was identified as the key scaffold for molecules in this enriched plane. There appears to be no preference for which cycle-3 synthon is connected to this scaffold by amide. No potential ssDNA-RNA interaction was identified based on motif analysis for “DEL Zipper” library (Supplementary Fig. 4A). Moreover, the “DEL Zipper” selection shows more potential patterns and higher enrichment compared to the conventional DEL library (Supplementary Fig. 5A & B).

**Fig. 4.**
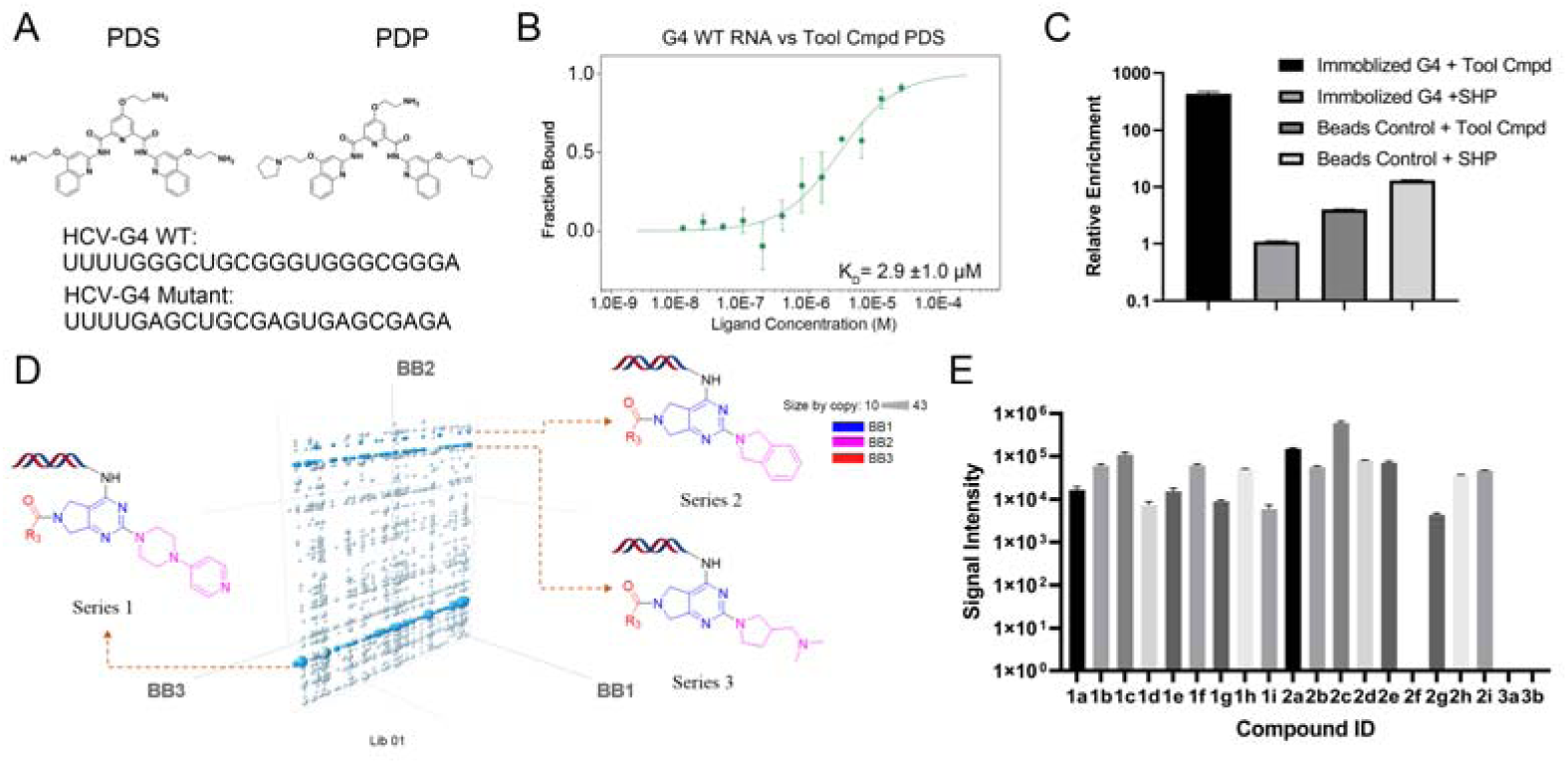
HCV G-quadruplex (G4) RNA pre-selection validation, enriched structural features in DEL selection using “DEL Zipper” library and primary hits validation using ASMS. (A) Sequence of HCV-G4 wildtype and HCV-G4 mutant and tool compound PDP, PDS structure used in pre-selection. (B) Microscale thermophoresis (MST) shows that biotinylated G4-RNA retains its binding capability against tool compound PDS. (C) HCV-G4 wildtype RNA retains binding activity against on-DNA tool compound PDP after immobilization to beads under selection condition. Data are the mean ± SEM of 3 replicates (n = 3). (D) Cubic plot of libraries with preferred scaffolds and the corresponding structural features. For series 1, the BB1 and BB2 showed clear structural features with various BB3. For series 2, the BB1 and BB2 showed enriched structural features with various BB3. (E) Primary validation of DEL selected hits against HCV-G4 wildtype RNA. 17 of 20 total selected compounds are validated in ASMS. Data are the mean ± SEM of 3 replicates (n = 3).

To validate potential hits, three series of off-DNA compounds were synthesized (see supplementary Table 2 for compound structures). As judged by ASMS, confirmation of compound binding against wildtype G4-RNA could be determined for compounds belonging to both series-1 and series-2. In contrast, no binding to the mutant G4-RNA mutant was detected (Fig. 4E, Supplementary Fig. 4B&4C and Supplementary Table 3) Two molecules from series 1 (1f & 1d) and another two molecules from series 2 (2h & 2d) that were confirmed binders by ASMS were further investigated by surface plasmon resonance (SPR) (Fig. 5). The binding of these four compounds against target wildtype G4-RNA were validated in the SPR assay and relative K_D_ value were determined. In contrast, no binding of SPR for these four compounds was observed for the mutant G4-RNA. Together, this selection, hit follow-up and compound profiling provide a proof-of-concept that the “DEL Zipper” RNA selection can be used to identify hit compounds against structured RNA such as HCV virus G4-RNA with no significant interference from DNA-RNA interactions.

**Figure 5.**
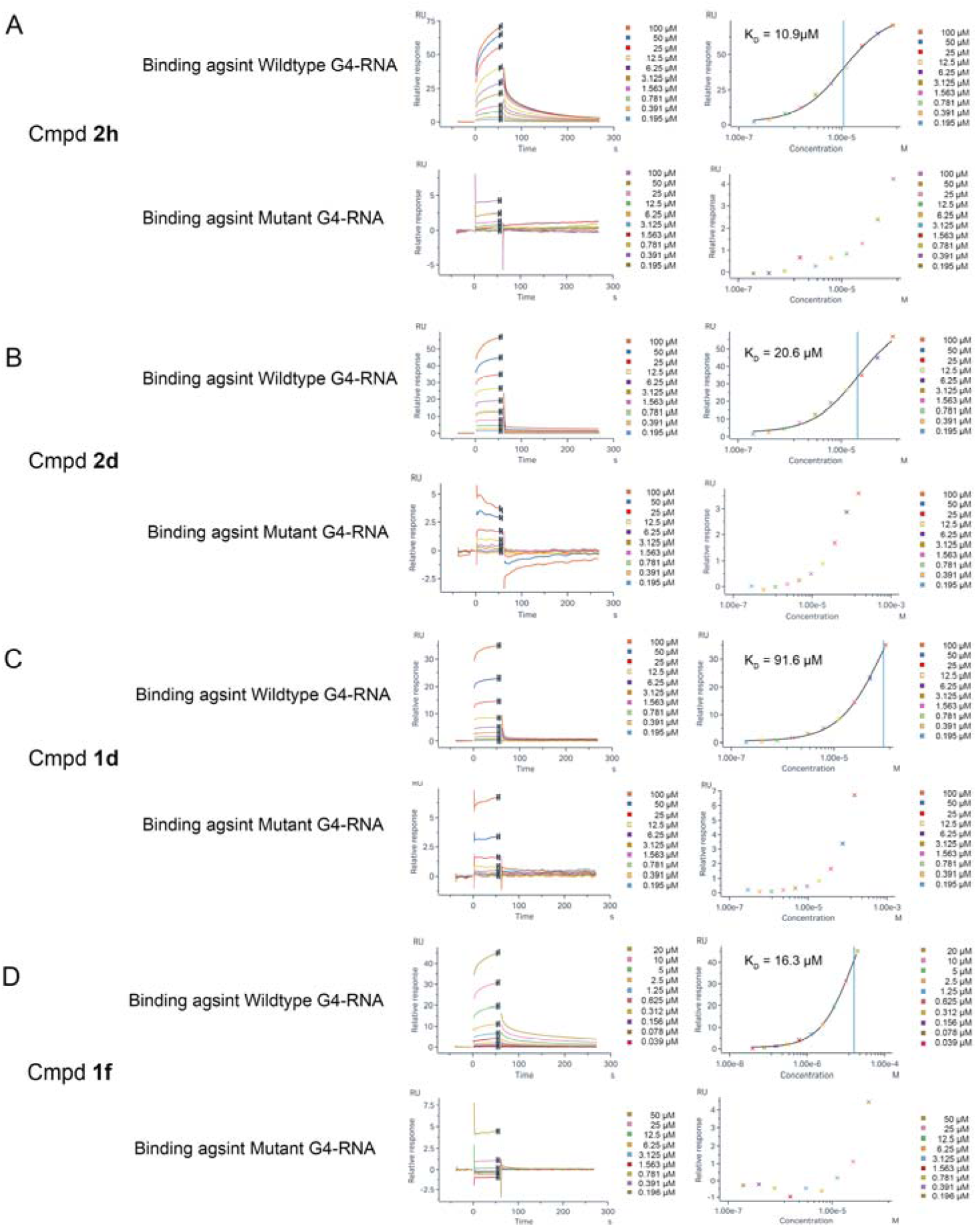
Surface plasmon resonance (SPR) results of DEL selected G4 compounds 2h, 2d, 1d and 1f. In SPR experiments, immobilized G4 wildtype RNA and immobilized G4 mutant RNA was tested against various concentration of DEL selected compounds **2h, 2d, 1d** (0.195 µM to 100 µM) and DEL selected compounds **1f** (0.039 µM to 20 µM in G4 Wildtype RNA and 0.196 µM to 50 µM in G4 mutant RNA due to compound low solubility). Corresponding sensogram response, kinetic curve was recorded and K_d_ were determined using steady state fit for each compound in wildtype G4-RNA. Relative signal Response of all 4 compounds is higher in wildtype G4-RNA comparing to mutant G4-RNA. K_d_ were not able to be determined in mutant G4-RNA due to low signal response.

## DISCUSSION

DNA Encoded Library (DEL) technology has been widely adopted for finding chemical starting points in early drug discovery. However, only limited cases have been reported on applying DEL to nucleic acid targets^20–22,33^, which we believe is mainly due to unwanted interactions between the DNA barcode and target RNA, difficulties in applying to coding mRNA targets and the complexity of making DELs compatible with RNA screening. Herein, we developed a specialized platform for RNA-targeted DEL screening, including using a blunt-end double-stranded “DEL Zipper” library and an RNA-focused motif analysis tool.

DNA-RNA interaction is one of the major concerns that prevents applying DEL technology to RNA-targeted drug discovery. In this study, we showed that “DEL Zipper” library can be easily prepared and screened, which can almost perfectly remove the potential DNA-RNA interaction that could generate false positive signals. As a testament to the success of this approach, multiple RNA targets with different levels of structural complexity have been screened. In the screening result, ssDNA-RNA non-specific binding event was almost undetectable. On the other hand, RNA-focused motif analysis tools helped to report all potential dsDNA-RNA interaction events so that these signals can be removed to avoid causing bias in the downstream compound selection process. Notably, the simplicity of library production in advance and the availability of effective motif analysis workflow make this platform amenable to a wide variety of RNA targets, in contrast to case-by-case blocking in previously reported RNA selection methods, where preliminary RNA structure information may also be necessary^22^. Another benefit of our method is that it improves the signal-to-noise of the screening result, which is observed in G4-RNA selection.

In this study, we applied the “DEL Zipper” based selection methodology to HCV-G4 RNA, since similar structure in the DNA formed G4 has been successfully screened before using DEL^33^. Several hits that bind to HCV-G4 were identified and validated by ASMS and SPR. Notably, these HCV-G4 hits showed no binding signals against G4-mutant sequence, indicating the inclusion of a proper counter-target may be important in finding selective hits.

In summary, our study presented a robust DEL screening methodology that could directly work on RNA targets, in which the specialized DEL library and RNA-focused bioinformatics analysis workflow significantly improve the screening results. This methodology shows the feasibility of applying DEL to advanced RNA-targeted drug discovery efforts with its successful identification of a series of compounds that binding against G4-RNA. This platform unlocks a large chemical space, used to be only accessible by protein, can now be accessed by RNA. This can significantly enhance the hit finding efficiency for RNA. With DEL technology now available to rapidly generate RNA binding molecules, we foresee the acceleration in RNA targeting studies including finding novel tool molecule to study RNA mechanism-of-action, profiling RNA-small molecule binding interface and identifying novel chemical matters that can modulate RNA activities. Together with other progressions of the RNA-targeting field in the future, there is a great chance to expand the druggable space in the human genome.

## ACKNOWLEDGEMENTS

We would like to thank CSU team in WuXi AppTec for assistance in synthesis of off-DNA compounds. We also thank Early Discovery team in WuXi biology for assistance in DEL selection experiments.

## AUTHOR CONTRIBUTIONS

Conceptualization, Z.M. and W.S.; Methodology, Z.M. and B.Z.; Investigation, Q.Zhu., R.Z, X.W., X.G.,W.F. and W.L.; Software: B.Z.,X.H. Writing – Original Draft, Z.M. and W.S.; Writing, Review & Editing, Z.M. and W.S.; Resources, L.X., Y.H., W.C., and Q.Zhang; Supervision, Z.Y., L.K., and W.S.

## Methods

### RESOURCE AVAILABILITY

#### Lead contact

Further information and requests for resources and reagents should be directed to and will be fulfilled by the lead contact.

#### Materials availability

The DNA encoded libraries in this work are available from the lead contact upon request. The sequences are described in the text and can be purchased through oligonucleotide synthesis sources.

#### Data and code availability

The sequences of the RNA tested in this manuscript are described in the text. Additional data reported in this paper will be shared by the lead contact upon request. Methodology and software used for analysis are as described in the text. The scripts and further details are available upon request. Any additional information required to reanalyze the data reported in this paper is available from the lead contact upon request.

## METHOD DETAILS

### Specialized DNA Encoded Library (“DEL Zipper”) production

To produce specialized DEL (named as “DEL Zipper”), single-stranded region of DEL DNA was blunted using T4 DNA polymerase kit (NEB M0203L) in 150 μl 1X NEBuffer r2.1 supplemented with 100 μM of each dNTP. 1unit T4 DNA Polymerase per microgram DEL DNA was added. The reaction system was incubated 4 hrs at 12°C then stopped by heating to 75°C for 20 minutes. The products were isolated by ethanol precipitation with 75% v/v of ethanol in water and 125 mM of NaCl. Then the supernatant was pipetted out and the remaining ethanol was evaporated. Finally, the solid residue was re-dissolved in ddH2O. Gel electrophoresis was performed to confirm the conversion is successful.

### RNA Production for DEL Affinity Selection

Biotin-TEG tagged G-quadruplexes (G4) wild-type RNA (5’ biotin-TEG-UUUUGG GCUGCGG GUGGGCGGGA-3’) and 5’ Biotin-TEG tagged G4 mutant RNA (5’ biotin-TEG-UUUUGAG CUGCGAGUGAGCGAGA-3’) were synthesized by General Biol and annealed in 50 mM Tris, 300mM KCl, pH 7.5 by heating to 95℃ 5 min, on ice 2 min, then 37℃ 30 min.

Biotin-TEG tagged THF riboswitch RNA (5’ biotin-TEG-GGACAGAGUAGGUAAACGUGCG UAAAGUGCCUGAGGGACGGGGAGUUGUCCUCAGGACGAACACCGAAAGGUGGCGGUACGU UUACCGCAUCUCGCUGUUC-3’) was synthesized by GenScript and annealed in 100 mM HEPES, 100 mM NaCl, 10 mM MgCl_2_, pH 7.4 by heating to 95℃, 5 min on ice 2 min, then 37℃ 30 min.

### RNA immobilization test

Relative amount of biotinylated and non-biotinylated RNA was annealed as described above. Then the RNA was incubated with Pierce™ Streptavidin Magnetic Beads (20 μL, Thermo Fisher Scientific 88817) for 30mins at RT. After that, the RNA-beads mixture solution was placed into a DynaMag-2 magnet and corresponding supernatant was collected as our flow through (FT).

Then, the Beads were resuspended in the elution buffer (1x NuPAGE™ Sample Reducing Agent (ThermoFisher NP0009), 20 mM Tris, pH 7.5, 10 mM EDTA, 0.05% Tween 20) and heated to 95℃ for 10min. The supernatants were collected as retained on beads. The retained-on beads, FT and Input were loaded into agarose gel contains SYBR-Safe and further visualized by Bio-Rad Gel imager. Detection pipeline for DNA-RNA interaction and DNA motif For the pool ID +UMI region (ssDNA for conventional DEL and blunted dsDNA for DEL Zipper), motifs in the UMI region were classified based on copy of a single UMI and TOP100 UMI motifs of each sample were fragmented using a sliding window of 5-mer size and aligned against the sequence of corresponding RNA. Position and copy information of alignment were subjected to GraphPad to perform analysis. TOP10000 UMI motifs were descending ranked and accumulated to draw copy accumulation curve.

For the coding region, we developed an inhouse pipeline for detecting coding region Hoogsteen base-pairing motifs from warheadless libraries and removed all the potential coding region motif signals (Fig. 1B & Supplementary Fig. 1). Warheadless libraries, which were also called DNA only libraries, were acted as the representative of BB tags to find motifs, as they contained all BB tags in mono-synthon level of DEL. The relative fold change of a mono-synthon to average copy count of corresponding mono-synthons was calculated. Mono-synthons with fold change greater than 10 were considered as motifs in the coding region (Supplementary Fig. 1 Motif warning pipeline). DELs that sharing the same sequence with predicted motifs were labelled in final report and removed accordingly. Predicted motifs were fragmented using a sliding window of 5-mer size and aligned against the sequence of corresponding RNA using an in-house pipeline. Related scripts as well as instructions can be downloaded from github (https://github.com/WXDEL/RNA.git). Position and copy information of alignment was subjected to GraphPad to perform analysis (Supplementary Fig. 1 RNA-focused pipeline).

### RNA-DEL Affinity Selection

Pierce™ Streptavidin Magnetic Beads (20 μL, Thermo Fisher Scientific 88817) for each condition were washed three times with 200 μL of selection buffer (20 mM Tris, 300 mM KCl, 20 mM MgCl_2_, 0.05% Tween 20, 0.1 mg/mL sheared salmon sperm DNA [Invitrogen AM9680], pH 7.5) and placed into a DynaMag-2 magnet to collect the beads and discard the supernatant. 50 μM annealed 5’ Biotin-TEG tagged RNA was diluted to 200 pmol or 400 pmol in 100 μL 1x selection buffer. Then, RNA was immobilized on Pierce™ Streptavidin Magnetic Beads for 0.5 h at room temperature, followed by two washes (each 5 min) in 100 μL biotin blocking buffer (20 mM Tris, 300 mM KCl, 20 mM MgCl_2_, 0.1 mg/ml sssDNA, 0.05% Tween 20, 20 μM biotin, pH 7.5) at room temperature. Then RNA-matrix complex and the Conventional DEL or the “DNA Zipper” DEL were incubated in 100 μL of selection buffer for 1 h at room temperature, then washed in 200 μL selection buffer three times. To obtain eluted samples, 40 μL of selection buffer was added and incubated at 95 °C for 10 min on a Thermomixer (Eppendorf) with 800 rpm mixing. The eluted samples were purified by QIAquick PCR Purification Kit (Qiagen 28104). The remaining was used as the input for the next round of selection with the same procedure from the first round of selection. To exclude background signal of DEL to affinity matrix, the independent incubation of DEL without RNA was also conducted under the same condition as negative control.

After each round, the output was quantified by qPCR using PowerUp™ SYBR™ Green Master Mix-UDG (ThermoFisher 957005). When qPCR results showed that total copies of each purified sample were around 10^7^, the selection was done and the output was amplified by PCR using Platinum™ Taq DNA Polymerase High Fidelity kit (ThermoFisher 11304029) and UDG Enzyme (NEB M5508), then sequencing was performed for PCR amplified samples on an Illumina NovaSeq DNA sequencer. FastQ data were quality filtered, decoded, the unique molecule identifier (UMI) corrected, and copy counted by an in-house DEL decoding pipeline and motifs were detected by the motif warning pipeline described above.

For pre-selection, around 2E11 copies of tool compound or SHP-blank were used, instead of DEL and other materials and methods are the same as RNA-DEL Affinity Selection. After only one round, the output was quantified by qPCR to calculate recovery rate.

### Post-screening data analysis with Spotfire

The post-screening data analysis was proceeded with the Spotfire software. The data was imported into the 3D scatter plot in Spotfire, and the X axis, Y axis, Z axis of the plot were set to the BB ID used in each cycle of library respectively. The size of dot in plot was sized by corresponding enrichment value. After set the filter of the corresponding enrichment value, the binder identification and compound selection are carried out according to the pattern and data performance in the cubic plot.

### Surface plasmon resonance (SPR)

SPR experiments were performed on Biacore 8K instruments (GE Healthcare). Immobilization of target RNA was carried out at 25 °C on a SA chip (streptavidin, GE Healthcare, 29699621) using biotin tag in running buffer, containing 520 mM Tris, pH 7.5, 300 mM KCl, 20 mM MgCl_2_, 0.05% Tween 20, 2% DMSO. The annealed target RNA (prepared at 450 nM in immobilization buffer with 520 mM Tris, pH 7.5, 300 mM KCl, 20 mM MgCl_2_, 0.05% Tween 20) were streptavidin-coupled to a density of 2500 to 3000 Response Units (RU). Sensorgrams from reference surfaces and blank injections were subtracted from the raw data prior to data analysis using Biacore 8K evaluation software. Sensorgrams recorded at different compound concentrations in multi-cycle experiments were fitted using a 1:1 interaction model, with a term for mass-transport included.

### Microscale Thermophoresis (MST) assay

Annealed biotinylated RNA was labeled by the Biotinylated Target Labeling Kit (NanoTemper Technologies, NT-L020) in labeling buffer containing 20 mM Tris, pH 7.5, 300 mM KCl, and 5 mM MgCl_2_. Dispense the labeled RNA and mix with the serial diluted compound solutions the in 384-well plate(s). The MST assay buffer contained 20 mM Tris, pH 7.5, 300 mM KCl, 5 mM MgCl_2_ and 1% DMSO. The final concentration of RNA was at 30 nM. After incubating for 15 min at room temperature, the samples were loaded into Monolith Premium Capillary Chips (NanoTemper Technologies). Binding reactions were measured using Monolith X instrument (NanoTemper Technologies) at 25°C, medium MST power and 100% LED power. The data was analyzed by Spectral shift-based method^34^. The K_d_ fit function of the NanoTemper Analysis Software MO Affinity Analysis (v 3.0.6) was used to fit the curve and simulated the K_d_ value.

### Affinity Selection Mass Spectrometry (ASMS)

RNA binding assays were prepared in a 30 μL volume in 384-well plates. The incubation buffer was 20 mM Tris, pH7.5, 300 mM KCl, 10 mM MgCl_2_. 30 μM annealed RNA and 10 µM compounds were incubated for 30 mins, final DMSO concentration was 2%, all the compounds were checked by MS before used. Sample was injected by autosampler for two-dimensional LC-MS system. First dimension was used for binder/non-binder separation with high-throughput SEC column (PolyLC Inc., 052HY0502), SEC buffer was 20 mM Tris, pH7.5, 300 mM KCl, 10 mM MgCl_2_.

Second dimension was used for compounds elution by C18 column (Agilent,857768-901), mobile phase: 0.1% formic acid-water (A) and 0.1% formic acid-acetonitrile (B), gradient elute. Agilent 6546 QTOF was used for the compound’s detection, polarity was positive mode, mass detection range was 100-1700. Masshunter workstation was used for the data collection and analysis. Based on the noise signal, MS signal <3000 (Peak height) was considered as non-significant, mass threshold of peak identification was 5ppm.

### Detailed description of the synthesis of compounds

#### compound 1a-10b

The general procedure for the synthesis of compounds 1a to 1i was showed in scheme S1.

**Scheme S1.**
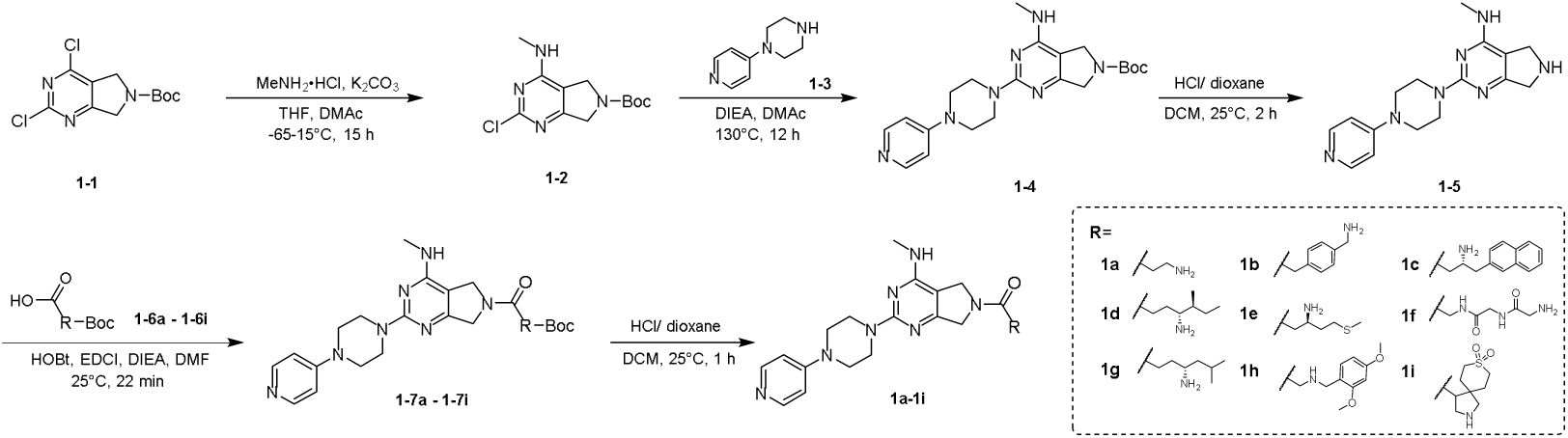
General procedures for the synthesis of compounds 1a to 1i.

Synthesis of 3-amino-1-(4-(methylamino)-2-(4-(pyridin-4-yl)piperazin-1-yl)-5,7-dihydro-6H-pyrrolo[3,4-d]pyrimidin-6-yl)propan-1-one (1a): Preparation of tert-butyl 2-chloro-4-(methylamino)-5,7-dihydro-6H-pyrrolo[3,4-d]pyrimidine-6-carboxylate (1-2)

To a solution of tert-butyl 2,4-dichloro-5,7-dihydro-6H-pyrrolo[3,4-d]pyrimidine-6-carboxylate (1.0 g, 3.45 mmol, 1.00 eq) in tetrahydrofuran (5.0 mL), the mixture was stirred at −60°C for 1 h. Methanamine hydrochloride (349 mg, 5.17 mmol, 1.5 eq) and potassium carbonate (1.43 g, 10.3 mmol, 3.00 eq) in dimethylacetamide (5.0 mL) was added in the mixture. The mixture was stirred at −60∼ 0°C for 12 h. Methanamine hydrochloride (349 mg, 5.17 mmol, 1.50 eq) was adeed in the mitxure and stirred at −60∼15°C for 2 h. The mixture was poured into water (50.0 mL) and extracted with ethyl acetate (30.0 mL*3). The combined organic layers were dried over sodium sulfate, filtered and concentrated under reduced pressure to give a residue. The residue was purified by flash silica gel chromatography. tert-butyl 2-chloro-4-(methylamino)-5,7-dihydro-6H-pyrrolo[3,4-d]pyrimidine-6-carboxylate (1-2, 550 mg, 1.93 mmol, 56.0% yield) was obtained as a white solid. H NMR (400 MHz, DMSO-d_6_): δ = 8.04 - 7.64 (m, 1H), 4.45 - 4.14 (m, 4H), 2.90 - 2.77 (m, 3H), 1.45 (d, J = 5.2 Hz, 9H).

#### Preparation of tert-butyl 4-(methylamino)-2-(4-(pyridine-4-yl)piperazin-1-yl)-5,7-dihydro-6H-pyrrolo[3,4-d]pyrimidine-6-carboxylate (1-4)

To a solution of tert-butyl 2-chloro-4-(methylamino)-5,7-dihydro-6H-pyrrolo[3,4-d]pyrimidine-6-carboxylate (1-2, 520 mg, 1.83 mmol, 1.00 eq), 1-(pyridin-4-yl)piperazine (1-3, 447 mg, 2.74 mmol, 1.50 eq) in dimethylacetamide (5.0 mL) was added N,N-diisopropylethylamine (708 mg, 5.48 mmol, 954 uL, 3.00 eq). The mixture was stirred at 130 °C for 12 h. The mixture was poured into water (50.0 mL) and extracted with dichloromethane (30.0 mL*3). The combined organic layers were dried over sodium sulfate], filtered and concentrated under reduced pressure to give a residue. The residue was purified by column chromatography (SiO2, Dichloromethane: Methanol=10:1). Tert-butyl 4-(methylamino)-2-(4-(pyridine-4-yl)piperazin-1-yl)-5,7-dihydro-6H-pyrrolo[3,4-d]pyrimidine-6-carboxylate (1-4, 160 mg, 388 umol, 21.2% yield) was obtained as a brown solid. H NMR (400 MHz, DMSO-d_6_): δ = 8.18 (d, J = 5.6 Hz, 2H), 7.06 - 6.95 (m, 1H), 6.90 (d, J = 5.6 Hz, 2H), 4.29 - 4.19 (m, 4H), 3.80 (s, 4H), 3.59 (s, 4H), 2.83 (t, J = 4.8 Hz, 3H), 1.44 (d, J = 4.0 Hz, 9H).

#### Preparation of (S)-N-(1-cyclohexyl-2-(methylamino)-2-oxoethyl)-7-(3,5-dimethylphenyl)-4-methoxy-1H-indole-2-carboxamide (1-5)

To a solution of tert-butyl 4-(methylamino)-2-(4-(pyridin-4-yl)piperazin-1-yl)-5,7-dihydro-6H-pyrrolo [3,4-d]pyrimidine-6-carboxylate (1-4, 160 mg, 388 umol, 1.00 eq) in dichloromethane (2.0 mL) was added hydrochloric acid /dioxane (4 M, 2.0 mL, 20.5 eq). The mixture was stirred at 25°C for 2 h. The reaction mixture was concentrated under reduced pressure to give a residue. N-methyl-2-(4-(pyridin-4-yl) piperazin-1-yl)-6,7-dihydro-5H-pyrrolo[3,4-d]pyrimidin-4-amine (1-5, 200 mg, crude, hydrochloric acid) was obtained as a brown solid. H NMR (400 MHz, DMSO-d_6_): δ = 13.95 - 13.23 (m, 1H), 10.09 - 9.99 (m, 1H), 8.33 - 8.08 (m, 2H), 7.47 - 6.92 (m, 2H), 4.36 - 4.21 (m, 4H), 3.94 (s, 4H), 3.83 (s, 4H), 2.94 - 2.87 (m, 3H).

#### Preparation of 7 - tert-butyl (3-(4-(methylamino)-2-(4-(pyridin-4-yl)piperazin-1-yl)-5,7-dihydro-6H-pyrrolo[3,4-d]pyrimidin-6-yl)-3-oxopropyl)carbamate (1-7a)

To a solution of 3-(tert-butoxycarbonylamino)propanoic acid (1-6a, 45.3 mg, 239 umol, 1.00 eq) in dimethyl formamide (1.0 mL) was added N,N-diisopropylethylamine (92.8 mg, 718 umol, 125 uL, 3.00 eq), hydroxybenzotriazole (48.5 mg, 359 umol, 1.50 eq) and 1-(3-dimethylaminopropyl)-3-ethylcarbodiimide hydrochloride (1-5, 68.8 mg, 359 umol, 1.50 eq). The mixture was stirred at 25 °C for 2 min. N-methyl-2-(4-(pyridin-4-yl)piperazin-1-yl)-6,7-dihydro-5H-pyrrolo[3,4-d]pyrimidin-4-amine (100 mg, 287.48 umol, 1.2 eq, hydrochloric acid) was added in the mixture stirred at 25°C for 20 min. The reaction mixture was diluted with water 30.0 mL and extracted with n-butyl alcohol (30.0 mL * 3). The combined organic layers were dried over sodium sulfate, filtered and concentrated under reduced pressure to give a residue. Tert-butyl(3-(4-(methylamino)-2-(4-(pyridin-4-yl)piperazin-1-yl)-5,7-dihydro-6H-pyrrolo[3,4-d] pyrimidin-6-yl)-3-oxopropyl)carbamate (1-7a, 250 mg, crude) was obtained as a black solid. LC-MS: R_t_ = 0.274 min, [M+H]^+^ = 483.3.

#### Preparation of 3-amino-1-(4-(methylamino)-2-(4-(pyridin-4-yl)piperazin-1-yl)-5,7-dihydro-6H-pyrrolo[3,4-d]pyrimidin-6-yl)propan-1-one (1a)

To a solution of tert-butyl (3-(4-(methylamino)-2-(4-(pyridin-4-yl)piperazin-1-yl)-5,7-dihydro-6H-pyrrolo [3,4-d]pyrimidin-6-yl)-3-oxopropyl)carbamate (1-7a, 150 mg, 310 umol, 1.00 eq) in dichloromethane (2.0 mL) was added hydrochloric acid/dioxane (4 M, 1.50 mL, 19.3 eq). The mixture was stirred at 25°C for 1 h. The reaction mixture was concentrated under reduced pressure to give a residue. The residue was purified by prep-HPLC (column: Welch Xtimate C18 150*25mm*5um;mobile phase: [water(trifluoroacetic acid)-acetonitrile];B%: 0%-20%,10min). 3-amino-1-(4-(methylamino)-2-(4-(pyridin-4-yl)piperazin-1-yl)-5,7-dihydro-6H-pyrrolo[3,4-d]pyrimidin-6-yl)propan-1-one (1a, 16.2 mg, 32.3 umol, 10.3% yield, 99% purity, trifluoroacetic acid) was obtained as a white solid. H NMR: (400 MHz, MeOD-d_4_):, δ = 8.29 - 8.15 (m, 2H), 7.19 (d, J = 7.6 Hz, 2H), 4.80 (s, 1H), 4.68 (s, 1H), 4.62 (s, 1H), 4.50 (s, 1H), 4.12 - 4.05 (m, 4H), 4.01 - 3.94 (m, 4H), 3.28 (t, J = 6.0 Hz, 2H), 3.09 (d, J = 4.4 Hz, 3H), 2.83 (td, J = 6.4, 8.8 Hz, 2H). HPLC: 99% purity. MS: [M+H]^+^ = 383.1.

Synthesis of 2-[4-(aminomethyl)phenyl]-1-[4-(methylamino)-2-[4-(4-pyridyl)piperazin-1-yl]-5,7-dihydropyrrolo[3,4-d]pyrimidin-6-yl]ethanone (1b): White solid, 16.18% yield. H NMR (400 MHz, DMSO-d_6_): δ = 8.36 (s, 1H), 8.18 (d, J = 6.4 Hz, 2H), 7.38 - 7.31 (m, 2H), 7.30 - 7.25 (m, 2H), 7.05 (t, J = 4.8 Hz, 1H), 6.85 (d, J = 6.4 Hz, 2H), 4.56 (s, 2H), 4.31 (d, J = 7.2 Hz, 2H), 3.89 (s, 2H), 3.80 (s, 4H), 3.71 (d, J = 7.2 Hz, 2H), 3.43 - 3.34 (m, 6H), 2.85 (dd, J = 4.4, 8.4 Hz, 3H). HPLC: 96.22% purity. MS:[M+H]^+^ = 459.4.

#### Synthesis of (R)-3-amino-1-(4-(methylamino)-2-(4-(pyridin-4-yl)piperazin-1-yl)-5,7-dihydro-6H-pyrrolo[3,4-d]pyrimidin-6-yl)-4-(naphthalen-2-yl)butan-1-one (1c)

White solid, 8.95% yield. H NMR (400 MHz, DMSO-d_6_): δ = 13.95 - 13.66 (m, 1H), 8.36 - 8.20 (m, 5H), 7.93 - 7.82 (m, 3H), 7.79 (d, J = 6.4 Hz, 1H), 7.51 - 7.42 (m, 3H), 7.22 ( d, J = 7.6 Hz, 2H), 4.65 ( s, 1H), 4.55 - 4.34 (m, 2H), 4.32 - 4.25 (m, 1H), 4.21 - 4.12 (m, 1H), 4.00 ( s, 4H), 3.87 ( s, 5H), 3.07 (dd, J = 13.6, 8.4 Hz, 1H), 2.92 (d, J = 3.6 Hz, 3H), 2.79 - 2.69 (m, 1H), 2.69 - 2.60 (m, 1H). HPLC: 98% purity. MS: [M+H]^+^ = 523.1.

#### Synthesis of ((4R,5S)-4-Amino-5-methyl-1-(4-(methylamino)-2-(4-(pyridin-4-yl)piperazin-1-yl)-5,7-dihydro-6H-pyrrolo[3,4-d]pyrimidin-6-yl)heptan-1-one (1d)

White solid, 72.8% yield. H NMR (400 MHz, DMSO-d_6_): δ 13.78 (s, 1H), 8.34 - 8.21 (m, 2H), 8.08 (s, 3H), 7.25 - 7.10 (d, J = 7.2 Hz, 2H), 4.87 - 4.64 (m, 1H), 4.63 - 4.43 (m, 2H), 4.29 (s, 1H), 3.97 (s, 4H), 3.83 (br s, 4H), 3.09 - 3.00 (m, 1H), 2.95 - 2.85 (m, 3H), 2.57 - 2.51 (m, 2H), 1.86 - 1.75 (m, 1H), 1.71 - 1.51 (m, 2H), 1.49 - 1.28 (m, 1H), 1.24 - 1.09 (m, 1H), 0.92 - 0.72 (m, 6H). HPLC: 98% purity. MS: [M+H]^+^ = 453.3.

#### Synthesis of (R)-3-amino-1-(4-(methylamino)-2-(4-(pyridin-4-yl)piperazin-1-yl)-5,7-dihydro-6H-pyrrolo[3,4-d]pyrimidin-6-yl)-5-(methylthio)pentan-1-one (1e)

Yellow gum, 16.04% yield. H NMR (400 MHz, DMSO-d_6_): δ = 13.78 ( s, 1H), 8.30 (d, J = 5.6 Hz, 2H), 8.22 - 8.07 (m, 3H), 7.28 - 7.16 (m, 2H), 4.83 - 4.69 (m, 1H), 4.56 ( s, 2H), 4.38 ( s, 1H), 4.01 ( s, 4H), 3.87 ( s, 4H), 2.94 ( t, J = 3.6 Hz, 3H), 2.86 - 2.65 (m, 3H), 2.60 (br t, J = 7.4 Hz, 2H), 2.06 (s, 3H), 2.00 - 1.85 (m, 2H). HPLC: 96% purity. MS: [M+H]^+^ = 457.1. SFC: 1.999min, 2.117min.

#### Synthesis of 2-amino-N-(2-((2-(4-(methylamino)-2-(4-(pyridin-4-yl)piperazin-1-yl)-5,7-dihydro-6H-pyrrolo[3,4-d]pyrimidin-6-yl)-2-oxoethyl)amino)-2-oxoethyl)acetamide (1f)

White solid, 69.24% yield, HPLC: 94.337% purity. MS: [M+H]^+^ = 483.26.

#### Synthesis of (R)-4-Amino-6-methyl-1-(4-(methylamino)-2-(4-(pyridin-4-yl)piperazin-1-yl)-5,7-dihydro-6H-pyrrolo[3,4-d]pyrimidin-6-yl)heptan-1-one (1g)

Brown solid, 98.2% yield. H NMR (400 MHz, DMSO-d_6_): δ 13.99 (s, 1H), 9.19 - 8.51 (m, 1H), 8.45 - 8.22 (d, J = 5.6 Hz, 2H), 8.14 (s, 2H), 7.39 - 7.05 ( d, J = 6.8 Hz, 2H), 4.98 - 4.70 (m, 1H), 4.69 - 4.68 (m, 2H), 4.40 - 4.31 (d, J = 14.4 Hz, 1H), 4.05 (s, 4H), 3.89 (s, 5H), 3.17 (s, 1H), 2.95 (s, 3H), 2.74 - 2.58 (m, 1H), 1.98 - 1.65 (m, 3H), 1.57 - 1.32 (m, 2H), 1.05 - 0.65 (d, J = 5.6 Hz, 6H). HPLC: 99.1% purity. MS: [M+H]^+^ = 453.3.

#### Synthesis of 2-((2,4-dimethoxybenzyl)amino)-1-(4-(methylamino)-2-(4-(pyridin-4-yl)piperazin-1-yl)-5,7-dihydro-6H-pyrrolo[3,4-d]pyrimidin-6-yl)ethan-1-one (1h)

White oil, 42.63% yield. HPLC: 99.481% purity. MS: [M+H]^+^ = 519.28.

#### Synthesis of (8,8-dioxido-8-thia-2-azaspiro[4.5]decan-4-yl)(4-(methylamino)-2-(4-(pyridin-4-yl)piperazin-1-yl)-5,7-dihydro-6H-pyrrolo[3,4-d]pyrimidin-6-yl)methanone (1i)

White solid, 91.30% yield. HPLC: 94.952% purity. MS: [M+H]^+^ = 527.25. The general procedure for the synthesis of compounds 2a to 2i is showed in scheme S2.

**Scheme S2.**
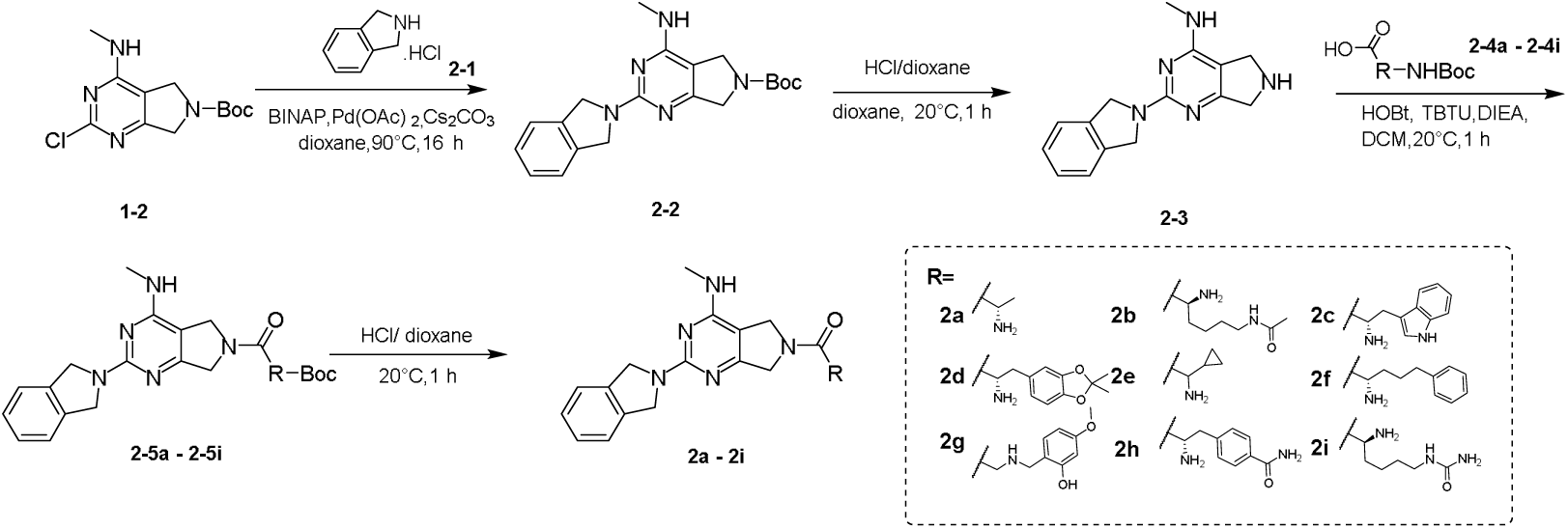
General procedures for the synthesis of compounds 2a to 2i.

#### Synthesis of (S)-2-amino-1-(2-(isoindolin-2-yl)-4-(methylamino)-5,7-dihydro-6H-pyrrolo[3,4-d]pyrimidin-6-yl)propan-1-one (2a): Preparation of tert-butyl 2-(isoindolin-2-yl)-4-(methylamino)-5,7-dihydro-6H-pyrrolo[3,4-d]pyrimidine-6-carboxylate (2-2)

A mixture of tert-butyl 2-chloro-4-(methylamino)-5,7-dihydro-6H-pyrrolo[3,4-d]pyrimidine-6-carboxylate (1-2, 2.50 g, 8.78 mmol, 1.00 eq), isoindoline (2-1, 1.37 g, 8.78 mmol, 1.30 mL, 1.00 eq, hydrochloride), [1-(2-diphenylphosphanyl-1-naphthyl)-2-naphthyl]-diphenyl-phosphane (1.09 g, 1.76 mmol, 0.200 eq), diacetoxypalladium (197 mg, 877 umol, 0.100 eq) and cesium carbonate (8.58 g, 26.3 mmol, 3.00 eq) in dioxane (50.0 mL) was degassed and purged with nitrogen for 3 times, and then the mixture was stirred at 90°C for 3 h under nitrogen atmosphere. The reaction mixture was diluted with water 200 mL and extracted with ethyl acetate 600.0 mL (200.0 mL * 3). The combined organic layers were washed with brine 200.0 mL, dried over sodium sulfate, filtered and concentrated under reduced pressure to give a residue. The residue was purified by flash silica gel chromatography. Compound tert-butyl2-(isoindolin-2-yl)-4-(methylamino)-5,7-dihydro-6H-pyrrolo[3,4-d]pyrimidine-6-carboxylate(2-2, 0.700 g, 1.91 mmol, 21.7% yield) was obtained as a gray solid. HNMR (400 MHz, DMSO-d_6_): δ = 7.42 - 7.35 (m, 4H), 7.29 - 7.24 (m, 5H), 4.77 (s, 4H), 4.32 - 4.22 (m, 4H), 2.88 (d, J = 5.2 Hz, 3H), 1.45 (d, J = 2.8 Hz, 9H).

#### Preparation of 2-(isoindolin-2-yl)-N-methyl-6,7-dihydro-5H-pyrrolo[3,4-d] pyrimidin-4-amine (2-3)

To a solution of tert-butyl 2-(isoindolin-2-yl)-4-(methylamino)-5,7-dihydro-6H-pyrrolo[3,4-d] pyrimidine-6-carboxylate (2-3, 0.300 g, 816 umol, 1.00 eq) in dioxane (2.00 mL) was added hydrochloric acid (4 M in dioxane) (4 M, 5.0 mL, 24.5 eq). The mixture was stirred at 25°C for 16 h. The reaction mixture was concentrated under reduced pressure to give a residue. 2-(isoindolin-2-yl)-N-methyl-6,7-dihydro-5H-pyrrolo[3,4-d] pyrimidin-4-amine (337 mg, crude) was obtained as a gray solid. HNMR (400 MHz, DMSO-d_6_): δ =10.42 (s, 2H), 9.14 - 8.82 (m, 1H), 7.43 (d, J = 1.6 Hz, 2H), 7.36 (dd, J = 3.2, 5.6 Hz, 2H), 4.93 (s, 4H), 4.47 (s, 2H), 4.28 (s, 2H), 2.99 (d, J = 3.6 Hz, 3H).

#### Preparation of tert-butyl (S)-(1-(2-(isoindolin-2-yl)-4-(methylamino)-5,7-dihydro-6H-pyrrolo[3,4-d]pyrimidin-6-yl)-1-oxopropan-2-yl)carbamate (2-5a)

A mixture of 2-(isoindolin-2-yl)-N-methyl-6,7-dihydro-5H-pyrrolo[3,4-d]pyrimidin-4-amine (2-3, 150 mg, 561 umol, 1.00 eq), (2S)-2-(tert-butoxycarbonylamino)propanoic acid (2-4a, 106 mg, 561 umol, 1.00 eq), N,N-diisopropylethylamine (217 mg, 1.68 mmol, 293 uL, 3.00 eq), [benzotriazol-1-yloxy(dimethylamino) methylidene]-dimethylazanium;hexafluorophosphate (255 mg, 673 umol, 1.20 eq) and 1-hydroxybenzotriazole (151 mg, 1.12 mmol, 2.00 eq) in dichloromethane (5.0 mL) was degassed and purged with nitrogen for 3 times, and then the mixture was stirred at 25°C for 1 h under nitrogen atmosphere. The reaction mixture was partitioned between water 20.0 mL and ethyl acetate 20.0 mL and extracted with ethyl acetate 60.0 mL (20.0 mL * 3). The organic phase was separated, washed with brine 20.0 mL, dried over sodium sulfate, filtered and concentrated under reduced pressure to give a residue. The residue was purified by flash silica gel chromatography. Tert-butyl (S)-(1-(2-(isoindolin-2-yl)-4-(methylamino)-5,7-dihydro-6H-pyrrolo[3,4-d] pyrimidin-6-yl)-1-oxopropan-2-yl) carbamate (120 mg, 273 umol, 48.7% yield) was obtained as a white solid. H NMR (400 MHz, DMSO-d_6_): δ =7.40 - 7.35 (m, 2H), 7.31 - 7.23 (m, 3H), 7.07 (t, J = 7.6 Hz, 1H), 6.98 (s, 1H), 4.77 (s, 5H), 4.43 - 4.13 (m, 4H), 2.90 (t, J = 4.8 Hz, 3H), 1.45 - 1.41 (m, 3H), 1.38 (s, 9H).

#### Preparation of compound-(S)-2-amino-1-(2-(isoindolin-2-yl)-4-(methylamino)-5,7-dihydro-6H-pyrrolo[3,4-d] pyrimidin-6-yl) propan-1-one (2a)

To a solution of tert-butyl (S)-(1-(2-(isoindolin-2-yl)-4-(methylamino)-5,7-dihydro-6H-pyrrolo[3,4-d] pyrimidin-6-yl)-1-oxopropan-2-yl) carbamate (2-5a, 120 mg, 273 umol, 1.00 eq) in dioxane (1.00 mL) was added HCl/dioxane (4 M, 2.0 mL, 29.2 eq). The mixture was stirred at 25°C for 1 h. The reaction mixture was concentrated under reduced pressure to give a residue. The residue was purified by prep-HPLC (column: Phenomenex luna C18 150*25mm* 10um; mobile phase: [water (FA)-ACN]; B%: 1%-28%,15min). (S)-2-amino-1-(2-(isoindolin-2-yl)-4-(methylamino)-5,7-dihydro-6H-pyrrolo[3,4-d] pyrimidin-6-yl) propan-1-one (11.74 mg, 33.14 umol, 12.11% yield, 95.533% purity) was obtained as a off-white solid. H NMR (400 MHz, DMSO-d_6_): δ =8.32 (s, 1H), 7.43 - 7.35 (m, 2H), 7.33 - 7.25 (m, 2H), 7.08 - 6.92 (m, 1H), 4.91 - 4.62 (m, 5H), 4.58 - 4.47 (m, 1H), 4.43 - 4.25 (m, 2H), 3.83 - 3.62 (m, 3H), 2.94 - 2.88 (m, 3H), 1.26 - 1.17 (m, 3H). HPLC: 95.533% purity. MS: [M+H]^+^ = 339.3. SFC: R_t_ = 1.853 min.

#### Synthesis of (S)-N-(5-amino-6-(2-(isoindolin-2-yl)-4-(methylamino)-5,7-dihydro-6H-pyrrolo[3,4-d] pyrimidin-6-yl)-6-oxohexyl) acetamide (2b)

White solid, 46.71% yield. H NMR (400 MHz, DMSO-d_6_): δ = 9.40 - 8.84 (m, 1H), 8.48 (s, 3H), 8.08 - 7.96 (m, 1H), 7.47 - 7.32 (m, 4H), 5.14 - 4.58 (m, 7H), 4.52 - 4.33 (m, 1H), 4.15 - 3.93 (m, 1H), 3.03 ( d, J = 5.2 Hz, 2H), 2.96 ( s, 1H), 2.99 ( d, J = 4.4 Hz, 2H), 1.88 - 1.73 (m, 5H), 1.43 ( d, J = 6.4 Hz, 4H). MS: [M+H]^+^ = 438.3. SFC: Rt = 1.521,1.558 min.

#### Synthesis of (S)-2-amino-3-(1H-indol-3-yl)-1-(2-(isoindolin-2-yl)-4-(methylamino)-5,7-dihydro-6H-pyrrolo[3,4-d] pyrimidin-6-yl) propan-1-one (2c)

Yellow solid,12.31% yield. H NMR (400 MHz, DMSO-d_6_): δ = 11.12 ( s, 1H), 8.94 - 8.41 (m, 4H), 7.56 (dd, J = 4.4, 7.2 Hz, 1H), 7.47 - 7.25 (m, 6H), 7.08 - 6.97 (m, 2H), 4.96 ( s, 4H), 4.71 - 4.57 (m, 1H), 4.40 ( d, J = 17.6 Hz, 1H), 4.32 - 3.91 (m, 3H), 3.29 ( dd, J = 6.4, 10.4 Hz, 2H), 3.01 - 2.88 (m, 3H). MS: [M+H]^+^ = 454.1. SFC: Rt = 1.812,1.982 min.

#### Synthesis of (S)-2-amino-3-(2,2-dimethylbenzo[d][1,3] dioxol-5-yl)-1-(2-(isoindolin-2-yl)-4-(methylamino)-5,7-dihydro-6H-pyrrolo[3,4-d]pyrimidin-6-yl)propan-1-one (2d)

White solid, 10.00% yield. H NMR (400 MHz, DMSO-d_6_): δ = 7.42 - 7.34 (m, 2H), 7.32 - 7.26 (m, 2H), 6.96 ( dd, J = 4.8, 11.6 Hz, 1H), 6.77 - 6.61 (m, 3H), 4.87 - 4.67 (m, 4H), 4.61 - 4.50 (m, 1H), 4.37 - 4.22 (m, 2H), 4.22 - 3.94 (m, 1H), 3.75 - 3.53 (m, 1H), 2.90 (dd, J = 3.2, 4.4 Hz, 3H), 2.80 - 2.66 (m, 1H), 2.64 - 2.51 (m, 2H), 1.64 - 1.46 (m, 7H). MS: [M+H]^+^ = 487.2. SFC: Rt = 1.163,1.367 min.

#### Synthesis of 2-amino-2-cyclopropyl-1-[2-isoindolin-2-yl-4-(methylamino)-5,7-dihydropyrrolo[3,4-d]pyrimidin-6-yl]ethanone (2e)

White solid, 23.00% yield. MS: [M+H]^+^ = 365.

#### Synthesis of (S)-2-Amino-1-(2-(isoindolin-2-yl)-4-(methylamino)-5,7-dihydro-6H-pyrrolo[3,4-d] pyrimidin-6-yl)-5-phenylpentan-1-one (2f)

White solid, 10.1% yield. H NMR (400 MHz, MeOD-d_4_): δ 7.49 - 7.33 (m, 4H), 7.30 - 7.10 (m, 5H), 4.91 (s, 4H), 4.81 - 4.75 (m, 2H), 4.72 - 4.58 (m, 2H), 4.55 - 4.41 (m, 1H), 4.31 - 4.20 (m, 1H), 3.15 - 3.11 (d, J = 9.2 Hz, 3H), 2.82 - 2.59 (m, 2H), 2.01 - 1.90 (m, 1H), 1.88 - 1.76 (m, 2H). MS: [M+H]^+^ = 443.3.

#### Synthesis of 2-amino-2-cyclopropyl-1-[2-isoindolin-2-yl-4-(methylamino)-5,7-dihydropyrrolo[3,4-d] pyrimidin-6-yl]ethanone (2g)

White solid, 11.69% yield. MS: [M+H]^+^ = 460.53.

#### Synthesis of (S)-4-(2-Amino-3-(2-(isoindolin-2-yl)-4-(methylamino)-5,7-dihydro-6H-pyrrolo[3,4-d] pyrimidin-6-yl)-3-oxopropyl) benzamide (2h)

Yellow solid, 80.9% yield. ^1^H NMR (400 MHz, DMSO-d_6_): δ 7.97 - 7.75 (m, 3H), 7.44 - 7.23 (m, 7H), 7.08 - 6.94 (m, 1H), 4.87 - 4.61 (m, 4H), 4.46 - 4.20 (m, 2H), 4.19 - 4.00 (m, 1H), 3.08 - 3.02 (m, 1H), 2.92 - 2.80 (d, J = 4.4 Hz, 3H), 1.90 (s, 4H). HPLC: 93.0% purity. MS: [M+H]^+^ = 458.3.

#### Synthesis of (S)-1-(5-Amino-6-(2-(isoindolin-2-yl)-4-(methylamino)-5,7-dihydro-6H-pyrrolo[3,4-d] pyrimidin-6-yl)-6-oxohexyl) urea (2i)

White solid, 15.93% yield. H NMR (400 MHz, MeOH-d_4_): δ 7.36 - 7.23 (m, 4H), 4.81 - 4.42 (m, 8H), 3.76 - 3.65 (m, 1H), 3.15 – 3.01 (m, 5H), 1.77 - 1.70 (m, 1H), 1.64 - 1.39 (m, 5H). HPLC: 90.9% purity. MS: [M+H]^+^ = 439.2. The general procedure for the synthesis of compounds 3a and 3b is showed in scheme S3.

**Scheme S3.**
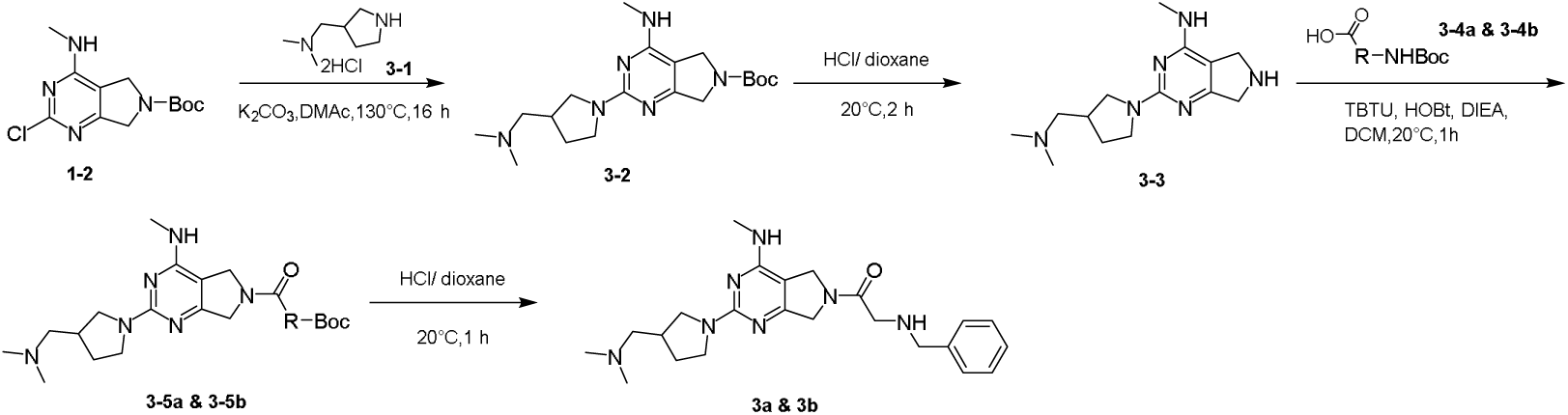
General procedures for the synthesis of compounds 3a and 3b.

#### Preparation of compound 3 - tert-butyl 2-(3-((dimethylamino)methyl)pyrrolidin-1-yl)-4-(methylamino)-5,7-dihydro-6H-pyrrolo[3,4-d]pyrimidine-6-carboxylate (3-2)

To a solution of tert-butyl 2-chloro-4-(methylamino)-5,7-dihydro-6H-pyrrolo[3,4-d] pyrimidine-6-carboxylate (1-2, 900 mg, 3.16 mmol, 1.00 eq) in N,N-dimethylacetamide (10.0 mL) was added potassium carbonate (1.75 g, 12.6 mmol, 4.00 eq) and N,N-dimethyl-1-pyrrolidin-3-yl-methanamine (3-1, 635 mg, 3.16 mmol, 1.00 eq, 2 hydrochloric). The mixture was stirred at 130°C for 16 h. The reaction mixture was poured into water (20.0 mL) and extracted with dichloromethane: isopropanol (5:1) 60 mL (20.0 mL * 3). The combined organic layers were washed with brine 20 mL, dried over sodium sulfate, filtered and concentrated under reduced pressure to give a residue. Tert-butyl 2-(3-((dimethylamino)methyl)pyrrolidin-1-yl)-4-(methylamino)-5,7-dihydro-6H-pyrrolo[3,4-d]pyrimidine-6-carboxylate (3-2, 1.10 g, 2.92 mmol, 92.4% yield) was obtained as a white solid. H NMR (400 MHz, DMSO-d_6_): δ= 6.79 (quin, J = 4.8 Hz, 1H), 4.30 - 4.11 (m, 3H), 3.71 - 3.45 (m, 2H), 3.16 - 3.04 (m, 1H), 2.94 (s, 4H), 2.84 - 2.76 (m, 6H), 2.38 (td, J = 7.2, 14.4 Hz, 1H), 2.28 - 2.17 (m, 2H), 2.14 (s, 5H), 2.02 - 1.86 (m, 5H), 1.64 - 1.52 (m, 1H), 1.44 (d, J = 5.2 Hz, 7H).

#### Preparation of compound-2-(3-((dimethylamino)methyl)pyrrolidin-1-yl)-N-methyl-6,7-dihydro-5H-pyrrolo[3,4-d]pyrimidin-4-amine (3-3)

To a solution of tert-butyl 2-(3-((dimethylamino)methyl) pyrrolidin-1-yl)-4-(methylamino)-5,7-dihydro-6H-pyrrolo[3,4-d]pyrimidine-6-carboxylate (3-2, 1.10 g, 2.92 mmol, 1.00 eq) in dioxane (10.0 mL) was added hydrochloric/dioxane (4 M, 10.0 mL, 13.69 eq). The mixture was stirred at 25°C for 1 h. The reaction mixture was concentrated under reduced pressure to give a residue. 2-(3-((dimethylamino)methyl) pyrrolidin-1-yl)-N-methyl-6,7-dihydro-5H-pyrrolo[3,4-d]pyrimidin-4-amine (3-3, 0.800 g, 2.89 mmol, 99.1% yield) was obtained as a gray solid.MS: [M+H]^+^ = 277.1.

#### Preparation of compound - tert-butyl benzyl(2-(2-(3-((dimethylamino)methyl)pyrrolidin-1-yl)-4-(methylamino)-5,7-dihydro-6H-pyrrolo[3,4-d]pyrimidin-6-yl)-2-oxoethyl)carbamate (3-5a)

A mixture of 2-(3-((dimethylamino)methyl)pyrrolidin-1-yl)-N-methyl-6,7-dihydro-5H-pyrrolo[3,4-d] pyrimidin-4-amine (3-3, 0.2 g, 723.64 umol, 1.00 eq), N-benzyl-N-(tert-butoxycarbonyl)glycine (3-4a, 191 mg, 724 umol, 1.00 eq), N,N-diisopropylethylamine (467 mg, 3.62 mmol, 630 uL, 5.00 eq), [benzotriazol-1-yloxy(dimethylamino)methylidene]-dimethylazanium;hexafluorophosphate (329 mg, 868 umol, 1.20 eq) and 1-hydroxybenzotriazole (195 mg, 1.45 mmol, 2.00 eq) in dichloromethane (6.0 mL) was degassed and purged with nitrogen for 3 times, and then the mixture was stirred at 25°C for 1 h under nitrogen _2_ atmosphere. The reaction mixture was diluted with water 20.0 mL and extracted with dichloromethane 60.0 mL (20.0 mL * 3). The combined organic layers were washed with brine 20 mL, dried over sodium sulfate, filtered and concentrated under reduced pressure to give a residue. The residue was purified by flash silica gel chromatography. Tert-butyl benzyl(2-(2-(3-((dimethylamino)methyl) pyrrolidin-1-yl)-4-(methylamino)-5,7-dihydro-6H-pyrrolo[3,4-d]pyrimidin-6-yl)-2-oxoethyl)carbamate (0.225 g, 429.66 umol, 59.37% yield) was obtained as a white solid. H NMR (400 MHz, DMSO-d_6_): δ= 7.37 - 7.30 (m, 3H), 7.30 - 7.23 (m, 5H), 4.46 - 4.36 (m, 5H), 4.30 ( d, J = 18.0 Hz, 2H), 3.90 ( d, J = 8.0 Hz, 1H), 3.67 - 3.60 (m, 1H), 3.56 - 3.52 (m, 1H), 3.14 - 3.06 (m, 2H), 3.04 - 2.94 (m, 1H), 2.86 - 2.77 (m, 3H), 2.44 - 2.37 (m, 2H), 2.27 (s, 6H), 1.35 (s, 9H), 1.21 (d, J = 6.4 Hz, 4H).

#### Preparation of 2-(benzylamino)-1-(2-(3-((dimethylamino)methyl)pyrrolidin-1-yl)-4-(methylamino)-5,7-dihydro-6H-pyrrolo[3,4-d]pyrimidin-6-yl)ethan-1-one (3a)

To a solution of tert-butylbenzyl(2-(2-(3-((dimethylamino)methyl)pyrrolidin-1-yl)-4-(methylamino)-5,7-dihydro-6H-pyrrolo[3,4-d]pyrimidin-6-yl)-2-oxoethyl)carbamate (3-5a, 0.200 g, 381 umol, 1.00 eq) in methanol (1.00 mL) was added hydrochloric acid (4 M in dioxane) (4 M, 2.0 mL, 20.9 eq). The mixture was stirred at 20°C for 1 h. The reaction mixture was concentrated under reduced pressure to give a residue. The residue was purified by prep-HPLC (column: Welch Xtimate C18 150*25mm*5um; mobile phase: [water (TFA)-ACN];B%: 0%-26%,10min). 2-(benzylamino)-1-(2-(3-((dimethylamino)methyl) pyrrolidin-1-yl)-4-(methylamino)-5,7-dihydro-6H-pyrrolo[3,4-d] pyrimidin-6-yl)ethan-1-one (85.23 mg, 185.40 umol, 48.54% yield, 92.134% purity) was obtained as a red gum. H NMR (400 MHz, DMSO-d_6_): δ= 9.89 - 9.40 (m, 2H), 7.55 - 7.40 (m, 5H), 4.68 - 4.33 (m, 3H), 4.24 - 3.97 (m, 5H), 3.84 (d, J = 12.4 Hz, 2H), 3.74 (s, 2H), 3.54 - 3.37 (m, 2H), 3.34 - 3.12 (m, 3H), 2.90 (d, J = 4.4 Hz, 2H), 2.83 (s, 4H), 2.77 - 2.63 (m, 1H), 2.27 - 2.06 (m, 1H), 1.88 - 1.61 (m, 1H). MS: [M+H]^+^ = 424.1.

#### Synthesis of 1-[2-[3-[(dimethylamino)methyl]pyrrolidin-1-yl]-4-(methylamino)-5,7-dihydropyrrolo[3,4-d]pyrimidin-6-yl]-2-piperazin-1-yl-ethanone (3b)

Yellow gum, 18.79% yield. ^1^H NMR (400 MHz, DMSO-d_6_): δ = 10.08 - 9.50 (m, 1H), 9.15 - 7.84 (m, 3H), 4.77 - 4.52 (m, 2H), 4.50 - 4.32 (m, 2H), 3.94 - 3.70 (m, 6H), 3.20 (d, J = 14.8 Hz, 6H), 2.93 (dd, J = 4.4, 8.0 Hz, 6H), 2.83 (s, 5H), 2.77 - 2.67 (m, 1H), 2.21 (s, 1H), 1.93 - 1.67 (m, 1H). MS: [M+H]^+^ = 402.29.

Detailed description of synthesis of the off-DNA compound 1a-10b for G4-RNA The general procedure for the synthesis of compounds 1a to 1i is showed in scheme S1.

**Scheme S1.**
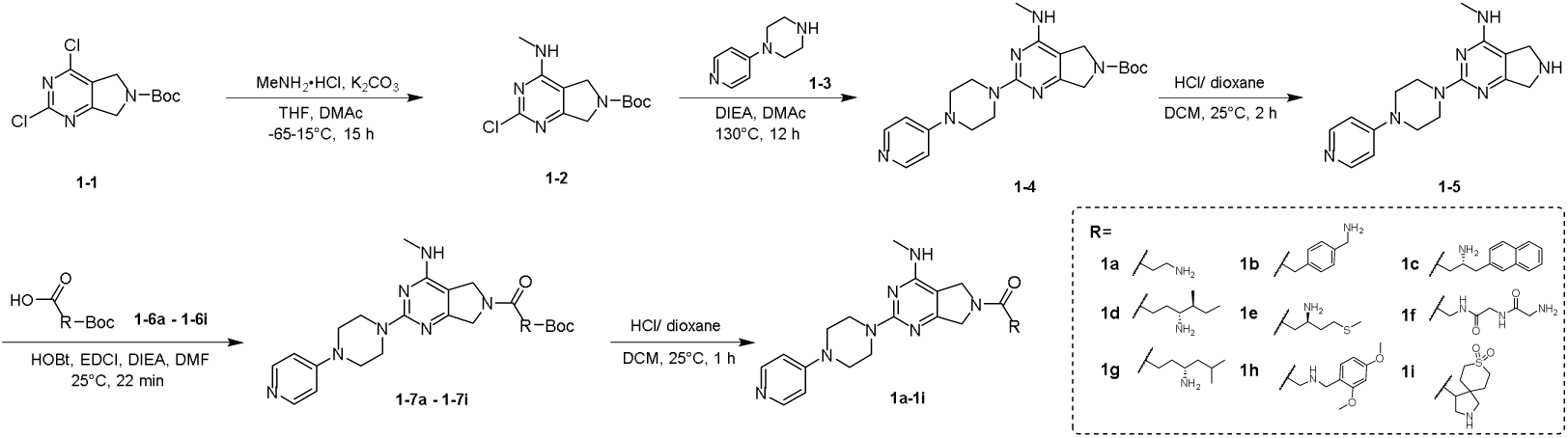
General procedures for the synthesis of compounds 1a to 1i.

#### Synthesis of 3-amino-1-(4-(methylamino)-2-(4-(pyridin-4-yl)piperazin-1-yl)-5,7-dihydro-6H-pyrrolo[3,4-d]pyrimidin-6-yl)propan-1-one (1a): Preparation of tert-butyl 2-chloro-4-(methylamino)-5,7-dihydro-6H-pyrrolo[3,4-d]pyrimidine-6-carboxylate (1-2)

To a solution of tert-butyl 2,4-dichloro-5,7-dihydro-6H-pyrrolo[3,4-d]pyrimidine-6-carboxylate (1.0 g, 3.45 mmol, 1.00 eq) in tetrahydrofuran (5.0 mL), the mixture was stirred at −60°C for 1 h. Methanamine hydrochloride (349 mg, 5.17 mmol, 1.5 eq) and potassium carbonate (1.43 g, 10.3 mmol, 3.00 eq) in dimethylacetamide (5.0 mL) was added in the mixture. The mixture was stirred at −60∼ 0°C for 12 h. Methanamine hydrochloride (349 mg, 5.17 mmol, 1.50 eq) was adeed in the mitxure and stirred at −60∼15°C for 2 h. The mixture was poured into water (50.0 mL) and extracted with ethyl acetate (30.0 mL*3). The combined organic layers were dried over sodium sulfate, filtered and concentrated under reduced pressure to give a residue. The residue was purified by flash silica gel chromatography. tert-butyl 2-chloro-4-(methylamino)-5,7-dihydro-6H-pyrrolo[3,4-d]pyrimidine-6-carboxylate (1-2, 550 mg, 1.93 mmol, 56.0% yield) was obtained as a white solid. H NMR (400 MHz, DMSO-d_6_): δ = 8.04 - 7.64 (m, 1H), 4.45 - 4.14 (m, 4H), 2.90 - 2.77 (m, 3H), 1.45 (d, J = 5.2 Hz, 9H).

#### Preparation of tert-butyl 4-(methylamino)-2-(4-(pyridine-4-yl)piperazin-1-yl)-5,7-dihydro-6H-pyrrolo[3,4-d]pyrimidine-6-carboxylate (1-4)

To a solution of tert-butyl 2-chloro-4-(methylamino)-5,7-dihydro-6H-pyrrolo[3,4-d]pyrimidine-6-carboxylate (1-2, 520 mg, 1.83 mmol, 1.00 eq), 1-(pyridin-4-yl)piperazine (1-3, 447 mg, 2.74 mmol, 1.50 eq) in dimethylacetamide (5.0 mL) was added N,N-diisopropylethylamine (708 mg, 5.48 mmol, 954 uL, 3.00 eq). The mixture was stirred at 130 °C for 12 h. The mixture was poured into water (50.0 mL) and extracted with dichloromethane (30.0 mL*3). The combined organic layers were dried over sodium sulfate], filtered and concentrated under reduced pressure to give a residue. The residue was purified by column chromatography (SiO2, Dichloromethane: Methanol=10:1). Tert-butyl 4-(methylamino)-2-(4-(pyridine-4-yl)piperazin-1-yl)-5,7-dihydro-6H-pyrrolo[3,4-d]pyrimidine-6-carboxylate (1-4, 160 mg, 388 umol, 21.2% yield) was obtained as a brown solid. H NMR (400 MHz, DMSO-d_6_): δ = 8.18 (d, J = 5.6 Hz, 2H), 7.06 - 6.95 (m, 1H), 6.90 (d, J = 5.6 Hz, 2H), 4.29 - 4.19 (m, 4H), 3.80 (s, 4H), 3.59 (s, 4H), 2.83 (t, J = 4.8 Hz, 3H), 1.44 (d, J = 4.0 Hz, 9H).

#### Preparation of (S)-N-(1-cyclohexyl-2-(methylamino)-2-oxoethyl)-7-(3,5-dimethylphenyl)-4-methoxy-1H-indole-2-carboxamide (1-5)

To a solution of tert-butyl 4-(methylamino)-2-(4-(pyridin-4-yl)piperazin-1-yl)-5,7-dihydro-6H-pyrrolo [3,4-d]pyrimidine-6-carboxylate (1-4, 160 mg, 388 umol, 1.00 eq) in dichloromethane (2.0 mL) was added hydrochloric acid /dioxane (4 M, 2.0 mL, 20.5 eq). The mixture was stirred at 25°C for 2 h. The reaction mixture was concentrated under reduced pressure to give a residue. N-methyl-2-(4-(pyridin-4-yl) piperazin-1-yl)-6,7-dihydro-5H-pyrrolo[3,4-d]pyrimidin-4-amine (1-5, 200 mg, crude, hydrochloric acid) was obtained as a brown solid. H NMR (400 MHz, DMSO-d_6_): δ = 13.95 - 13.23 (m, 1H), 10.09 - 9.99 (m, 1H), 8.33 - 8.08 (m, 2H), 7.47 - 6.92 (m, 2H), 4.36 - 4.21 (m, 4H), 3.94 (s, 4H), 3.83 (s, 4H), 2.94 - 2.87 (m, 3H).

#### Preparation of 7 - tert-butyl (3-(4-(methylamino)-2-(4-(pyridin-4-yl)piperazin-1-yl)-5,7-dihydro-6H-pyrrolo[3,4-d]pyrimidin-6-yl)-3-oxopropyl)carbamate (1-7a)

To a solution of 3-(tert-butoxycarbonylamino)propanoic acid (1-6a, 45.3 mg, 239 umol, 1.00 eq) in dimethyl formamide (1.0 mL) was added N,N-diisopropylethylamine (92.8 mg, 718 umol, 125 uL, 3.00 eq), hydroxybenzotriazole (48.5 mg, 359 umol, 1.50 eq) and 1-(3-dimethylaminopropyl)-3-ethylcarbodiimide hydrochloride (1-5, 68.8 mg, 359 umol, 1.50 eq). The mixture was stirred at 25 °C for 2 min. N-methyl-2-(4-(pyridin-4-yl)piperazin-1-yl)-6,7-dihydro-5H-pyrrolo[3,4-d]pyrimidin-4-amine (100 mg, 287.48 umol, 1.2 eq, hydrochloric acid) was added in the mixture stirred at 25°C for 20 min. The reaction mixture was diluted with water 30.0 mL and extracted with n-butyl alcohol (30.0 mL * 3). The combined organic layers were dried over sodium sulfate, filtered and concentrated under reduced pressure to give a residue. Tert-butyl(3-(4-(methylamino)-2-(4-(pyridin-4-yl)piperazin-1-yl)-5,7-dihydro-6H-pyrrolo[3,4-d] pyrimidin-6-yl)-3-oxopropyl)carbamate (1-7a, 250 mg, crude) was obtained as a black solid. LC-MS: R_t_ = 0.274 min, [M+H]^+^ = 483.3.

#### Preparation of 3-amino-1-(4-(methylamino)-2-(4-(pyridin-4-yl)piperazin-1-yl)-5,7-dihydro-6H-pyrrolo[3,4-d]pyrimidin-6-yl)propan-1-one (1a)

To a solution of tert-butyl (3-(4-(methylamino)-2-(4-(pyridin-4-yl)piperazin-1-yl)-5,7-dihydro-6H-pyrrolo [3,4-d]pyrimidin-6-yl)-3-oxopropyl)carbamate (1-7a, 150 mg, 310 umol, 1.00 eq) in dichloromethane (2.0 mL) was added hydrochloric acid/dioxane (4 M, 1.50 mL, 19.3 eq). The mixture was stirred at 25°C for 1 h. The reaction mixture was concentrated under reduced pressure to give a residue. The residue was purified by prep-HPLC (column: Welch Xtimate C18 150*25mm*5um;mobile phase: [water(trifluoroacetic acid)-acetonitrile];B%: 0%-20%,10min). 3-amino-1-(4-(methylamino)-2-(4-(pyridin-4-yl)piperazin-1-yl)-5,7-dihydro-6H-pyrrolo[3,4-d]pyrimidin-6-yl)propan-1-one (1a, 16.2 mg, 32.3 umol, 10.3% yield, 99% purity, trifluoroacetic acid) was obtained as a white solid. H NMR: (400 MHz, MeOD-d_4_):, δ = 8.29 - 8.15 (m, 2H), 7.19 (d, J = 7.6 Hz, 2H), 4.80 (s, 1H), 4.68 (s, 1H), 4.62 (s, 1H), 4.50 (s, 1H), 4.12 - 4.05 (m, 4H), 4.01 - 3.94 (m, 4H), 3.28 (t, J = 6.0 Hz, 2H), 3.09 (d, J = 4.4 Hz, 3H), 2.83 (td, J = 6.4, 8.8 Hz, 2H). HPLC: 99% purity. MS: [M+H]^+^ = 383.1.

#### Synthesis of 2-[4-(aminomethyl)phenyl]-1-[4-(methylamino)-2-[4-(4-pyridyl)piperazin-1-yl]-5,7-dihydropyrrolo[3,4-d]pyrimidin-6-yl]ethanone (1b)

White solid, 16.18% yield. H NMR (400 MHz, DMSO-d_6_): δ = 8.36 (s, 1H), 8.18 (d, J = 6.4 Hz, 2H), 7.38 - 7.31 (m, 2H), 7.30 - 7.25 (m, 2H), 7.05 (t, J = 4.8 Hz, 1H), 6.85 (d, J = 6.4 Hz, 2H), 4.56 (s, 2H), 4.31 (d, J = 7.2 Hz, 2H), 3.89 (s, 2H), 3.80 (s, 4H), 3.71 (d, J = 7.2 Hz, 2H), 3.43 - 3.34 (m, 6H), 2.85 (dd, J = 4.4, 8.4 Hz, 3H). HPLC: 96.22% purity. MS:[M+H]^+^ = 459.4.

#### Synthesis of (R)-3-amino-1-(4-(methylamino)-2-(4-(pyridin-4-yl)piperazin-1-yl)-5,7-dihydro-6H-pyrrolo[3,4-d]pyrimidin-6-yl)-4-(naphthalen-2-yl)butan-1-one (1c)

White solid, 8.95% yield. H NMR (400 MHz, DMSO-d_6_): δ = 13.95 - 13.66 (m, 1H), 8.36 - 8.20 (m, 5H), 7.93 - 7.82 (m, 3H), 7.79 (d, J = 6.4 Hz, 1H), 7.51 - 7.42 (m, 3H), 7.22 ( d, J = 7.6 Hz, 2H), 4.65 ( s, 1H), 4.55 - 4.34 (m, 2H), 4.32 - 4.25 (m, 1H), 4.21 - 4.12 (m, 1H), 4.00 ( s, 4H), 3.87 ( s, 5H), 3.07 (dd, J = 13.6, 8.4 Hz, 1H), 2.92 (d, J = 3.6 Hz, 3H), 2.79 - 2.69 (m, 1H), 2.69 - 2.60 (m, 1H). HPLC: 98% purity. MS: [M+H]^+^ = 523.1.

#### Synthesis of ((4R,5S)-4-Amino-5-methyl-1-(4-(methylamino)-2-(4-(pyridin-4-yl)piperazin-1-yl)-5,7-dihydro-6H-pyrrolo[3,4-d]pyrimidin-6-yl)heptan-1-one (1d)

White solid, 72.8% yield. H NMR (400 MHz, DMSO-d_6_): δ 13.78 (s, 1H), 8.34 - 8.21 (m, 2H), 8.08 (s, 3H), 7.25 - 7.10 (d, J = 7.2 Hz, 2H), 4.87 - 4.64 (m, 1H), 4.63 - 4.43 (m, 2H), 4.29 (s, 1H), 3.97 (s, 4H), 3.83 (br s, 4H), 3.09 - 3.00 (m, 1H), 2.95 - 2.85 (m, 3H), 2.57 - 2.51 (m, 2H), 1.86 - 1.75 (m, 1H), 1.71 - 1.51 (m, 2H), 1.49 - 1.28 (m, 1H), 1.24 - 1.09 (m, 1H), 0.92 - 0.72 (m, 6H). HPLC: 98% purity. MS: [M+H]^+^ = 453.3.

#### Synthesis of (R)-3-amino-1-(4-(methylamino)-2-(4-(pyridin-4-yl)piperazin-1-yl)-5,7-dihydro-6H-pyrrolo[3,4-d]pyrimidin-6-yl)-5-(methylthio)pentan-1-one (1e)

Yellow gum, 16.04% yield. H NMR (400 MHz, DMSO-d_6_): δ = 13.78 ( s, 1H), 8.30 (d, J = 5.6 Hz, 2H), 8.22 - 8.07 (m, 3H), 7.28 - 7.16 (m, 2H), 4.83 - 4.69 (m, 1H), 4.56 ( s, 2H), 4.38 ( s, 1H), 4.01 ( s, 4H), 3.87 ( s, 4H), 2.94 ( t, J = 3.6 Hz, 3H), 2.86 - 2.65 (m, 3H), 2.60 (br t, J = 7.4 Hz, 2H), 2.06 (s, 3H), 2.00 - 1.85 (m, 2H). HPLC: 96% purity. MS: [M+H]^+^ = 457.1. SFC: 1.999min, 2.117min.

#### Synthesis of 2-amino-N-(2-((2-(4-(methylamino)-2-(4-(pyridin-4-yl)piperazin-1-yl)-5,7-dihydro-6H-pyrrolo[3,4-d]pyrimidin-6-yl)-2-oxoethyl)amino)-2-oxoethyl)acetamide (1f)

White solid, 69.24% yield, HPLC: 94.337% purity. MS: [M+H]^+^ = 483.26.

#### Synthesis of (R)-4-Amino-6-methyl-1-(4-(methylamino)-2-(4-(pyridin-4-yl)piperazin-1-yl)-5,7-dihydro-6H-pyrrolo[3,4-d]pyrimidin-6-yl)heptan-1-one (1g)

Brown solid, 98.2% yield. H NMR (400 MHz, DMSO-d_6_): δ 13.99 (s, 1H), 9.19 - 8.51 (m, 1H), 8.45 - 8.22 (d, J = 5.6 Hz, 2H), 8.14 (s, 2H), 7.39 - 7.05 ( d, J = 6.8 Hz, 2H), 4.98 - 4.70 (m, 1H), 4.69 - 4.68 (m, 2H), 4.40 - 4.31 (d, J = 14.4 Hz, 1H), 4.05 (s, 4H), 3.89 (s, 5H), 3.17 (s, 1H), 2.95 (s, 3H), 2.74 - 2.58 (m, 1H), 1.98 - 1.65 (m, 3H), 1.57 - 1.32 (m, 2H), 1.05 - 0.65 (d, J = 5.6 Hz, 6H). HPLC: 99.1% purity. MS: [M+H]^+^ = 453.3.

#### Synthesis of 2-((2,4-dimethoxybenzyl)amino)-1-(4-(methylamino)-2-(4-(pyridin-4-yl)piperazin-1-yl)-5,7-dihydro-6H-pyrrolo[3,4-d]pyrimidin-6-yl)ethan-1-one (1h)

White oil, 42.63% yield. HPLC: 99.481% purity. MS: [M+H]^+^ = 519.28.

#### Synthesis of (8,8-dioxido-8-thia-2-azaspiro[4.5]decan-4-yl)(4-(methylamino)-2-(4-(pyridin-4-yl)piperazin-1-yl)-5,7-dihydro-6H-pyrrolo[3,4-d]pyrimidin-6-yl)methanone (1i)

White solid, 91.30% yield. HPLC: 94.952% purity. MS: [M+H]^+^ = 527.25.

The general procedure for the synthesis of compounds 2a to 2i is showed in scheme S2.

**Scheme S2.**
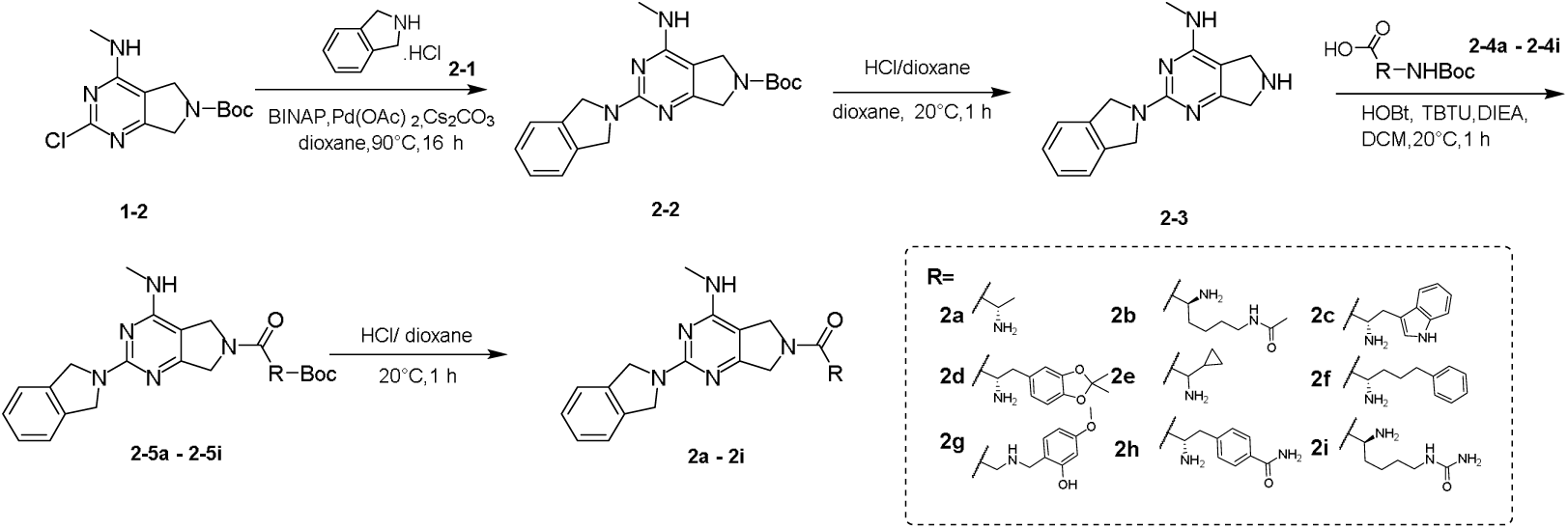
General procedures for the synthesis of compounds 2a to 2i.

#### Synthesis of (S)-2-amino-1-(2-(isoindolin-2-yl)-4-(methylamino)-5,7-dihydro-6H-pyrrolo[3,4-d]pyrimidin-6-yl)propan-1-one (2a): Preparation of tert-butyl 2-(isoindolin-2-yl)-4-(methylamino)-5,7-dihydro-6H-pyrrolo[3,4-d]pyrimidine-6-carboxylate (2-2)

A mixture of tert-butyl 2-chloro-4-(methylamino)-5,7-dihydro-6H-pyrrolo[3,4-d]pyrimidine-6-carboxylate (1-2, 2.50 g, 8.78 mmol, 1.00 eq), isoindoline (2-1, 1.37 g, 8.78 mmol, 1.30 mL, 1.00 eq, hydrochloride), [1-(2-diphenylphosphanyl-1-naphthyl)-2-naphthyl]-diphenyl-phosphane (1.09 g, 1.76 mmol, 0.200 eq), diacetoxypalladium (197 mg, 877 umol, 0.100 eq) and cesium carbonate (8.58 g, 26.3 mmol, 3.00 eq) in dioxane (50.0 mL) was degassed and purged with nitrogen for 3 times, and then the mixture was stirred at 90°C for 3 h under nitrogen atmosphere. The reaction mixture was diluted with water 200 mL and extracted with ethyl acetate 600.0 mL (200.0 mL * 3). The combined organic layers were washed with brine 200.0 mL, dried over sodium sulfate, filtered and concentrated under reduced pressure to give a residue. The residue was purified by flash silica gel chromatography. Compound tert-butyl2-(isoindolin-2-yl)-4-(methylamino)-5,7-dihydro-6H-pyrrolo[3,4-d]pyrimidine-6-carboxylate(2-2, 0.700 g, 1.91 mmol, 21.7% yield) was obtained as a gray solid. HNMR (400 MHz, DMSO-d_6_): δ = 7.42 - 7.35 (m, 4H), 7.29 - 7.24 (m, 5H), 4.77 (s, 4H), 4.32 - 4.22 (m, 4H), 2.88 (d, J = 5.2 Hz, 3H), 1.45 (d, J = 2.8 Hz, 9H).

#### Preparation of 2-(isoindolin-2-yl)-N-methyl-6,7-dihydro-5H-pyrrolo[3,4-d]pyrimidin-4-amine (2-3)

To a solution of tert-butyl 2-(isoindolin-2-yl)-4-(methylamino)-5,7-dihydro-6H-pyrrolo[3,4-d] pyrimidine-6-carboxylate (2-3, 0.300 g, 816 umol, 1.00 eq) in dioxane (2.00 mL) was added hydrochloric acid (4 M in dioxane) (4 M, 5.0 mL, 24.5 eq). The mixture was stirred at 25°C for 16 h. The reaction mixture was concentrated under reduced pressure to give a residue. 2-(isoindolin-2-yl)-N-methyl-6,7-dihydro-5H-pyrrolo[3,4-d]pyrimidin-4-amine (337 mg, crude) was obtained as a gray solid. HNMR (400 MHz, DMSO-d_6_): δ =10.42 (s, 2H), 9.14 - 8.82 (m, 1H), 7.43 (d, J = 1.6 Hz, 2H), 7.36 (dd, J = 3.2, 5.6 Hz, 2H), 4.93 (s, 4H), 4.47 (s, 2H), 4.28 (s, 2H), 2.99 (d, J = 3.6 Hz, 3H).

#### Preparation of tert-butyl (S)-(1-(2-(isoindolin-2-yl)-4-(methylamino)-5,7-dihydro-6H-pyrrolo[3,4-d]pyrimidin-6-yl)-1-oxopropan-2-yl)carbamate (2-5a)

A mixture of 2-(isoindolin-2-yl)-N-methyl-6,7-dihydro-5H-pyrrolo[3,4-d]pyrimidin-4-amine (2-3, 150 mg, 561 umol, 1.00 eq), (2S)-2-(tert-butoxycarbonylamino)propanoic acid (2-4a, 106 mg, 561 umol, 1.00 eq), N,N-diisopropylethylamine (217 mg, 1.68 mmol, 293 uL, 3.00 eq), [benzotriazol-1-yloxy(dimethylamino) methylidene]-dimethylazanium;hexafluorophosphate (255 mg, 673 umol, 1.20 eq) and 1-hydroxybenzotriazole (151 mg, 1.12 mmol, 2.00 eq) in dichloromethane (5.0 mL) was degassed and purged with nitrogen for 3 times, and then the mixture was stirred at 25°C for 1 h under nitrogen atmosphere. The reaction mixture was partitioned between water 20.0 mL and ethyl acetate 20.0 mL and extracted with ethyl acetate 60.0 mL (20.0 mL * 3). The organic phase was separated, washed with brine 20.0 mL, dried over sodium sulfate, filtered and concentrated under reduced pressure to give a residue. The residue was purified by flash silica gel chromatography. Tert-butyl (S)-(1-(2-(isoindolin-2-yl)-4-(methylamino)-5,7-dihydro-6H-pyrrolo[3,4-d]pyrimidin-6-yl)-1-oxopropan-2-yl)carbamate (120 mg, 273 umol, 48.7% yield) was obtained as a white solid. H NMR (400 MHz, DMSO-d_6_): δ =7.40 - 7.35 (m, 2H), 7.31 - 7.23 (m, 3H), 7.07 (t, J = 7.6 Hz, 1H), 6.98 (s, 1H), 4.77 (s, 5H), 4.43 - 4.13 (m, 4H), 2.90 (t, J = 4.8 Hz, 3H), 1.45 - 1.41 (m, 3H), 1.38 (s, 9H).

#### Preparation of compound-(S)-2-amino-1-(2-(isoindolin-2-yl)-4-(methylamino)-5,7-dihydro-6H-pyrrolo[3,4-d]pyrimidin-6-yl)propan-1-one (2a)

To a solution of tert-butyl (S)-(1-(2-(isoindolin-2-yl)-4-(methylamino)-5,7-dihydro-6H-pyrrolo[3,4-d] pyrimidin-6-yl)-1-oxopropan-2-yl)carbamate (2-5a, 120 mg, 273 umol, 1.00 eq) in dioxane (1.00 mL) was added HCl/dioxane (4 M, 2.0 mL, 29.2 eq). The mixture was stirred at 25°C for 1 h. The reaction mixture was concentrated under reduced pressure to give a residue. The residue was purified by prep-HPLC (column: Phenomenex luna C18 150*25mm* 10um;mobile phase: [water(FA)-ACN];B%: 1%-28%,15min). (S)-2-amino-1-(2-(isoindolin-2-yl)-4-(methylamino)-5,7-dihydro-6H-pyrrolo[3,4-d]pyrimidin-6-yl)propan-1-one (11.74 mg, 33.14 umol, 12.11% yield, 95.533% purity) was obtained as a off-white solid. H NMR (400 MHz, DMSO-d_6_): δ=8.32 (s, 1H), 7.43 - 7.35 (m, 2H), 7.33 - 7.25 (m, 2H), 7.08 - 6.92 (m, 1H), 4.91 - 4.62 (m, 5H), 4.58 - 4.47 (m, 1H), 4.43 - 4.25 (m, 2H), 3.83 - 3.62 (m, 3H), 2.94 - 2.88 (m, 3H), 1.26 - 1.17 (m, 3H). HPLC: 95.533% purity. MS: [M+H]^+^ = 339.3. SFC: R_t_ = 1.853 min.

#### Synthesis of (S)-N-(5-amino-6-(2-(isoindolin-2-yl)-4-(methylamino)-5,7-dihydro-6H-pyrrolo[3,4-d] pyrimidin-6-yl)-6-oxohexyl)acetamide (2b)

White solid, 46.71% yield. H NMR (400 MHz, DMSO-d_6_): δ = 9.40 - 8.84 (m, 1H), 8.48 (s, 3H), 8.08 - 7.96 (m, 1H), 7.47 - 7.32 (m, 4H), 5.14 - 4.58 (m, 7H), 4.52 - 4.33 (m, 1H), 4.15 - 3.93 (m, 1H), 3.03 ( d, J = 5.2 Hz, 2H), 2.96 ( s, 1H), 2.99 ( d, J = 4.4 Hz, 2H), 1.88 - 1.73 (m, 5H), 1.43 ( d, J = 6.4 Hz, 4H). MS: [M+H]^+^ = 438.3. SFC: Rt = 1.521,1.558 min.

#### Synthesis of (S)-2-amino-3-(1H-indol-3-yl)-1-(2-(isoindolin-2-yl)-4 - (methylamino)-5,7-dihydro-6H-pyrrolo[3,4-d]pyrimidin-6-yl)propan-1-one (2c)

Yellow solid,12.31% yield. H NMR (400 MHz, DMSO-d_6_): δ = 11.12 ( s, 1H), 8.94 - 8.41 (m, 4H), 7.56 (dd, J = 4.4, 7.2 Hz, 1H), 7.47 - 7.25 (m, 6H), 7.08 - 6.97 (m, 2H), 4.96(s, 4H), 4.71 - 4.57 (m, 1H), 4.40 ( d, J = 17.6 Hz, 1H), 4.32 - 3.91 (m, 3H), 3.29 ( dd, J = 6.4, 10.4 Hz, 2H), 3.01 - 2.88 (m, 3H). MS: [M+H]^+^ = 454.1. SFC: Rt = 1.812,1.982 min.

#### Synthesis of (S)-2-amino-3-(2,2-dimethylbenzo[d][1,3]dioxol-5-yl)-1-(2-(isoindolin-2-yl)-4-(methylamino)-5,7-dihydro-6H-pyrrolo[3,4-d]pyrimidin-6-yl)propan-1-one (2d)

White solid, 10.00% yield. H NMR (400 MHz, DMSO-d_6_): δ = 7.42 - 7.34 (m, 2H), 7.32 - 7.26 (m, 2H), 6.96 ( dd, J = 4.8, 11.6 Hz, 1H), 6.77 - 6.61 (m, 3H), 4.87 - 4.67 (m, 4H), 4.61 - 4.50 (m, 1H), 4.37 - 4.22 (m, 2H), 4.22 - 3.94 (m, 1H), 3.75 - 3.53 (m, 1H), 2.90 (dd, J = 3.2, 4.4 Hz, 3H), 2.80 - 2.66 (m, 1H), 2.64 - 2.51 (m, 2H), 1.64 - 1.46 (m, 7H). MS: [M+H]^+^ = 487.2. SFC: Rt = 1.163,1.367 min.

#### Synthesis of 2-amino-2-cyclopropyl-1-[2-isoindolin-2-yl-4-(methylamino)-5,7-dihydropyrrolo[3,4-d]pyrimidin-6-yl]ethanone (2e)

White solid, 23.00% yield. MS: [M+H]^+^ = 365.

#### Synthesis of (S)-2-Amino-1-(2-(isoindolin-2-yl)-4-(methylamino)-5,7-dihydro-6H-pyrrolo[3,4-d]pyrimidin-6-yl)-5-phenylpentan-1-one (2f)

White solid, 10.1% yield. H NMR (400 MHz, MeOD-d_4_): δ 7.49 - 7.33 (m, 4H), 7.30 - 7.10 (m, 5H), 4.91 (s, 4H), 4.81 - 4.75 (m, 2H), 4.72 - 4.58 (m, 2H), 4.55 - 4.41 (m, 1H), 4.31 - 4.20 (m, 1H), 3.15 - 3.11 (d, J = 9.2 Hz, 3H), 2.82 - 2.59 (m, 2H), 2.01 - 1.90 (m, 1H), 1.88 - 1.76 (m, 2H). MS: [M+H]^+^ = 443.3.

#### Synthesis of 2-amino-2-cyclopropyl-1-[2-isoindolin-2-yl-4-(methylamino)-5,7-dihydropyrrolo[3,4-d]pyrimidin-6-yl]ethanone (2g)

White solid, 11.69% yield. MS: [M+H]^+^ = 460.53.

#### Synthesis of (S)-4-(2-Amino-3-(2-(isoindolin-2-yl)-4-(methylamino)-5,7-dihydro-6H-pyrrolo[3,4-d]pyrimidin-6-yl)-3-oxopropyl)benzamide (2h)

Yellow solid, 80.9% yield. ^1^H NMR (400 MHz, DMSO-d_6_): δ 7.97 - 7.75 (m, 3H), 7.44 - 7.23 (m, 7H), 7.08 - 6.94 (m, 1H), 4.87 - 4.61 (m, 4H), 4.46 - 4.20 (m, 2H), 4.19 - 4.00 (m, 1H), 3.08 - 3.02 (m, 1H), 2.92 - 2.80 (d, J = 4.4 Hz, 3H), 1.90 (s, 4H). HPLC: 93.0% purity. MS: [M+H]^+^ = 458.3.

#### Synthesis of (S)-1-(5-Amino-6-(2-(isoindolin-2-yl)-4-(methylamino)-5,7-dihydro-6H-pyrrolo[3,4-d]pyrimidin-6-yl)-6-oxohexyl)urea (2i)

White solid, 15.93% yield. H NMR (400 MHz, MeOH-d_4_): δ 7.36 - 7.23 (m, 4H), 4.81 - 4.42 (m, 8H), 3.76 - 3.65 (m, 1H), 3.15 – 3.01 (m, 5H), 1.77 - 1.70 (m, 1H), 1.64 - 1.39 (m, 5H). HPLC: 90.9% purity. MS: [M+H]^+^ = 439.2.

The general procedure for the synthesis of compounds 3a and 3b is showed in scheme S3.

**Scheme S3.**
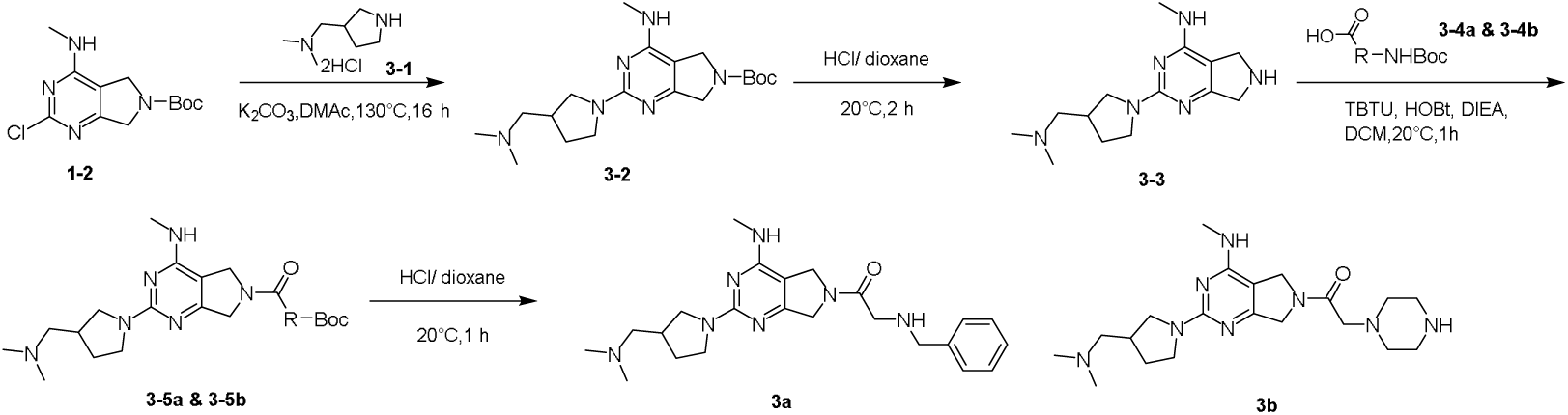
General procedures for the synthesis of compounds 3a and 3b.

#### Preparation of compound 3 - tert-butyl 2-(3-((dimethylamino)methyl)pyrrolidin-1-yl)-4-(methylamino)-5,7-dihydro-6H-pyrrolo[3,4-d]pyrimidine-6-carboxylate (3-2)

To a solution of tert-butyl 2-chloro-4-(methylamino)-5,7-dihydro-6H-pyrrolo[3,4-d] pyrimidine-6-carboxylate (1-2, 900 mg, 3.16 mmol, 1.00 eq) in N,N-dimethylacetamide (10.0 mL) was added potassium carbonate (1.75 g, 12.6 mmol, 4.00 eq) and N,N-dimethyl-1-pyrrolidin-3-yl-methanamine (3-1, 635 mg, 3.16 mmol, 1.00 eq, 2 hydrochloric). The mixture was stirred at 130°C for 16 h. The reaction mixture was poured into water (20.0 mL) and extracted with dichloromethane: isopropanol (5:1) 60 mL (20.0 mL * 3). The combined organic layers were washed with brine 20 mL, dried over sodium sulfate, filtered and concentrated under reduced pressure to give a residue. Tert-butyl 2-(3-((dimethylamino)methyl)pyrrolidin-1-yl)-4-(methylamino)-5,7-dihydro-6H-pyrrolo[3,4-d]pyrimidine-6-carboxylate (3-2, 1.10 g, 2.92 mmol, 92.4% yield) was obtained as a white solid. H NMR (400 MHz, DMSO-d_6_): δ= 6.79 (quin, J = 4.8 Hz, 1H), 4.30 - 4.11 (m, 3H), 3.71 - 3.45 (m, 2H), 3.16 - 3.04 (m, 1H), 2.94 (s, 4H), 2.84 - 2.76 (m, 6H), 2.38 (td, J = 7.2, 14.4 Hz, 1H), 2.28 - 2.17 (m, 2H), 2.14 (s, 5H), 2.02 - 1.86 (m, 5H), 1.64 - 1.52 (m, 1H), 1.44 (d, J = 5.2 Hz, 7H).

#### Preparation of compound-2-(3-((dimethylamino)methyl)pyrrolidin-1-yl)-N-methyl-6,7-dihydro-5H-pyrrolo[3,4-d]pyrimidin-4-amine (3-3)

To a solution of tert-butyl 2-(3-((dimethylamino)methyl)pyrrolidin-1-yl)-4-(methylamino)-5,7-dihydro-6H-pyrrolo[3,4-d]pyrimidine-6-carboxylate (3-2, 1.10 g, 2.92 mmol, 1.00 eq) in dioxane (10.0 mL) was added hydrochloric/dioxane (4 M, 10.0 mL, 13.69 eq). The mixture was stirred at 25°C for 1 h. The reaction mixture was concentrated under reduced pressure to give a residue. 2-(3-((dimethylamino)methyl)pyrrolidin-1-yl)-N-methyl-6,7-dihydro-5H-pyrrolo[3,4-d]pyrimidin-4-amine (3-3, 0.800 g, 2.89 mmol, 99.1% yield) was obtained as a gray solid.MS: [M+H]^+^ = 277.1.

#### Preparation of compound - tert-butyl benzyl(2-(2-(3-((dimethylamino)methyl)pyrrolidin-1-yl)-4-(methylamino)-5,7-dihydro-6H-pyrrolo[3,4-d]pyrimidin-6-yl)-2-oxoethyl)carbamate (3-5a)

A mixture of 2-(3-((dimethylamino)methyl)pyrrolidin-1-yl)-N-methyl-6,7-dihydro-5H-pyrrolo[3,4-d] pyrimidin-4-amine (3-3, 0.2 g, 723.64 umol, 1.00 eq), N-benzyl-N-(tert-butoxycarbonyl)glycine (3-4a, 191 mg, 724 umol, 1.00 eq), N,N-diisopropylethylamine (467 mg, 3.62 mmol, 630 uL, 5.00 eq), [benzotriazol-1-yloxy(dimethylamino)methylidene]-dimethylazanium;hexafluorophosphate (329 mg, 868 umol, 1.20 eq) and 1-hydroxybenzotriazole (195 mg, 1.45 mmol, 2.00 eq) in dichloromethane (6.0 mL) was degassed and purged with nitrogen for 3 times, and then the mixture was stirred at 25°C for 1 h under nitrogen _2_ atmosphere. The reaction mixture was diluted with water 20.0 mL and extracted with dichloromethane 60.0 mL (20.0 mL * 3). The combined organic layers were washed with brine 20 mL, dried over sodium sulfate, filtered and concentrated under reduced pressure to give a residue. The residue was purified by flash silica gel chromatography. Tert-butyl benzyl(2-(2-(3-((dimethylamino)methyl)pyrrolidin-1-yl)-4-(methylamino)-5,7-dihydro-6H-pyrrolo[3,4-d]pyrimidin-6-yl)-2-oxoethyl)carbamate (0.225 g, 429.66 umol, 59.37% yield) was obtained as a white solid. H NMR (400 MHz, DMSO-d_6_): δ= 7.37 - 7.30 (m, 3H), 7.30 - 7.23 (m, 5H), 4.46 - 4.36 (m, 5H), 4.30 ( d, J = 18.0 Hz, 2H), 3.90 ( d, J = 8.0 Hz, 1H), 3.67 - 3.60 (m, 1H), 3.56 - 3.52 (m, 1H), 3.14 - 3.06 (m, 2H), 3.04 - 2.94 (m, 1H), 2.86 - 2.77 (m, 3H), 2.44 - 2.37 (m, 2H), 2.27 (s, 6H), 1.35 (s, 9H), 1.21 (d, J = 6.4 Hz, 4H).

#### Preparation of 2-(benzylamino)-1-(2-(3-((dimethylamino)methyl)pyrrolidin-1-yl)-4-(methylamino)-5,7-dihydro-6H-pyrrolo[3,4-d]pyrimidin-6-yl)ethan-1-one (3a)

To a solution of tert-butylbenzyl(2-(2-(3-((dimethylamino)methyl)pyrrolidin-1-yl)-4-(methylamino)-5,7-dihydro-6H-pyrrolo[3,4-d]pyrimidin-6-yl)-2-oxoethyl)carbamate (3-5a, 0.200 g, 381 umol, 1.00 eq) in methanol (1.00 mL) was added hydrochloric acid (4 M in dioxane) (4 M, 2.0 mL, 20.9 eq). The mixture was stirred at 20°C for 1 h. The reaction mixture was concentrated under reduced pressure to give a residue. The residue was purified by prep-HPLC (column: Welch Xtimate C18 150*25mm*5um;mobile phase: [water(TFA)-ACN];B%: 0%-26%,10min). 2-(benzylamino)-1-(2-(3-((dimethylamino)methyl)pyrrolidin-1-yl)-4-(methylamino)-5,7-dihydro-6H-pyrrolo[3,4-d]pyrimidin-6-yl)ethan-1-one (85.23 mg, 185.40 umol, 48.54% yield, 92.134% purity) was obtained as a red gum. H NMR (400 MHz, DMSO-d_6_): δ=9.89 - 9.40 (m, 2H), 7.55 - 7.40 (m, 5H), 4.68 - 4.33 (m, 3H), 4.24 - 3.97 (m, 5H), 3.84 (d, J = 12.4 Hz, 2H), 3.74 (s, 2H), 3.54 - 3.37 (m, 2H), 3.34 - 3.12 (m, 3H), 2.90 (d, J = 4.4 Hz, 2H), 2.83 (s, 4H), 2.77 - 2.63 (m, 1H), 2.27 - 2.06 (m, 1H), 1.88 - 1.61 (m, 1H). MS: [M+H]^+^ = 424.1.

#### Synthesis of 1-[2-[3-[(dimethylamino)methyl]pyrrolidin-1-yl]-4-(methylamino)-5,7-dihydropyrrolo[3,4-d]pyrimidin-6-yl]-2-piperazin-1-yl-ethanone (3b)

Yellow gum, 18.79% yield. ^1^H NMR (400 MHz, DMSO-d_6_): δ = 10.08 - 9.50 (m, 1H), 9.15 - 7.84 (m, 3H), 4.77 - 4.52 (m, 2H), 4.50 - 4.32 (m, 2H), 3.94 - 3.70 (m, 6H), 3.20 (d, J = 14.8 Hz, 6H), 2.93 (dd, J = 4.4, 8.0 Hz, 6H), 2.83 (s, 5H), 2.77 - 2.67 (m, 1H), 2.21 (s, 1H), 1.93 - 1.67 (m, 1H). MS: [M+H]^+^ = 402.29.

#### Preparation of methyl 3-(4-bromophenyl)sulfanyl-2-nitro-benzoate

To a solution of methyl 3-fluoro-2-nitro-benzoate (2.00 g, 10.04 mmol, 1.00 eq) and 4-bromobenzenethiol (1.90 g, 10.04 mmol, 1.00 eq) in dimethylformamide (1.0 mL) was added N,N-diisopropylethylamine (3.89 g, 30.13 mmol, 5.3 mL, 3.00 eq). The mixture was stirred at 90 °C for 3 h. The mixture solution was diluted with water 50.0 mL and extracted with ethyl acetate 100.0 mL. And then the organic phase was washed with saturated sodium chloride 50.0 mL, dried over sodium sulfate and filtered and concentrated under reduced pressure to give a residue. The residue was purified by flash silica gel chromatography (ISCO®; 40 g SepaFlash® Silica Flash Column, Eluent of 100% Ethyl acetate ethergradient @ 60 mL/min). Methyl 3-(4-bromophenyl)sulfanyl-2-nitro-benzoate (1.27 g, 3.45 mmol, 34.34% yield) was obtained as a yellow solid. H NMR (400 MHz, DMSO-d_6_): δ = 3.86 (s, 3 H) 7.33 - 7.40 (m, 2 H) 7.59 - 7.66 (m, 2 H) 7.59 - 7.67 (m, 1 H) 7.67 - 7.77 (m, 1 H) 7.95 (dd, J=7.60, 1.19 Hz, 1 H).

#### Preparation of 3-(4-bromophenyl)sulfanyl-2-nitro-benzoic acid

To a solution of methyl 3-(4-bromophenyl)sulfanyl-2-nitro-benzoate (1.20 g, 3.26 mmol, 1.00 eq) in methanol (15.0 mL) and water (8.0 mL) was added Lithium hydroxide hydrate (684 mg, 16.30 mmol, 5.00 eq). The mixture was stirred at 25 °C for 3 h. The mixture solution was adjusted to pH = 2∼3 and extracted with ethyl acetate 150.0 mL. Then the organic phase was washed with saturated sodium chloride 200.0 mL, dried over sodium sulfate, filtered and concentrated under reduced pressure to give a residue. 3-(4-bromophenyl)sulfanyl-2-nitro-benzoic acid (1.10 g, 3.11 mmol, 95.30% yield) was obtained as a yellow solid. LCMS: R_t_ = 0.474 min, [M+H]^+^ = 493.4.

#### Preparation of 3-(4-bromophenyl)sulfanyl-N-methyl-2-nitro-benzamide

To a solution of 3-(4-bromophenyl)sulfanyl-2-nitro-benzoic acid (580 mg, 1.64 mmol, 1.00 eq) and methanamine;hydrochloride in dimethylformamide (12.0 mL) was added O-(7-azabenzotriazol-1-yl)-N,N,N,N-tetramethyluroniumhexafluorophosphate (934 mg, 2.46 mmol, 1.50 eq) and N,N-diisopropylethylamine (423 mg, 3.28 mmol, 570.48 μL, 2.00 eq). The mixture was stirred at 25 °C for 3 h. Ethyl acetate (30.0 mL) and water (30.0 mL) were added and layers were separated. The aqueous phase was extracted with ethyl acetate (30.0 mL x 2). Combined extracts werewashed with brine (60.0 mL), dried over magnesium sulfate, filtered, and concentrated under vacuum. The residue was purified by column chromatography (SiO_2_, Petroleum ether/Ethyl acetate=5/1). 3-(4-bromophenyl)sulfanyl-N-methyl-2-nitro-benzamide (485 mg, 1.32 mmol, 80.65% yield) was obtained as a yellow solid. LCMS: R_t_ = 0.413 min, [M+H]^+^ = 366.9.

Preparation of 3-[4-(4-tert-butoxyphenyl)phenyl]sulfanyl-N-methyl-2-nitro-benzamide To a solution of 3-(4-bromophenyl)sulfanyl-N-methyl-2-nitro-benzamide (330 mg, 898.65 μmol, 1.00 eq) and (4-tert-butoxyphenyl)boronic acid (174 mg, 898.65 μmol, 1.00 eq) in dioxane (3.0 mL) and H2O (0.6 mL) was added Methanesulfonato(2-dicyclohexylphosphino-2,4,6-tri-i-propyl-1,1-biphenyl) (2-amino-1,1-biphenyl-2-yl)palladium(ii) (761 mg, 898.65 μmol, 1.00 eq) and potassium phosphate (191 mg, 898.65 μmol, 1.00 eq). The mixture was stirred at 80 °C for 2 h under N_2_. The mixture solution was diluted with water 20.0 mL and extracted with ethyl acetate 90.0 mL.

And then the organic phase was washed with saturated sodium chloride 30 mL, dried over sodium sulfate and filtered and concentrated under reduced pressure to give a residue. The residue was purified by flash silica gel chromatography (ISCO®; 40 g SepaFlash® Silica Flash Column, Eluent of 0∼50% Ethyl acetate/Petroleum ethergradient @ 60 mL/min). 3-[4-(4-tert-butoxyphenyl)phenyl]sulfanyl-N-methyl-2-nitro-benzamide (360 mg, 824.70 μmol, 91.77% yield) was obtained as an orange solid. LCMS: R_t_ = 0.532 min, [M+H]^+^ = 437.2.

#### Preparation of 2-amino-3-[4-(4-tert-butoxyphenyl)phenyl]sulfanyl-N-methyl-benzamide (7b)

To a solution of 3-[4-(4-tert-butoxyphenyl)phenyl]sulfanyl-N-methyl-2-nitro-benzamide (310 mg, 710.16 μmol, 1.00 eq) in ethanol (4.0 mL) and water (0.8 mL) was added iron (238 mg, 4.26 mmol, 6.00 eq) and ammonium chloride (2282 mg, 4.26 mmol, 6.00 eq). The mixture was stirred at 85 °C for 2 h. The residue was filtered, and diluted with water 80.0 mL and extracted with ethyl acetate 100.0 mL. And then the organic phase was washed with saturated sodium chloride 50.0 mL, dried over sodium sulfate and filtered and concentrated under reduced pressure to give a residue. The residue was purified by prep-HPLC (column: Phenomenex luna C18 150*25mm* 10um; mobile phase: [water(FA)-ACN];B%: 66%-96%,15min). 2-amino-3-[4-(4-tert-butoxyphenyl)phenyl]sulfanyl-N-methyl-benzamide (62.71 mg, 152.56 μmol, 21.48% yield, 98.903% purity was obtained as a yellow solid. H NMR (400 MHz, DMSO-d_6_): δ = 1.31 (d, J=1.20 Hz, 9 H) 2.75 (d, J=4.40 Hz, 3 H) 6.58 - 6.72 (m, 3 H) 6.98 - 7.07 (m, 2 H) 7.10 - 7.18 (m, 2 H) 7.48 - 7.66 (m, 6 H) 8.33 - 8.48 (m, 1 H). LCMS: R_t_ = 2.224 min, [M+H]^+^ = 407.2. HPLC: 98.903% purity, R_t_ = 3.368 min.

#### Synthesis of 2-amino-3-[[5-(4-tert-butoxyphenyl)-2-pyridyl]amino]-N-methyl-benzamide (7c): Preparation of 3-fluoro-2-nitro-benzoic acid

To a solution of methyl 3-fluoro-2-nitro-benzoate (1.50 g, 7.53 mmol, 1.00 eq) in tetrahydrofuran (20.0 mL) and water (6.0 mL) was added lithium monohydroxide (2 M, 3.77 mL, 1.00 eq). The mixture was stirred at 25 °C for 12 h. TLC indicated Reactant 1 was consumed completely and one new spot formed. The reaction was clean according to TLC. The reaction mixture was concentrated under reduced pressure to remove solvent. The residue was diluted with water 20 mL and adjusted the pH to 3, then extracted with dichloromethane 100 mL (50 mL * 2). The combined organic layers were dried over sodium sulfate, filtered and concentrated under reduced pressure to give a brown solid. 3-fluoro-2-nitro-benzoic acid (1.30 g, 7.02 mmol, 93.2% yield) was obtained as a brown solid and was used into the next step without further purification.

#### Preparation of 3-fluoro-N-methyl-2-nitro-benzamide

To a solution of 3-fluoro-2-nitro-benzoic acid (1.30 g, 7.02 mmol, 1.00 eq) and methanamine;hydrochloride (474 mg, 7.02 mmol, 1.00 eq) in dichloromethane (1.0 mL) was added O-(7-azabenzotriazol-1-yl)-N,N,N,N-tetramethyluroniumhexafluorophosphate (4.01 g, 10.5 mmol, 1.50 eq) and N,N-diisopropylethylamine (2.72 g, 21.07 mmol, 3.67 mL, 3.00 eq). The mixture was stirred at 25 °C for 12 h. The reaction mixture was diluted with water 50 mL and extracted with dichloromethane 300 mL (100 mL * 3). The combined organic layers were washed with water 50 mL and brine 50 mL, dried over sodium sulfate, filtered and concentrated under reduced pressure to give a residue. The residue was purified by column choursomatography (SiO_2_, dichloromethane/MeOH=100/1 to 10/1). 3-fluoro-N-methyl-2-nitro-benzamide (1.25 g, 6.31 mmol, 89.8% yield) was obtained as a white solid. LC-MS: R_t_ = 0.212 min, [M+H]^+^ = 199.2.

#### Preparation of 3-[(5-iodo-2-pyridyl)amino]-N-methyl-2-nitro-benzamide

To a solution of 5-iodopyridin-2-amine (666 mg, 3.03 mmol, 1.20 eq) in tetrahydrofuran (10.0 mL) was added sodium hydride (202 mg, 5.05 mmol, 60% purity, 2.00 eq) at 0 °C for 30 minutes. Then 3-fluoro-N-methyl-2-nitro-benzamide (500 mg, 2.52 mmol, 1 eq) was added into the mixture. The mixture was stirred at 20 °C for 12 hours. The reaction mixture was quenched by addition ammonium chloride 20 mL at 0 °C, and then diluted with water 50 mL and extracted with ethyl acetate 200 mL (50 mL * 4). The combined organic layers were washed with water 50 mL, dried over sodium sulfate, filtered and concentrated under reduced pressure to give a residue. The residue was purified by column choursomatography (SiO_2_, dichloromethane/MeOH=100/1 to 10/1). 3-[(5-iodo-2-pyridyl)amino]-N-methyl-2-nitro-benzamide (400 mg, 1.00 mmol, 39.81% yield) was obtained as a yellow solid. H NMR (400 MHz, CDCl_3_): δ = 8.85 (s, 1H), 8.48 - 8.43 (m, 2H), 7.85 (dd, J = 2.0 Hz, 8.4 Hz, 1H), 7.50 (t, J = 7.6 Hz, 1H), 7.40 - 7.36 (m, 2H), 7.05 (dd, J = 0.4 Hz, 7.2 Hz, 1H), 3.04 (d, J = 5.2 Hz, 3H). LC-MS: R_t_ = 0.382 min, [M+H]^+^ = 399.0.

#### Preparation of 3-[[5-(4-tert-butoxyphenyl)-2-pyridyl]amino]-N-methyl-2-nitro-benzamide

To a solution of 3-[(5-iodo-2-pyridyl)amino]-N-methyl-2-nitro-benzamide (350 mg, 879 μmol, 1.00 eq) and (4-tert-butoxyphenyl)boronic acid (256 mg, 1.32 mmol, 1.50 eq) in dioxane (5.0 mL) and water (1.0 mL) was added [1,1-bis(diphenylphosphino)ferrocene]dichloropalladium(II) (64 mg, 88 μmol, 0.10 eq) and sodium carbonate (186 mg, 1.76 mmol, 2.00 eq). The mixture was stirred at 90 °C for 2 h. The reaction mixture was concentrated under reduced pressure to remove solvent. The residue was diluted with water 20 mL and extracted with dichloromethane 100 mL (50 mL * 2). The combined organic layers were washed with brine 20 mL, dried over sodium sulfate, filtered and concentrated under reduced pressure to give a residue. The residue was purified by column choursomatography (SiO_2_, Ethyl acetate/Petroleum ether=1/10 to 3/1). 3-[[5-(4-tert-butoxyphenyl)-2-pyridyl]amino]-N-methyl-2-nitro-benzamide (174 mg, 414 μmol, 47.1% yield) was obtained as a white solid. H NMR (400 MHz, CDCl_3_):δ = 9.04 (s, 1H), 8.56 - 8.49 (m, 2H), 7.84 (dd, J = 2.4 Hz, 8.4 Hz, 1H), 7.53 - 7.42 (m, 3H), 7.11 - 7.06 (m, 2H), 7.02 (dd, J = 1.2 Hz, 7.6 Hz, 1H), 6.96 (d, J = 8.8 Hz, 1H), 5.98 - 5.91 (m, 1H), 3.04 (d, J = 4.8 Hz, 3H), 1.40 (s, 9H). LC-MS: R_t_ = 0.414 min, [M+H]^+^ = 421.2.

#### Preparation of 2-amino-3-[[5-(4-tert-butoxyphenyl)-2-pyridyl]amino]-N-methyl-benzamide (7c)

To a solution of 3-[[5-(4-tert-butoxyphenyl)-2-pyridyl]amino]-N-methyl-2-nitro-benzamide (70.0 mg, 166 μmol, 1.00 eq) in ethyl acetate (2.0 mL) was added Pd/C (70.0 mg, 10% purity). The mixture was degassed and purged with hydrogen for 3 times, and then the mixture was stirred at 20 °C for 2 hours under hydrogen atmosphere(14.5 psi). The mixture was filtered and washed with ethyl acetate (20mL*2) then concentrated under reduced pressure to give a residue. 2-amino-3-[[5-(4-tert-butoxyphenyl)-2-pyridyl]amino]-N-methyl-benzamide (37.76 mg, 95.98 μmol, 57.65% yield, 99.252% purity) was obtained as a white solid without purification. H NMR (400 MHz, DMSO-d_6_): δ = 8.33 (d, J = 2.2 Hz, 1H), 8.25 (d, J = 4.4 Hz, 1H), 8.05 (s, 1H), 7.78 (dd, J = 2.4, 8.4 Hz, 1H), 7.50 (d, J = 8.4 Hz, 2H), 7.40 (d, J = 6.8 Hz, 1H), 7.34 (dd, J = 0.8, 3.6 Hz, 1H), 7.02 (d, J = 8.4 Hz, 2H), 6.63 (d, J = 8.4 Hz, 1H), 6.58 (t, J = 7.6 Hz, 1H), 6.19 (s, 2H), 2.75 (d, J = 4.4 Hz, 3H), 1.31 (s, 9H). LC-MS: R_t_ = 0.350 min, [M+H]^+^ =391.1. HPLC: 99.252% purity, R_t_ = 1.491 min. (S)-4-(5-chloro-3-methylimidazo[1,2-a]pyridine-2-carbonyl)-N-methylmorpholine-3-carboxamide (8a)

#### Preparation of tert-butyl (S)-3-(methylcarbamoyl)morpholine-4-carboxylate

To a solution of (3S)-4-tert-butoxycarbonylmorpholine-3-carboxylic acid (500 mg, 2.16 mmol, 0.800 eq) and methanamine;hydrochloride (224.90 mg, 3.33 mmol, 1.20 eq) in dichloromethane (5.00 mL)was added N,N-diisopropylethylamine (1.08 g, 8.33 mmol, 1.45 mL, 3.00 eq) and added 2,4,6-tributyl-1,3,5,2,4,6 trioxatriphosphinane 2,4,6-trioxide (2.40 g, 3.33 mmol, 1.20 eq) at 0 C. The mixture was stirred at 20°C for 1 h. The solution was quenched by water (50.0 mL) and extracted with ethyl acetate 150 mL (50.0 mL*3), dried over by sodium sulfate, filtered and concentrated under reduced pressure to give a tert-butyl (3S)-3-(methylcarbamoyl)morpholine-4-carboxylate (400 mg, 1.64 mmol, 59.0% yield) was obtained as a white solid. ^1^H NMR (400 MHz, DMSO-d_6_):δ = 7.83 ( d, J = 3.2 Hz, 1H), 4.29 - 4.05 (m, 2H), 3.74 ( s, 1H), 3.61 - 3.46 (m, 2H), 3.38 - 3.08 (m, 2H), 2.61 ( d, J = 4.4 Hz, 2H), 1.54 - 1.20 (m, 9H).

#### Preparation of (S)-N-methylmorpholine-3-carboxamide

To a solution of tert-butyl (3S)-3-(methylcarbamoyl)morpholine-4-carboxylate (400 mg, 1.64 mmol, 1.00 eq) in dimethylformamide (4.00 mL) was added hydrogen chloride in dioxane (4.00 M, 4.00 mL, 9.77 eq).The mixture was stirred at 20°C for 1 h. The mixture was concentrated to give (3S)-N-methylmorpholine-3-carboxamide (300 mg, crude) was obtained as a white solid. H NMR (400 MHz, DMSO-d_6_): δ = 10.35 - 9.91 (m, 1H), 9.48 - 9.04 (m, 1H), 8.88 (s, 1H), 4.22 - 3.81 (m, 3H), 3.68 ( t, J = 10.4 Hz, 1H), 3.62 - 3.53 (m, 1H), 3.26 - 2.94 (m, 2H), 2.63 ( d, J = 3.6 Hz, 3H).

#### Preparation of (S)-4-(5-chloro-3-methylimidazo[1,2-a]pyridine-2-carbonyl)-N-methylmorpholine-3-carboxamide (8a)

To a solution of (3S)-N-methylmorpholine-3-carboxamide (51.5 mg, 285 μmol, 1.20 eq, hydrochloric) and 5-chloro-3-methyl-imidazo[1,2-a] pyridine-2-carboxylic acid (50 mg, 237 μmol, 1.00 eq) in dimethylformamide (2.50 mL) was added N,N-diisopropylethylamine (92.0 mg, 712 μmol, 124 μL, 3.00eq), O-(7-azabenzotriazol-1-yl)-N,N,N,N-tetramethyluroniumhexafluorophosphate (135 mg, 356 μmol, 1.50 eq). The mixture was stirred at 20°C for 1 h. The solution was quenched by water (10 mL) and extracted with dichloromethane: isopropanol=4:1=30mL(10 mL*3), dried over by sodium sulfate, filtered and concentrated under reduced pressure to give a residue. The residue was purified by prep-TLC (dichloromethane/methanol = 10 / 1).to give a (3S)-4-(5-chloro-3-methyl-imidazo[1,2-a] pyridine-2-carbonyl)-N-methyl-morpholine-3-carboxamide (9.66 mg, 28.0 μmol, 11.8% yield, 97.6% purity) was obtained as a yellow solid. H NMR (400 MHz, DMSO-d_6_): δ = 8.02 - 7.78 (m, 1H), 7.54 ( dd, J = 8.8, 14.8 Hz, 1H), 7.29 - 7.17 (m, 1H), 7.08 ( d, J = 7.2 Hz, 1H), 4.83 (s, 1H), 4.34 - 4.12 (m, 1H), 4.00 - 3.79 (m, 1H), 3.44 ( d, J = 10.8 Hz, 3H), 2.86 ( d, J = 9.6 Hz, 3H), 2.66 ( d, J = 4.0 Hz, 2H), 2.57 ( d, J = 4.0 Hz, 2H). LC-MS: R_t_ =0.840min, [M+H]^+^=337.1. HPLC: 97.6% purity, R_t_ =0.986min. SFC: R_t_ =1.472min.

#### Synthesis of (S)-2-(5-chloro-3-methylimidazo[1,2-a]pyridine-2-carbonyl)-7-hydroxy-N-methyl-1,2,3,4-tetrahydroisoquinoline-3-carboxamide (8b): Preparation of compound 2 - tert-butyl (S)-7-hydroxy-3-(methylcarbamoyl)-3,4-dihydroisoquinoline-2(1H)-carboxylate

To a mixture of (S)-2-(tert-butoxycarbonyl)-7-hydroxy-1,2,3,4-tetrahydroisoquinoline-3-carboxylic acid (500 mg, 1.70 mmol, 1.00 eq) and methanamine;hydrochloride (138 mg, 2.05 mmol, 1.20 eq) in dichloromethane (5 mL) were added N,N-diisopropylethylamine (661 mg, 5.11 mmol, 891 μL, 3.00 eq) and 2,4,6-tributyl-1,3,5,2,4,6trioxatriphosphinane 2,4,6-trioxide (1.47 g, 2.05 mmol, 1.20 eq), then it was stirred at 25°C for 1 h. The reaction mixture was diluted with water (30 mL) then extracted with ethyl acetate (30 mL * 2). The combined organic layers were washed with brine (60 mL * 1), the collected organics were dried over anhydrous sodium sulfate, filtered and concentrated under reduced pressure to give a residue. The residue was purified by flash silica gel chromatography (ISCO®; 4 g SepaFlash® Silica Flash Column, Eluent of 0∼7% Methanol/Dichloromethane @ 50 mL/min). Tert-butyl (S)-7-hydroxy-3-(methylcarbamoyl)-3,4-dihydroisoquinoline-2(1H)-carboxylate (330 mg, 1.08 mmol, 63.2% yield) was obtained as a white solid. H NMR (400 MHz, DMSO-d_6_): δ = 9.35 - 9.10 (m, 1H), 7.82 - 7.62 (m, 1H), 7.07 - 6.79 (m, 1H), 6.65 - 6.37 (m, 2H), 4.66 - 4.32 (m, 2H), 4.30 - 4.20 (m, 1H), 3.17 (d, J = 4.8 Hz, 1H), 3.00 - 2.74 (m, 2H), 2.52 (s, 2H), 1.47 - 1.34 (m, 9H).

Preparation of (S)-7-hydroxy-N-methyl-1,2,3,4-tetrahydroisoquinoline-3-carboxamide. To a mixture of tert-butyl (S)-7-hydroxy-3-(methylcarbamoyl)-3,4-dihydroisoquinoline-2(1H)-carboxylate (330 mg, 1.08 mmol, 1.00 eq) in dichloromethane (1 mL) was added hydrochloric acid/dioxane (4 M, 2.5 mL, 9.28 eq), then it was stirred at 25°C for 1 h. The reaction mixture was concentrated under reduced pressure to give a residue. (S)-7-hydroxy-N-methyl-1,2,3,4-tetrahydroisoquinoline-3-carboxamide (260 mg, crude, hydrochloric acid) was obtained as a white solid. H NMR (400 MHz, DMSO-d_6_): δ = 9.78 - 9.61 (m, 1H), 9.60 - 9.47 (m, 1H), 9.45 - 9.29 (m, 1H), 8.61 (d, J = 4.3 Hz, 1H), 7.01 (d, J = 8.4 Hz, 1H), 6.70 (d, J = 8.4 Hz, 1H), 6.62 (s, 1H), 4.30 - 4.13 (m, 2H), 4.05 (s, 1H), 3.21 - 3.10 (m, 1H), 2.87 (dd, J = 15.6, 12.4 Hz, 1H), 2.70 (d, J = 4.0 Hz, 3H).

#### Preparation of (S)-2-(5-chloro-3-methylimidazo[1,2-a]pyridine-2-carbonyl)-7-hydroxy-N-methyl-1,2,3,4-tetrahydroisoquinoline-3-carboxamide (8b)

To a mixture of 5-chloro-3-methylimidazo[1,2-a]pyridine-2-carboxylic acid (44 mg, 206 μmol, 1.00 eq) in dichloromethane (1 mL) was added N,N-diisopropylethylamine (107 mg, 824 μmol, 144 μL, 4.00 eq) and 2,4,6-tributyl-1,3,5,2,4,6trioxatriphosphinane 2,4,6-trioxide (178 mg, 247 μmol, 1.20 eq), then it was stirred at 25°C for 10 min, then (S)-7-hydroxy-N-methyl-1,2,3,4-tetrahydroisoquinoline-3-carboxamide (50 mg, 206 μmol, 1.00 eq, hydrochloric acid) was added, it was stirred at 25°C for 0.5 h. The reaction mixture was diluted with water (30 mL) then extracted with ethyl acetate (30 mL * 2). The combined organic layers were washed with brine (60 mL * 1), the collected organics were dried over anhydrous sodium sulfate, filtered and concentrated under reduced pressure to give a residue. The residue was purified by flash silica gel chromatography (ISCO®; 4 g SepaFlash® Silica Flash Column, Eluent of 0∼7% Methanol/Dichloromethane @ 50 mL/min). (S)-2-(5-chloro-3-methylimidazo[1,2-a]pyridine-2-carbonyl)-7-hydroxy-N-methyl-1,2,3,4-tetrahydroisoquinoline-3-carboxamide (17.98 mg, 10.32 μmol, 5.01% yield, 95.725 % purity) was obtained as a off-white solid. ^1^H NMR (400 MHz, DMSO-d_6_): δ = 8.95 (s, 1H), 7.55 (d, J = 8.0 Hz, 2H), 7.28 - 7.18 (m, 1H), 7.05 (d, J = 7.2 Hz, 1H), 6.96 (d, J = 6.4 Hz, 1H), 6.65 - 6.37 (m, 2H), 5.12 - 4.83 (m, 2H), 4.59 - 4.37 (m, 1H), 3.03 (d, J = 4.6 Hz, 2H), 2.86 (s, 3H), 2.57 (s, 3H). LC-MS: R_t_ = 0.955 min, [M+H]^+^ = 399.1. HPLC: 95.725 % purity, R_t_ =1.129 min. SFC: R_t_ =1.648 min.

#### Synthesis of (E)-3-[4-[3-[[[3-(methylcarbamoyl)phenyl]methylamino]methyl]-1H-pyrazolo[3,4-b]pyridin-5-yl]phenyl]prop-2-enoic acid (9): Preparation of O5-tert-butyl O3-ethyl 1-(1-benzyl oxycarbonylpyrrolidin-3-yl)-6,7-dihydro-4H-pyrazolo[4,3-c]pyridine-3,5-dicarboxylate

To a solution of O5-tert-butyl O3-ethyl 1,4,6,7-tetrahydropyrazolo[4,3-c]pyridine-3,5-dicarboxylate (2.00 g, 6.77 mmol, 1.00 eq), benzyl 3-(p-tolylsulfonyloxy)pyrrolidine-1-carboxylate (3.81 g, 10.1 mmol, 1.50 eq) in dimethylformamide (40 mL) was added cesium carbonate (6.62 g, 20.3 mmol, 3.00 eq) and sodium iodide (1.52 g, 10.1 mmol, 1.50 eq). The mixture was stirred at 60°C for 3 h. The reaction mixture was quenched by addition water (400 mL), and then extracted with ethyl acetate (150 mL * 3). The combined organic layers were washed with brine (450 mL * 1), dried over sodium sulfate filtered and concentrated under reduced pressure to give a residue. The residue was purified by flash silica gel chromatography (ISCO®; 80 g SepaFlash® Silica Flash Column, Eluent of 0∼55% Ethyl acetate/Petroleum ethergradient @ 100 mL/min). O5-tert-butyl O3-ethyl 1-(1-benzyloxycarbonylpyrrolidin-3-yl)-6,7-dihydro-4H-pyrazolo [4,3-c]pyridine-3,5-dicarboxylate (1.10 g, 2.21 mmol, 32.6% yield) was obtained as a yellow oil. H NMR (400 MHz, DMSO-d_6_): δ = 1.28 (t, J=7.2 Hz, 3 H) 1.41 (s, 9 H) 2.22 - 2.35 (m, 2 H) 3.25 - 3.89 (m, 8 H) 4.26 (q, J=6.8 Hz, 2 H) 4.48 (s, 2 H) 4.96 - 5.11 (m, 3 H) 7.28 - 7.40 (m, 5 H).

#### Preparation of 1-(1-benzyloxycarbonylpyrrolidin-3-yl)-5-tert-butoxycarbonyl-6,7-dihydro-4H-pyrazolo[4,3-c]pyridine-3-carboxylic acid

To a mixture of O5-tert-butyl O3-ethyl 1-(1-benzyloxycarbonylpyrrolidin-3-yl)-6,7-dihydro-4H-pyrazolo [4,3-c]pyridine-3,5-dicarboxylate (1.05 g, 2.11 mmol, 1.00 eq) in tetrahydrofuran (21 mL) was added lithium monohydroxide (2 M, 3.16 mL, 3.00 eq) and methanol (8.31 g, 259 mmol, 10.5 mL, 123 eq), then it was stirred at 40°C for 1 h. Concentrate and remove methanol under reduced pressure, was added hydrochloric acid (1M) to make pH neutral, then the mixture was quenched by addition water (25 mL), and then extracted with ethyl acetate (20 mL * 3). The combined organic layers were washed with brine (60 mL * 1), dried over sodium sulfate filtered and concentrated under reduced pressure to give a residue. 1-(1-benzyloxycarbonylpyrrolidin-3-yl)-5-tert-butoxycarbony l-6,7-dihydro-4H-pyrazolo[4,3-c]pyridine-3-carboxylic acid (1.05 g, crude) was obtained as a brown oil. ^1^H NMR (400 MHz, DMSO-d_6_): δ = 1.41 (s, 9 H) 2.22 - 2.33 (m, 2 H) 2.66 - 2.74 (m, 2 H) 3.40 - 3.70 (m, 6 H) 4.45 (s, 2 H) 4.89 - 4.99 (m, 1 H) 5.08 (s, 2 H) 7.27 - 7.39 (m, 5 H).

#### Preparation of tert-butyl 1-(1-benzyloxycarbonyl pyrrolidin-3-yl)-3-(methylcarbamoyl)-6,7-dihydro-4H-pyrazolo[4,3-c]pyridine-5-carboxylate

To a mixture of 1-(1-benzyloxycarbonylpyrrolidin-3-yl)-5-tert-butoxycarbonyl-6,7-dihydro-4H-pyrazolo [4,3-c]pyridine-3-carboxylic acid (1.00 g, 2.13 mmol, 1.00 eq), methanamine;hydrochloride (172 mg, 2.55 mmol, 1.20 eq) in dichloromethane (20 mL), was added O-(7-azabenzotriazol-1-yl)-N,N,N,N-tetramethy luroniumhexafluorophosphate (969 mg, 2.55 mmol, 1.20 eq) and N,N-diisopropylethylamine (824 mg, 6.38 mmol, 1.11 mL, 3.00 eq), then it was stirred at 25°C for 1.5 h. The reaction mixture was quenched by addition water (30 mL), and then extracted with dichloromethane (20 mL * 3). The combined organic layers were washed with brine (60 mL * 1), dried over sodium sulfate filtered and concentrated under reduced pressure to give a residue. The residue was purified by flash silica gel chromatography (ISCO®; 20 g SepaFlash® Silica Flash Column, Eluent of 0∼60% Ethyl acetate/Petroleum ethergradient @ 100mL/min). Tert-butyl 1-(1-benzyloxycarbonylpyrrolidin-3-yl)-3-(methylcarbamoyl)-6,7-dihydro-4H-pyrazolo[4,3-c]pyridine-5-carboxylate (750 mg, 1.55 mmol, 72.9% yield) was obtained as a brown oil. H NMR (400 MHz, DMSO-d_6_): δ = 1.41 (s, 9 H) 2.23 - 2.37 (m, 2 H) 2.67 - 2.76 (m, 5 H) 3.43 - 3.62 (m, 4 H) 3.69 - 3.87 (m, 2 H) 4.47 (s, 2 H) 4.96 (d, J=5.6 Hz, 1 H) 5.08 (d, J=8.0 Hz, 2 H) 7.35 (d, J=9.6 Hz, 5 H) 7.84 (s, 1 H).

#### Preparation of tert-butyl 3-(methylcarbamoyl)-1-pyrrolidin-3-yl-6,7-dihydro-4H-pyrazolo[4,3-c]pyridine-5-carboxylate

A mixture of tert-butyl 1-(1-benzyloxycarbonylpyrrolidin-3-yl)-3-(methylcarbamoyl)-6,7-dihydro-4H-pyrazolo[4,3-c]pyridine-5-carboxylate (400 mg, 827 μmol, 1.00 eq) and Palladium hydroxide (162 mg, 231 μmol, 20% purity, 0.280 eq) in tetrahydrofuran (4 mL) was degassed and purged with hydrogen (1.67 mg, 827 μmol, 1.00 eq) for three times, then the resulting mixture was stirred at 25°C under a 15 Psi pressure for 1 h. The mixture was filtered over celite, washed with tetrahydrofuran (5 mL * 2) and concentrated in vacuum to give a residue. Tert-butyl 3-(methylcarbamoyl)-1-pyrrolidin-3-yl-6,7-dihydro-4H-pyrazolo[4,3-c]pyridine-5-carboxylate (280 mg, 801 μmol, 96.8% yield) was obtained as a brown oil. H NMR (400 MHz, DMSO-d_6_): δ = 1.41 (s, 9 H) 1.87 - 2.02 (m, 1 H) 2.06 - 2.18 (m, 1 H) 2.72 (d, J=4.8 Hz, 3 H) 2.98 (s, 2 H) 3.06 - 3.19 (m, 4 H) 3.59 (s, 2 H) 4.09 (s, 1 H) 4.46 (s, 2 H) 4.70 (d, J=3.6 Hz, 1 H) 7.95 (d, J=3.6 Hz, 1 H).

#### Preparation of tert-butyl 1-[1-[6-(2-chlorophenyl) pyridine-2-carbonyl]pyrrolidin-3-yl]-3-(methylcarbamoyl)-6,7-dihydro-4H-pyrazolo[4,3-c]pyridine-5-carboxylate

To a mixture of tert-butyl 3-(methylcarbamoyl)-1-pyrrolidin-3-yl-6,7-dihydro-4H-pyrazolo[4,3-c]pyridine-5-carboxylate (130 mg, 372 μmol, 1.00 eq), 6-(2-chlorophenyl)pyridine-2-carboxylic acid (104 mg, 446 μmol, 1.20 eq) in dichloromethane (3.5 mL) was added O-(7-azabenzotriazol-1-yl)-N,N,N,N-tetramethy luroniumhexafluorophosphate (169 mg, 446 μmol, 1.20 eq) and N,N-diisopropylethylamine (144 mg, 1.12 mmol, 194 μL, 3.00 eq), then it was stirred at 25°C for 1 h. The reaction mixture was quenched by addition water (10 mL), and then extracted with dichloromethane (5 mL * 3). The combined organic layers were washed with brine (15 mL * 1), dried over sodium sulfate filtered and concentrated under reduced pressure to give a residue. Tert-butyl 1-[1-[6-(2-chlorophenyl)pyridine-2-carbonyl]pyrrolidin-3-yl]-3-(methylcarbamoyl)-6,7-dihydro-4H-pyrazolo[4,3-c]pyridine-5-carboxylate (440 mg, crude) was obtained as a brown oil. LC-MS: R_t_ = 0.484 min, [M+H]^+^ = 565.3.

#### Preparation of (E)-3-[4-[3-[[[3-(methylcarbamoyl)phenyl]methylamino]methyl]-1H-pyrazolo[3,4-b]pyridin-5-yl]phenyl]prop-2-enoic acid (9)

To a mixture of tert-butyl 1-[1-[6-(2-chlorophenyl)pyridine-2-carbonyl]pyrrolidin-3-yl]-3-(methy lcarbamoyl)-6,7-dihydro-4H-pyrazolo[4,3-c]pyridine-5-carboxylate (220 mg, 389 μmol, 1.00 eq) in dichloromethane (1.1 mL) was added hydrochloric acid/dioxane (4 M, 1.10 mL, 11.3 eq) at 0°C, then it was stirred at 25°C for 1 h. The mixture was concentrated in vacuum to give a crude product. The residue was purified by prep-HPLC (column: Phenomenex C18 150 * 25 mm * 10 um; mobile phase:[water (NH4HCO3)-CAN]; gradient: 20%-50% B over 10 min). 1-[1-[6-(2-chlorophenyl)pyridine-2-carbonyl] pyrrolidin-3-yl]-N-methyl-4,5,6,7-tetrahydropyrazolo[4,3-c]pyridine-3-carboxamide (15.56 mg, 33.05 μmol, 8.49% yield, 98.765% purity) was obtained as a yellow solid. H NMR (400 MHz, DMSO-d_6_): δ = 2.34 (d, J=6.8 Hz, 2 H) 2.55 (s, 1 H) 2.63 (d, J=4.8 Hz, 1 H) 2.65 - 2.71 (m, 3 H) 2.80 - 2.98 (m, 2 H) 3.67 - 4.24 (m, 6 H) 4.93 - 5.07 (m, 1 H) 7.42 - 7.83 (m, 7 H) 8.06 (q, J=7.6 Hz, 1 H). LC-MS: R_t_ = 1.281 min, [M+H]^+^ = 465.2, method: 5-95N_3min.lcm. HPLC: 98.765% purity, R_t_ =0.950 min.

#### Synthesis of N-methyl-2-(4-(1-((1-((4-oxo-2,3,4,5-tetrahydrobenzo[b][1,4]thiazepin-7-yl)sulfonyl)azetidin-3-yl)methyl)azetidin-3-yl)piperazin-1-yl)acetamide (10a): Preparation of tert-butyl 3-(4-(2-(methylamino)-2-oxoethyl)piperazin-1-yl)azetidine-1-carboxylate

To a solution of tert-butyl 3-(piperazin-1-yl)azetidine-1-carboxylate (1.00 g, 4.14 mmol, 1.00 eq) in dimethyl formamide (12.0 mL) was added N,N-diisopropylethylamine (1.61 g, 12.4 mmol, 2.17 mL, 3.00 eq) and 2-bromo-N-methylacetamide (944 mg, 6.22 mmol, 1.50 eq). The mixture was stirred at 50°C for 2 h. reaction mixture was diluted with water 100 mL and extracted with ethyl acetate 60.0 mL (20.0 mL * 3). The combined organic layers were dried over sodium sulfate, filtered and concentrated under reduced pressure to give a residue. The residue was purified by flash silica gel chromatography (ISCO®; 12 g SepaFlash® Silica Flash Column, Eluent of 0∼5% dichloromethane/methanol @ 100 mL/min). Tert-butyl 3-(4-(2-(methylamino)-2-oxoethyl)piperazin-1-yl)azetidine-1-carboxylate (700 mg, 2.24 mmol, 54.0% yield) was obtained as a yellow oil. H NMR (400 MHz, DMSO-d_6_): δ = 7.62 (d, J = 4.0 Hz, 1H), 3.89 - 3.59 (m, 4H), 3.31 (d, J = 9.6 Hz, 3H), 3.07 - 2.81 (m, 4H), 2.61 - 2.56 (m, 3H), 2.33 (d, J = 12.8 Hz, 4H), 1.36 (d, J = 9.2 Hz, 9H).

#### Preparation of 2-(4-(azetidin-3-yl)piperazin-1-yl)-N-methylacetamide

To a solution of tert-butyl 3-(4-(2-(methylamino)-2-oxoethyl)piperazin-1-yl)azetidine-1-carboxylate (1.30 g, 4.16 mmol, 1.00 eq) in dichloromethane (13.0 mL) was added trifluoroacetic acid (3.99 g, 35.0 mmol, 2.60 mL, 8.41 eq). The mixture was stirred at 25°C for 2 h. The reaction mixture was concentrated under reduced pressure to give a residue and then adjust PH>7 by sodium carbonate. 2-(4-(azetidin-3-yl)piperazin-1-yl)-N-methylacetamide (2.50 g, crude) was obtained as a yellow oil. H NMR (400 MHz, DMSO-d_6_): δ = 7.69 (s, 1H), 3.76 - 3.62 (m, 5H), 3.20 (d, J = 6.8 Hz, 1H), 2.66 (d, J = 4.4 Hz, 3H), 2.51 - 2.32 (m, 8H), 1.30 (s, 1H).

#### Preparation of tert-butyl 3-((3-(4-(2-(methylamino)-2-oxoethyl)piperazin-1-yl)azetidin-1-yl)methyl)azetidine-1-carboxylate

To a solution of 2-(4-(azetidin-3-yl)piperazin-1-yl)-N-methylacetamide (700 mg, 1.65 mmol, 1.00 eq) and tert-butyl 3-formylazetidine-1-carboxylate (305 mg, 1.65 mmol, 1.00 eq) in tetrahydrofuran (7.0 mL) was added sodium triacetoxyhydroborate (698 mg, 3.30 mmol, 2.00 eq) and acetic acid (99.0 mg, 1.65 mmol, 94.3 μL, 1.00 eq). The mixture was stirred at 30°C for 1 h. The reaction mixture was concentrated under reduced pressure to give a residue. The residue was purified by flash silica gel chromatography (ISCO®; 12 g SepaFlash® Silica Flash Column, Eluent of 0∼100% Ethyl acetate/Petroleum ethergradient @ 60 mL/min). Tert-butyl 3-((3-(4-(2-(methylamino)-2-oxoethyl)piperazin-1-yl)azetidin-1-yl)methyl) azetidine-1-carboxylate (425 mg, 891 μmol, 54.0% yield, 80% purity) was obtained as a yellow oil. H NMR (400 MHz, DMSO-d_6_): δ = 7.69 (s, 1H), 3.76 - 3.62 (m, 5H), 3.20 (d, J = 6.8 Hz, 1H), 2.66 (d, J = 4.4 Hz, 3H), 2.51 - 2.32 (m, 8H), 1.30 (s, 1H)

#### Preparation of 2-(4-(1-(azetidin-3-ylmethyl) azetidin-3-yl)piperazin-1-yl)-N-methylacetamide

To a solution of tert-butyl 3-((3-(4-(2-(methylamino)-2-oxoethyl)piperazin-1-yl)azetidin-1-yl)methyl) azetidine-1-carboxylate (200 mg, 419 μmol, 1.00 eq) in dichloromethane (2.0 mL) was added trifluoroacetic acid (245 mg, 2.15 mmol, 160 μL, 5.14 eq). The mixture was stirred at 25°C for 1 h. The reaction mixture was concentrated under reduced pressure to give a residue. The crude product 2-[4-[1-(azetidin-3-ylmethyl)azetidin-3-yl]piperazin-1-yl]-N-methyl-acetamide (297 mg, crude, trifluoroacetic acid) was used into the next step without further purification. 2-(4-(1-(azetidin-3-ylmethyl) azetidin-3-yl)piperazin-1-yl)-N-methylacetamide (297 mg, crude, trifluoroacetic acid) was obtained as a yellow oil. H NMR (400 MHz, DMSO-d_6_): δ = 8.98 (d, J = 5.6 Hz, 1H), 8.85 (d, J = 4.0 Hz, 1H), 4.29 - 4.03 (m, 2H), 4.02 - 3.63 (m, 8H), 3.62 - 3.25 (m, 5H), 3.22 - 2.78 (m, 4H), 2.67 (d, J = 4.8 Hz, 3H), 2.51 (s, 3H).

#### Preparation of N-methyl-2-(4-(1-((1-((4-oxo-2,3,4,5-tetrahydrobenzo[b][1,4]thiazepin-7-yl)sulfonyl)azetidin-3-yl)methyl)azetidin-3-yl)piperazin-1-yl)acetamide (10a)

A mixture of 2-(4-(1-(azetidin-3-ylmethyl)azetidin-3-yl)piperazin-1-yl)-N-methylacetamide (80.0 mg, 111 μmol, 2.00 eq, trifluoroacetic acid) and potassium carbonate (23.0 mg, 166 μmol, 3.00 eq) in tetrahydrofuran (1.0 mL) was degassed and purged with nitrogen for 3 times, and then the 4-oxo-2,3,4,5-tetrahydrobenzo[b][1,4]thiazepine-7-sulfonyl chloride (15.4 mg, 55.6 μmol, 1.00 eq) in tetrahydrofuran (0.1 mL) was added in the mixture at 0°C, the mixture was stirred at 0°C for 0.5 h under nitrogen atmosphere. The reaction mixture was concentrated under reduced pressure to give a residue. The residue was purified by prep-HPLC (column: Phenomenex C18 150*25mm*10um;mobile phase: [water(ammonium bicarbonate)-acetonitrile];gradient:10%-40% B over 10 min). N-methyl-2-(4-(1-((1-((4-oxo-2,3,4,5-tetrahydrobenzo[b][1,4]thiazepin-7-yl)sulfonyl)azetidin-3-yl)methyl)azetidin-3-yl)piperazin-1-yl)acetamide (15.6 mg, 29.3 μmol, 52.7% yield, 98% purity) was obtained as white solid. H NMR (400 MHz, DMSO-d_6_): δ = 7.82 (d, J = 8.0 Hz, 1H), 7.58 (d, J = 4.4 Hz, 1H), 7.51 (dd, J = 1.6, 8.0 Hz, 1H), 7.45 (s, 1H), 3.76 (t, J = 8.4 Hz, 2H), 3.49 (t, J = 6.8 Hz, 2H), 3.34 (s, 2H), 3.17 (t, J = 6.0 Hz, 2H), 2.85 (s, 2H), 2.71 (t, J = 6.4 Hz, 1H), 2.63 - 2.53 (m, 7H), 2.37 (d, J = 5.6 Hz, 5H), 2.18 (d, J = 6.8 Hz, 6H). LC-MS: R_t_ = 1.361 min, [M+H]^+^ = 523.2. HPLC: 98% purity, R_t_ = 1.335 min.

#### Synthesis of 2-(4-(1-((1-(benzo[c][1,2,5]thiadiazol-5-ylsulfonyl)azetidin-3-yl)methyl)azetidin-3-yl)piperazin-1-yl)-N-methylacetamide (10b): Preparation of3-(dimethoxymethyl)-1-((4-fluoro-3-nitrophenyl)sulfonyl)azetidine

To a solution of 3-(dimethoxymethyl)azetidine (821 mg, 6.26 mmol, 1.50 eq) in dichloromethane (10 mL) was added triethylamine (422 mg, 4.17 mmol, 581 μL, 1.00 eq) and a solution of 4-fluoro-3-nitrobenzenesulfonyl chloride (1.00 g, 4.17 mmol, 1.00 eq) in dichloromethane (10 mL) at 0°C. The mixture was stirred at 0°C for 1 h. The mixture was concentrated in vacuum to give a residue. The residue was purified by flash silica gel chromatography (ISCO®; 20 g SepaFlash® Silica Flash Column, Eluent of 0∼30% Ethyl acetate/Petroleum ethergradient @ 50 mL/min). 3-(dimethoxymethyl)-1-(4-fluoro-3-nitro-phenyl)sulfonyl-azetidine (350 mg, 1.04 mmol, 24.8% yield, 99.0% purity) was obtained as a yellow solid. H NMR (400 MHz, DMSO-d_6_): δ = 8.43 (dd, J = 2.4, 6.8 Hz, 1H), 8.27 - 8.20 (m, 1H), 7.92 (dd, J = 8.8, 10.8 Hz, 1H), 4.22 (d, J = 5.6 Hz, 1H), 3.82 (t, J = 8.8 Hz, 2H), 3.58 (dd, J = 6.8, 8.4 Hz, 2H), 3.10 (s, 6H), 2.78 - 2.68 (m, 1H).

#### Preparation of compound 5-4-((3-(dimethoxymethyl)azetidin-1-yl)sulfonyl)-2-nitroaniline

To a solution of 3-(dimethoxymethyl)-1-(4-fluoro-3-nitro-phenyl)sulfonyl-azetidine (300 mg, 897 μmol, 1.00 eq) in acetonitrile (7.5 mL) was added ammonium carbonate (862 mg, 8.97 mmol, 958 μL, 10.0 eq). The mixture was stirred at 60°C for 2 h. The mixture was poured into water (50 mL) and extracted with dichloromethane (50 mL*3). The combined organic layers were dried over sodium sulfate, filtered and concentrated under reduced pressure to give a crude product. 4-[3-(dimethoxymethyl)azetidin-1-yl] sulfonyl-2-nitro-aniline (310 mg, crude) was obtained as a yellow oil. H NMR (400 MHz, DMSO-d_6_): δ = 8.30 (s, 1H), 8.11 (s, 2H), 7.73 (d, J = 8.8 Hz, 1H), 7.21 (d, J = 9.2 Hz, 1H), 4.21 (d, J = 5.6 Hz, 1H), 3.73 (t, J = 8.4 Hz, 2H), 3.49 (t, J = 7.2 Hz, 2H), 3.12 (s, 6H), 2.72 (dd, J = 6.8, 14.0 Hz, 1H).

#### Preparation of 4-((3-(dimethoxymethyl)azetidin-1-yl)sulfonyl)benzene-1,2-diamine

To a solution of 4-[3-(dimethoxymethyl)azetidin-1-yl]sulfonyl-2-nitro-aniline (477 mg, 1.44 mmol, 1.00 eq) in ethanol (15 mL) and water (2.0 mL) was added iron powder (803 mg, 14.4 mmol, 10.0 eq) and ammonium chloride (770 mg, 14.4 mmol, 10.0 eq). The mixture was stirred for 1.5 h at 80°C. The mixture was filtered and filtrate was concentrated in vacuum to give a crude product. 4-[3-(dimethoxymethyl)azetidin-1-yl]sulfonylbenzene-1,2-diamine (420 mg, crude) was obtained as a brown solid. LCMS: R_t_ = 0.224 min, [M+H]^+^ = 302.1.

Preparation of 5-((3-(dimethoxymethyl)azetidin-1-yl)sulfonyl)benzo[c][1,2,5]thiadiazole To a solution of 4-[3-(dimethoxymethyl)azetidin-1-yl]sulfonylbenzene-1,2-diamine (50 mg, 166 μmol, 1.00 eq), triethylamine (252 mg, 2.49 mmol, 346 μL, 15.0 eq) in THF (1.0 mL) was drop-wise added a solution of sulfurous dichloride (99 mg, 829 μmol, 60 μL, 5.00 eq) in THF (1.0 mL) at 0°C. The mixture was stirred at 35°C for 12 h. The mixture was poured into water (5 mL) and extracted with DCM (5 mL*3). The organic phase was separated, dried over sodium sulfate, filtered and concentrated under reduced pressure to give a crude product. 5-[3-(dimethoxymethyl)azetidin-1-yl]sulfonyl-2,1,3-benzothiadiazole (450 mg, crude) was obtained as a black solid. LCMS: R_t_ = 0.313 min, [M+H]^+^ = 330.1.

#### Preparation of 1-(benzo[c][1,2,5]thiadiazol-5-ylsulfonyl)azetidine-3-carbaldehyde

To a solution of 5-[3-(dimethoxymethyl)azetidin-1-yl]sulfonyl-2,1,3-benzothiadiazole (200 mg, 607 μmol, 1.00 eq) in dichloromethane (2.0 mL) was added trifluoroacetic acid (614 mg, 5.38 mmol, 0.4 mL, 8.87 eq). The mixture was stirred at 20°C for 2 h. The mixture was concentrated poured into sodium bicarbonate.aq (30 mL) and extracted with dichloromethane (30 mL*4). The combined organic layers were dried over sodium sulfate, filtered and concentrated under reduced pressure to give crude product. 1-(2,1,3-benzothiadiazol-5-ylsulfonyl)azetidine-3-carbaldehyde (140 mg, crude) was obtained as a black oil. LC-MS: [M+H]^+^ = 284.0.

#### Preparation of 2-(4-(1-((1-(benzo[c][1,2,5]thiadiazol-5-ylsulfonyl)azetidin-3-yl)methyl)azetidin-3-yl)piperazin-1-yl)-N-methylacetamide (10b)

To a solution of 1-(2,1,3-benzothiadiazol-5-ylsulfonyl)azetidine-3-carbaldehyde (120 mg, 423 μmol, 1.00 eq), 2-[4-(azetidin-3-yl)piperazin-1-yl]-N-methyl-acetamide (179 mg, 847 μmol, 2.00 eq) in tetrahydrofuran (2.5 mL) was added sodium acetate (104 mg, 1.27 mmol, 3.00 eq) and stirred for 2 h at 35°C, then sodium triacetoxyhydroborate (179 mg, 847 μmol, 2.00 eq) was added at 0°C. The mixture was stirred at 20°C for 1 h. The mixture was poured into saturated sodium bicarbonate.aq and extracted with ethyl acetate (30 mL*3). The combined organic layers were washed dried over sodium sulfate, filtered and concentrated under reduced pressure to give a residue. The residue was purified by Prep-HPLC (column: Waters xbridge 150*25mm 10um;mobile phase: [water(NH4HCO3)-ACN];gradient:10%-40% B over 14 min).2-[4-[1-[[1-(2,1,3-benzothiadiazol-5-ylsulfonyl)azetidin-3-yl]methyl]azetidin-3-yl]piperazin-1-yl]-N-methyl-acetamide (7.94 mg, 15.40 μmol, 3.64% yield, 93.05% purity) was obtained as a white solid. H NMR (400 MHz, DMSO-d_6_):δ = 8.62 - 8.52 (m, 1H), 8.43 - 8.31 (m, 1H), 8.02 (d, J = 9.0 Hz, 1H), 7.63 - 7.52 (m, 1H), 3.84 (t, J = 8.0 Hz, 2H), 3.47 (t, J = 7.2 Hz, 2H), 3.07 (t, J = 6.0 Hz, 2H), 2.84 (s, 2H), 2.61 - 2.56 (m, 3H), 2.54 (d, J = 6.4 Hz, 2H), 2.49 - 2.47 (m, 2H), 2.40 (d, J = 7.2 Hz, 2H), 2.33 (s, 3H), 2.22 (d, J = 6.8 Hz, 2H), 2.11 (s, 3H). LCMS: R_t_ = 1.100 min, [M+H]^+^ = 480.2. HPLC: 93.05% purity, R = 1.441 min.

