## Supplemental File for "Development of a DNA-encoded library screening method “DEL Zipper” to empower the study of RNA-targeted chemical matter"

**Supplementary figures**

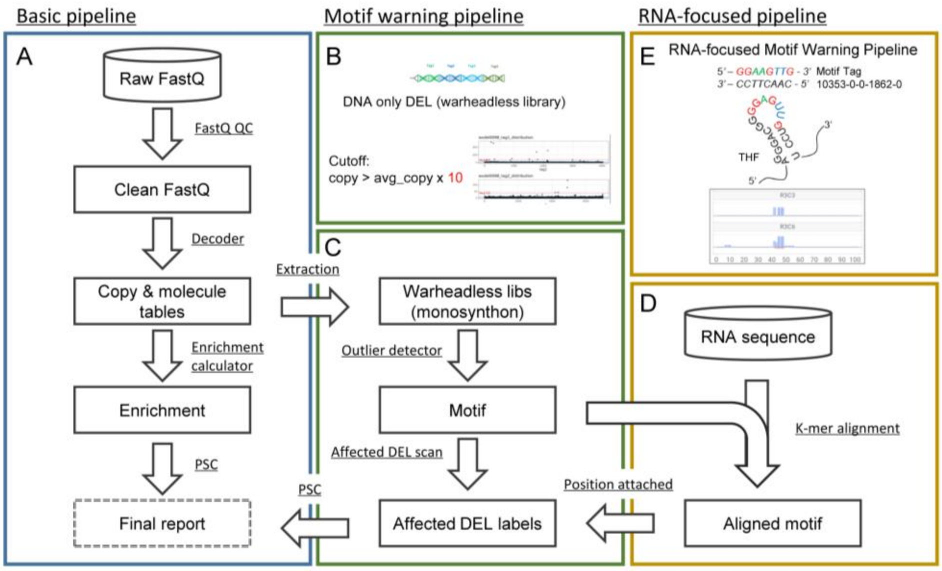


**Supplementary Figure 1. Schematic Flowchart of RNA Specialized Building Block Motif Warning Pipeline used in this study.** Flowchart summarizing the major steps of the RNA-focused data-processing pipeline. (A) is the basic pipeline from raw FastQ to final report. (B) shows the building block (BB) tag structure of only DEL and the example of motif warning results. (C) is the BB motif warning pipeline using warhead-less libraries (D) is the RNA-focused pipeline to find & validate RNA sequence related BB motifs using K-mer alignment and (E) is the example of RNA sequence related BB motif detected in THF riboswitch DEL selection using RNA specialized motif warning pipeline. The Detected Motif is *AGGAAGTTGGA*.

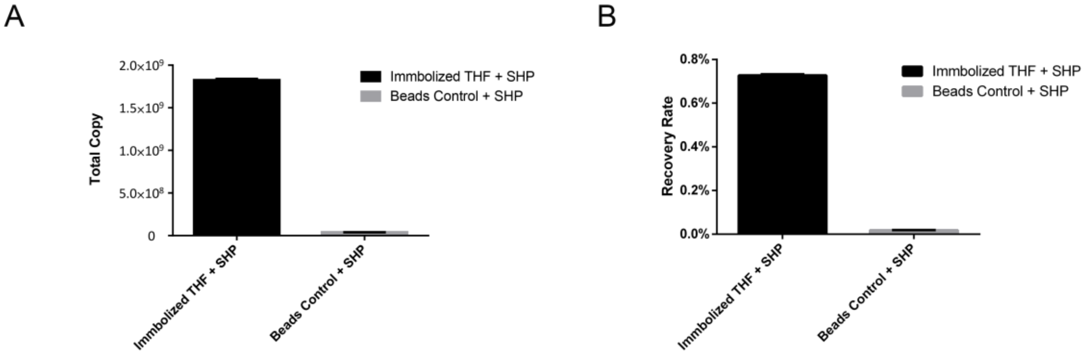


**Supplementary Figure 2. Pre-selection results of single-stranded DEL tag Spacer-headpiece (SHP) negative control against THF riboswitch RNA.** In pre-selection experiments, single-stranded DEL tag SHP negative controls show more (A) after selection Total Copy and (B) after selection recovery rate in immobilized THF group comparing to beads control group. These results indicate the potential RNA derived RNA-DEL specific binding noise. Data are the mean ± S.D. of 3 replicates (n = 3).

| No. | Sample ID |
|---|---|
| M | Marker |
| 1 | Library Before Closing Tag Ligation |
| 2 | Library After Closing Tag Ligation |
| 3 | DEL Zipper 1 |
| 4 | DEL Zipper 2 |
| 5 | DEL Zipper 3 |
| 6 | DEL Zipper 4 |
| 7 | DEL Zipper 5 |
| 8 | DEL Zipper 6 |
| 9 | DEL Zipper 7 |
| 10 | DEL Zipper 8 |

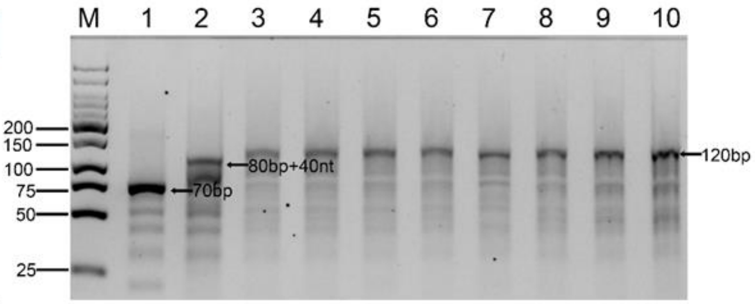


**Supplementary Figure 3. Gel Image of "DEL Zipper" library before and after conversion.** Gel electrophoresis shows that a clear size difference of DEL Library can be observed after closing tag ligation (Lane 1 vs lane 2) and after "DEL Zipper" conversion (Lane 2 vs Lane 3 to Lane 10).

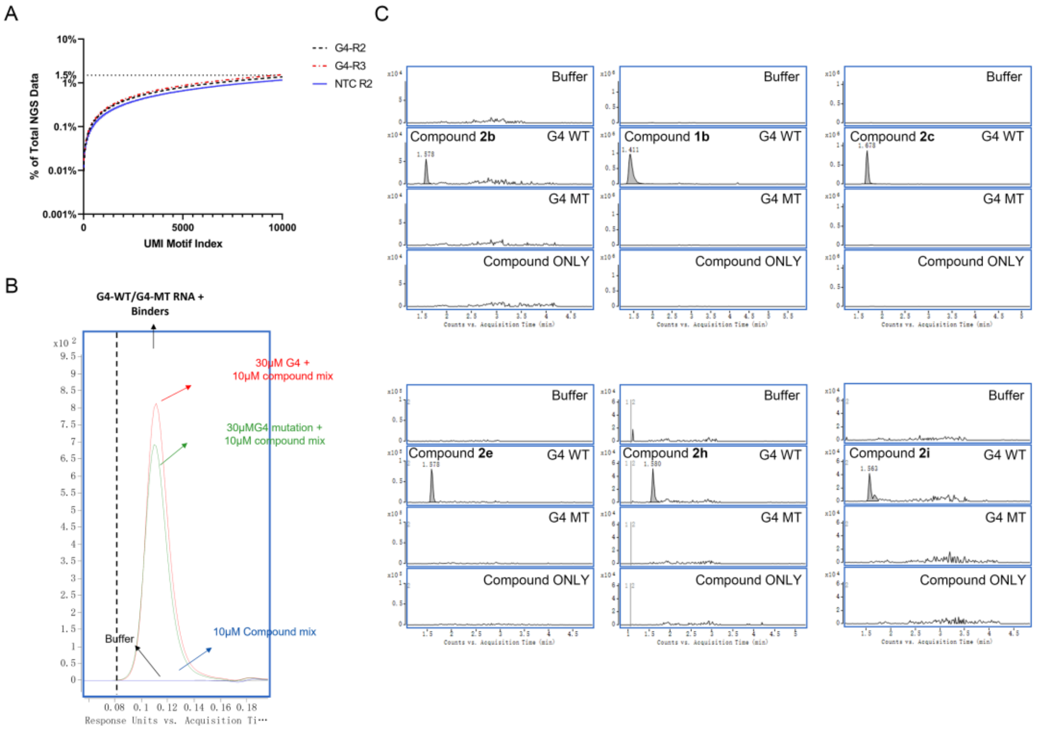


**Supplementary Figure 4. DNA-RNA binding events in G4-RNA round 2 & round 3 selection and pre-selection DEL selected G4 compounds shows selectivity signals in ASMS. (A)** Copy accumulation curve of potential UMI motif and overall proportion of total NGS data. In G4-RNA Round 2(R2) & round 3 (R3) selection, the potential UMI motif consist of less than 1.5% of total NGS data and shows similar pattern to No Target Control. **(B to C)** In ASMS experiments, G4 wild type (WT,30 μM) or G4 mutant (G4 MT, 30 μM) are incubated with DEL selected compounds (10 μM) and further analyzed by two-dimensional ASMS. (B) G4 WT and G4 MT can both be detected in SEC-UV with relative similar signal response. (C) LC-MS results of buffer, G4-WT, G4-MT and compound only. Only in G4 WT condition, corresponding DEL selected compound can be detected. No compound can be detected in Buffer, G4 MT or Compound only condition. 6 compounds are ASMS results shown as example, all compounds detailed MS signal intensity are described in supplementary Table 2, supplementary Table 3 and supplementary Table 4.

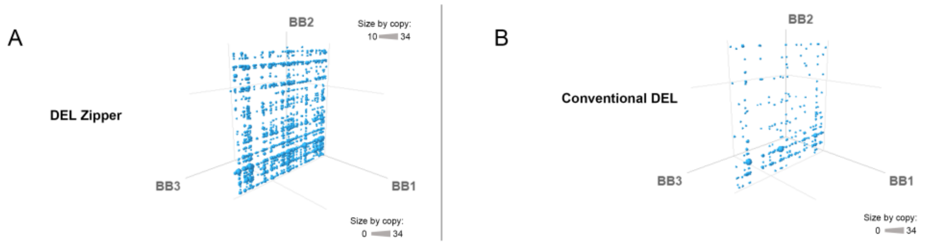


**Supplementary Fig.5 Cubic plots of comparison between DEL zipper library and conventional DEL library side by side in the selection against G4.** Dots in all cube are filtered out the signals below 10 folds enriched than average. (A) Compounds in DEL Zipper library 01 and (B) conventional DEL library 01 against G4.

**Supplementary Table 1.** Detected Coding Region Motif in THF riboswitch DEL selection

| Tag_ID | Tag_Seq | Alignment Position In THF | Affected Mono-synthon count | Relative Copy After NGS | | | | | |
|---|---|---|---|---|---|---|---|---|---|
| | | | | A | B | C | D | E | F |
| 10341-0-224-0-0 | ATACTCCTCAG | 49-55 | 24 | 1.83 | 2.64 | 0.79 | 1.85 | 2.99 | 11.93 |
| 10341-0-5947-0-0 | GAGTTGAAGGT | 43-48 | 20 | 0.87 | 2.12 | 0.59 | 0.75 | 1.17 | 7.23 |
| 10348-0-5947-0-0 | GAGTTGAAGGT | 43-48 | 20 | 1.76 | 3.75 | 0.98 | 0.75 | 3.04 | 15.70 |
| 10353-0-0-1387-0 | AGAGAGAGAGA | \ | 20 | 2.07 | 3.30 | 0.71 | 0.00 | 3.50 | 21.59 |
| 10353-0-0-1862-0 | AGGAAGTTGGA | 41-48 | 15 | 1.48 | 4.83 | 1.71 | 0.78 | 1.22 | 15.41 |
| 10353-0-0-1902-0 | AGGAGTAAGGA | 6-10 | 15 | 32.25 | 6.62 | 0.00 | 0.00 | 65.59 | 47.88 |
| 10353-0-0-0-866 | ATCAACTCCTT | 48-42 | 16 | 4.67 | 3.00 | 0.43 | 0.00 | 8.46 | 16.90 |

A: Conventional DEL selection on THF riboswitch, Round 2
B: “DEL Zipper” selection on THF riboswitch, Round 2
C: Conventional DEL selection on No Target Control, Round 2
D: “DEL Zipper” selection on No Target Control, Round 2
E: Conventional DEL selection on THF riboswitch, Round 3
F: “DEL Zipper” selection on THF riboswitch, Round 3

Values over 10 relative copy thresholds are Red labeled and are identified as potential dsDNA-ssRNA interaction motif in our motif pipeline. No Motif is identified in Round2 selection of Zipper-DEL selection against THF-riboswitch RNA. Only 6 tag sequence are identified as potential motif after three rounds of Zipper-DEL selection, which mostly are below 20 relative copies.

**Supplementary Table 2.** Off-DNA resynthesized compounds of G4.

| | 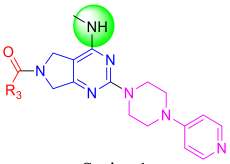 Series 1 | | 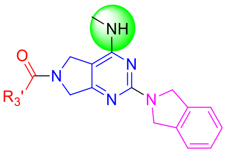 Series 2 | | 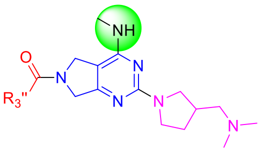 Series 3 |
|---|---|---|---|---|---|
| **Cmpd ID** | R3 | **Cmpd ID** | R3' | **Cmpd ID** | R3'' |
| **1a** | 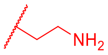 | **2a** | 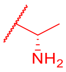 | **3a** | 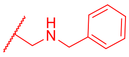 |
| **1b** | 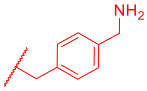 | **2b** | 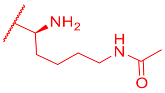 | **3b** | 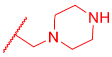 |
| **1c** | 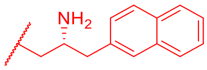 | **2c** | 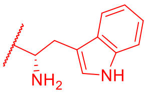 | | |
| **1d** | 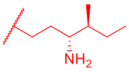 | **2d** | 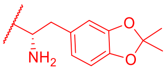 | | |
| **1e** | 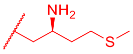 | **2e** | 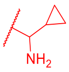 | | |
| **1f** | 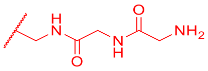 | **2f** | 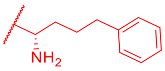 | | |
| **1g** | 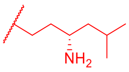 | **2g** | 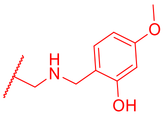 | | |
| **1h** | 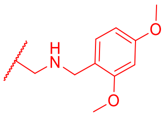 | **2h** | 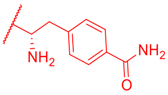 | | |
| **1i** | 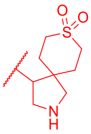 | **2i** | 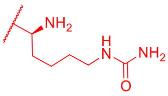 | | |

**Supplementary Table 3.** ASMS signal intensity of DEL-Selected Off-DNA compound against G4.

| Target Name | Product ID | formula | MW | RT | m/z | MW | Score | Error | Signal Intensity |
|---|---|---|---|---|---|---|---|---|---|
| G4_WT | **1a** | C19H26N8O | 382.472 | 1.344 | 383.2296 | 382.222 | 62.1 | -2.48 | 29573 |
| G4_WT | **1b** | C25H30N8O | 458.57 | 1.411 | 459.2613 | 458.254 | 99.61 | -0.55 | 990989 |
| G4_WT | **1c** | C30H34N8O | 522.657 | 1.594 | 523.2934 | 522.2853 | 90.69 | -0.43 | 16069 |
| G4_WT | **1d** | C24H36N8O | 452.607 | 1.511 | 453.3081 | 452.3013 | 75.84 | 0.29 | 14259 |
| G4_WT | **1e** | C22H32N8OS | 456.613 | 1.461 | 495.2061 | 456.2406 | 55.43 | -3.05 | 29207 |
| G4_WT | **1f** | C22H30N10O3 | 482.549 | 1.396 | 483.2569 | 482.2494 | 95.77 | -1.65 | 41925 |
| G4_WT | **1g** | C24H36N8O | 452.607 | 1.597 | 453.3081 | 452.3015 | 77.42 | 0.75 | 11476 |
| G4_WT | **1h** | C27H34N8O3 | 528.622 | 1.613 | 519.2814 | 518.2747 | 90.05 | -1.23 | 27127 |
| G4_WT | **1i** | C25H34N8O3S | 526.66 | 1.463 | 527.2543 | 526.2496 | 79.27 | 4.11 | 20637 |
| G4_WT | **2a** | C18H22N6O | 338.415 | 1.527 | 339.193 | 338.1857 | 99.49 | 0.7 | 282961 |
| G4_WT | **2b** | C23H31N7O2 | 437.548 | 1.578 | 438.2607 | 437.2533 | 84.95 | -1.32 | 69091 |
| G4_WT | **2c** | C26H27N7O | 453.55 | 1.678 | 454.2348 | 453.2275 | 98.97 | -0.41 | 1027855 |
| G4_WT | **2d** | C27H30N6O3 | 486.576 | 1.744 | 487.2444 | 486.2367 | 89.26 | -2.45 | 30565 |
| G4_WT | **2e** | C20H24N6O | 364.453 | 1.578 | 365.208 | 364.2007 | 97 | -1.34 | 90396 |
| G4_WT | **2f** | C26H30N6O | 442.567 | | | | | | |
| G4_WT | **2g** | C25H28N6O3 | 460.538 | 1.68 | 461.2294 | 460.2233 | 63.2 | 2.23 | 14489 |
| G4_WT | **2h** | C25H27N7O2 | 457.538 | 1.58 | 496.1839 | 457.2219 | 67.34 | -1.53 | 58428 |
| G4_WT | **2i** | C22H30N8O2 | 438.536 | 1.563 | 477.2121 | 438.2484 | 83.04 | -1.86 | 55849 |
| G4_WT | **3a** | C23H33N7O | 423.565 | | | | | | |
| G4_WT | **3b** | C20H34N8O | 402.547 | | | | | | |
